# Molecular analysis of binding region of an ACE2 as a receptor for SARS-CoV-2 between humans and mammals

**DOI:** 10.1101/2020.07.09.196378

**Authors:** Takuma Hayashi, Kaoru Aboko, Masaki Mandan, Nobuo Yaegashi, Ikuo Konishi

## Abstract

In June 2020, a second wave of coronavirus disease-2019 (COVID-19) infections raised concern in Beijing, where salmon sold a fresh fish wholesale market was suspected of being the source of severe acute respiratory syndrome coronavirus 2 (SARS-CoV-2) infections. It has raised questions in the press and elsewhere about the scientific basis of salmon as a source of infection. With the number of cases growing, the surface of a salmon chopping board in the market was examined for the presence of SARS-CoV-2 and a positive reaction was observed. Following these test results, there has been debate over whether salmon can be infected with SARS-CoV-2. To find assess this, we investigated the structural homology of angiotensin-converting enzyme 2 (ACE2), a host-side receptor for SARS-CoV-2, between humans and other species including salmon and mink. As a result, a high structural homology between ACE2 and mink, which has reportedly transmitted SARS-CoV-2 to humans, was confirmed. However, a non-high structural homology of ACE2 between salmon and humans was observed. Further experiments are needed to find the source of SARS-CoV-2 transmission to the salmon.

## Introduction

After approximately two months during which no new cases of severe acute respiratory syndrome coronavirus-type 2 (SARS-CoV-2) had been observed in Beijing, a new outbreak in the city was identified in mid-June 2020. With more than 100 coronavirus disease 2019 (COVID-19) infections reported in the days following June 12, Beijing was forced to raise its emergency response level (1,2). The source of the infections is believed to have originated at a fresh fish market in Beijing.

Chairman Zhang Yuan, of the Beijing Fresh Fish Wholesale Market, reported that a sampling survey conducted by China’s National Institute of Health had detected the presence of SARS-CoV-2 on a chopping board used for cutting up imported salmon sourced from a deep seafood market in Beijing. On June 13, a spokesman for the Beijing Health Commission reported that 40 samples had been confirmed positive for SARS-CoV-2 at the Beijing Fresh Fish Wholesale Market. At the same time, as the Chinese public became aware of the new cases, salmon was quickly removed from sale at supermarkets and in Japanese restaurants in the city.

At this time, there is no scientific evidence that salmon can be infected with SARS-CoV-2 or carry the pathogen; therefore, the likelihood of transmission to humans from fish contaminated with SARS-CoV-2 is extremely low. Instead, it is highly possible that the surface of the fish or the cutting board had been contaminated.

In April 2020, 16 international organizations, including the Food and Agriculture Organization of the United Nations and the Chinese Fisheries Science Research Institute, jointly published a paper in *Asian Fisheries Science* to assert that no evidence had indicated that marine products, including fish, crustaceans, mollusks, and amphibians, could be infected with SARS-CoV-2.

It is known that the five genera of coronaviruses only infect birds and mammals, while the beta coronaviruses to which SARS-CoV-2 belongs are believed to infect mammals alone. Therefore, marine products are unlikely to be the source of the new infections.

On June 10, 2020, mink infected with SARS-CoV-2 were found in more than 10 farms in the Netherlands (3). The following month, the Dutch Ministry of Agriculture reported the transmission of SARS-CoV-2 from mink at one of these farms, to become the first recorded case of an animal infecting a human (3).

In this COVID-19 pandemic, SARS-CoV-2 infections of species other than mammals and birds have not been observed. A common amino acid locus with angiotensin-converting enzyme 2 (ACE2), a host-side receptor for mammalian SARS-CoV-2, has been found in thermophilic animals (4,5,6). Therefore, we investigated the molecular structural homology of ACE2 between humans and animals including mink and salmon. As a result, a high structural homology of ACE2 was observed between mink and humans. However, a non-high structural homology of ACE2 was observed between salmon and humans. From these results, the infection of salmon with SARS-CoV-2 has not been demonstrated. Further experiments are needed to examine the source of the new Beijing infections.

## METHODS

### Phylogenetic Analysis and Annotation

Reference genomes and amino acids of human, dog, cat, tiger, mink, salmon, bat, pangolin, and snake ACE2 were obtained from the National Center for Biotechnology Information (NCBI) orthologs at the National Library of Medicine. Amino acid homological analysis was performed using Align Sequences Protein BLAST (algorithm protein–protein BLAST) with the protein accession numbers of ACE2s listed in the NCBI Reference Sequence Database in order to determine the whole amino acid homology of ACE2 between humans and other animals. Phylogenetic analyses of the complete protein and major coding regions were performed with RAxML software (version 8.2.9) with 1000 bootstrap replicates using the general time-reversible nucleotide substitution model. Details of the protein accession numbers of ACE2s are available in the supplementary materials.

### Amino Acid Homology Analysis of the Binding Region of ACE2 for Interaction with SARS-CoV-2 Spike Glycoprotein Between Humans and Other Animals

The binding region for interaction with the SARS-CoV-2 spike glycoprotein (82.aa**-MYP**-84.aa, 353.**aa-KGDFR**-357.aa) of the verified genome amino acid sequences of human ACE2 was predicted using the NCBI protein database and Geneious software (version 11.1.5; Auckland, New Zealand), and was annotated using the NCBI Conserved Domain Database. Amino acid homological analysis was performed using Align Sequences Protein BLAST (algorithm protein–protein BLAST) with the protein accession numbers of human and individual animal ACE2s listed in the NCBI Reference Sequence Database. Details of the protein accession numbers of ACE2s are available in the supplementary materials.

### Analysis of the Three-Dimensional Structure of the Binding Site Between Mink, Salmon, and Human ACE2

Spanner is a structural homology modeling pipeline that threads a query amino-acid sequence onto a template protein structure. This is unique in that it handles gaps by spanning the region of interest using fragments of known structures. To create a model, we must provide a template structure, as well as an alignment of the query sequence to be modeled onto the template sequence. Spanner then replaces mismatched residues and fills in any gaps caused by insertions or deletions. Details of the analysis methods are available in the supplementary materials.

## RESULTS

The outbreak of COVID-19 caused by the SARS-CoV-2 virus has become a pandemic, but there is still little understanding about the zoonotic transmission and the antigenicity of the virus. We therefore examined the homology of the whole ACE2 molecule, which is reported to be a cellular receptor for the spike glycoprotein on the virion surface of SARS-CoV-2, between humans, dogs, cats, tigers, mink, salmon, and other mammals. Our findings show that the ACE2 molecule demonstrated 79%–92% homology between humans and dogs, 91%–92% homology between humans and cats, 92% homology between humans and tigers, 93% homology between humans and minks, and 74% homology between humans and salmon (Figure 1, Supplementary Materials). The ACE2 molecule showed 80%–89% homology between humans and bats, 91% homology between humans and pangolins, and 74%–75% homology between humans and snakes (Figure 1, Supplementary Materials).

**Figure 1.**
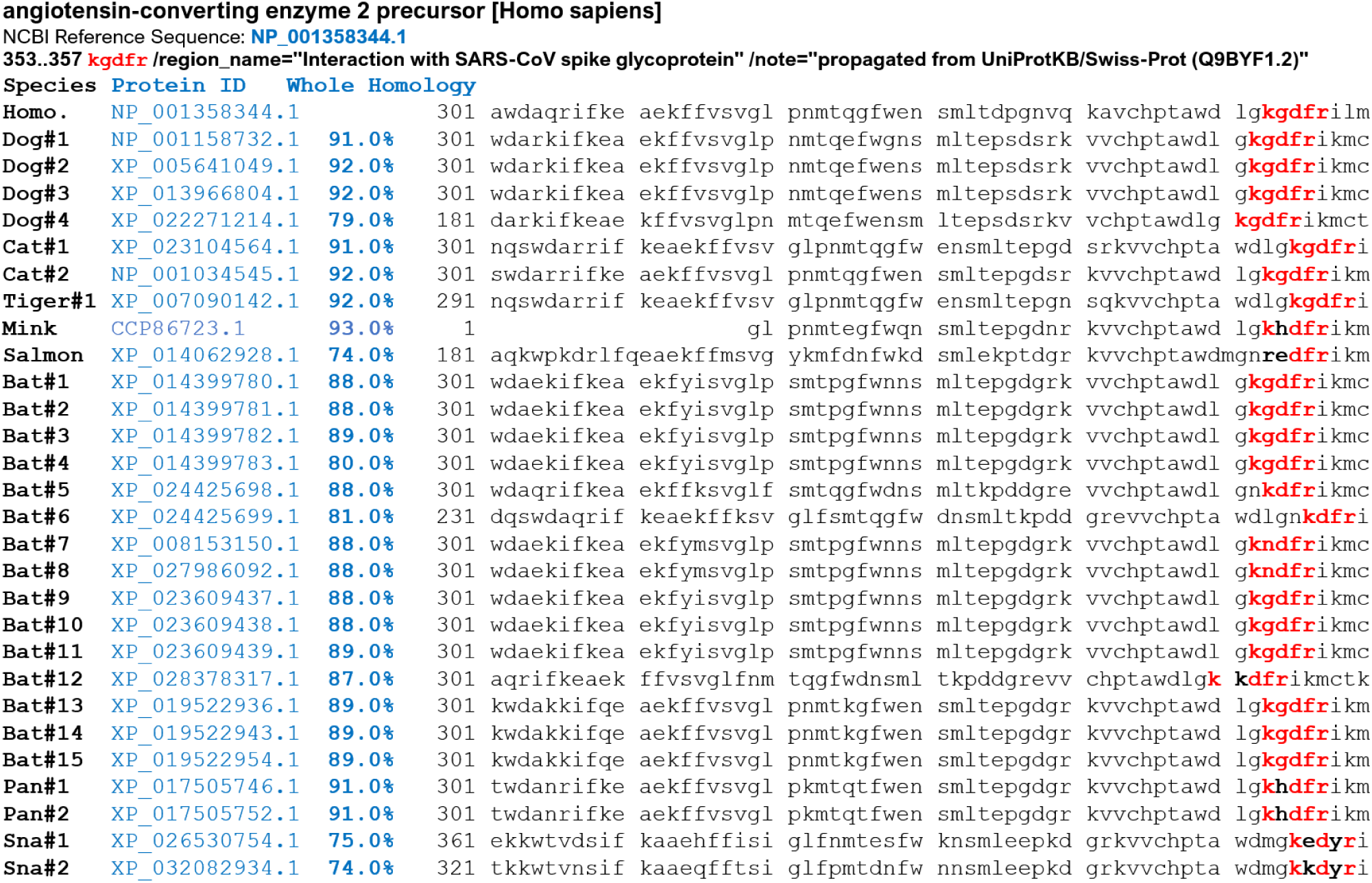
Whole amino acid molecule homology and amino acid sequence alignment of the binding region for an angiotensin-converting enzyme 2 (ACE2) as a receptor for SARS-CoV-2 spike glycoprotein and its phylogeny. Whole molecule homology and the homologous binding region of an angiotensin-converting enzyme 2 (ACE2) between human and other animal species including dogs, cats, tigers, mink, salmon, bats, pangolins, and snakes are indicated in Supplementary Materials. The key 5 amino acid residues KGDFR involved in the interaction with human SARS-CoV-2 spike glycoprotein are marked in red text. Detailed information regarding protein accession numbers of an ACE2 in other animal species can be found in the supplementary materials section.

In addition, we examined the homology of the five amino acid residues KGDFR located in the binding region of the ACE2 molecule, which directly recognize and bind the spike glycoprotein on the virion surface of SARS-CoV-2 in humans, dogs, cats, and other mammals. Our findings show that these five amino acid residues have 100% homology among humans, dogs, cats, tigers, and bats, 80% homology between humans and pangolins, and 60% homology between humans and snakes (Figure 1, Supplementary Materials). These five amino acid residues have 80% homology between humans and mink (Figure 1, Supplementary Materials). Our findings also show that these five amino acid residues have 60% homology between humans and salmon (Figure 1, Supplementary Materials). These results suggest that SARS-CoV-2 may transmit from humans to dogs, cats, tigers, and minks.

We analyzed the three-dimensional structure of the binding region in mink and human ACE2s for the spike glycoprotein of SARS-CoV-2 using Spanner, a structural homology modeling pipeline method (Supplementary Materials). The three-dimensional structure of the binding region containing five amino acids of mink ACE2 for the spike glycoprotein of SARS-CoV-2 was highly conserved in comparison with the structure of human ACE2 (Figure 2A). However, the three-dimensional structure of the binding region containing five amino acids of salmon ACE2 for the spike glycoprotein of SARS-CoV-2 was incompletely conserved in comparison with the structure of human ACE2 (Figure 2A). The analysis revealed that the three-dimensional structure of the binding region of salmon ACE2 is an imperfect match with the structure of the receptor binding domain (RBD) of the spike glycoprotein of SARS-CoV-2 (Figure 2B). In the case of salmon, the key (RBD) and the keyhole (ACE2-binding residues) do not completely match (Figure 2B). From these results, there is no medical or scientific evidence that SARS-CoV-2 infects salmon. Our research also suggests that SARS-CoV-2 zoonotic disease may spread worldwide (Figure 3). Our findings, alongside recent studies, may provide important insights into animal models for SARS-CoV-2 and animal management for COVID-19 control. The World Health Organization considers zoonotic infections to be a special case; however, it has called for individuals to also avoid cross-transmission of SARS-CoV-2 between humans, their pets, and other animals. The information provided from our examinations will support precision vaccine design and the discovery of antiviral therapeutics which will, in turn, accelerate the development of medical countermeasures.

**Figure 2.**
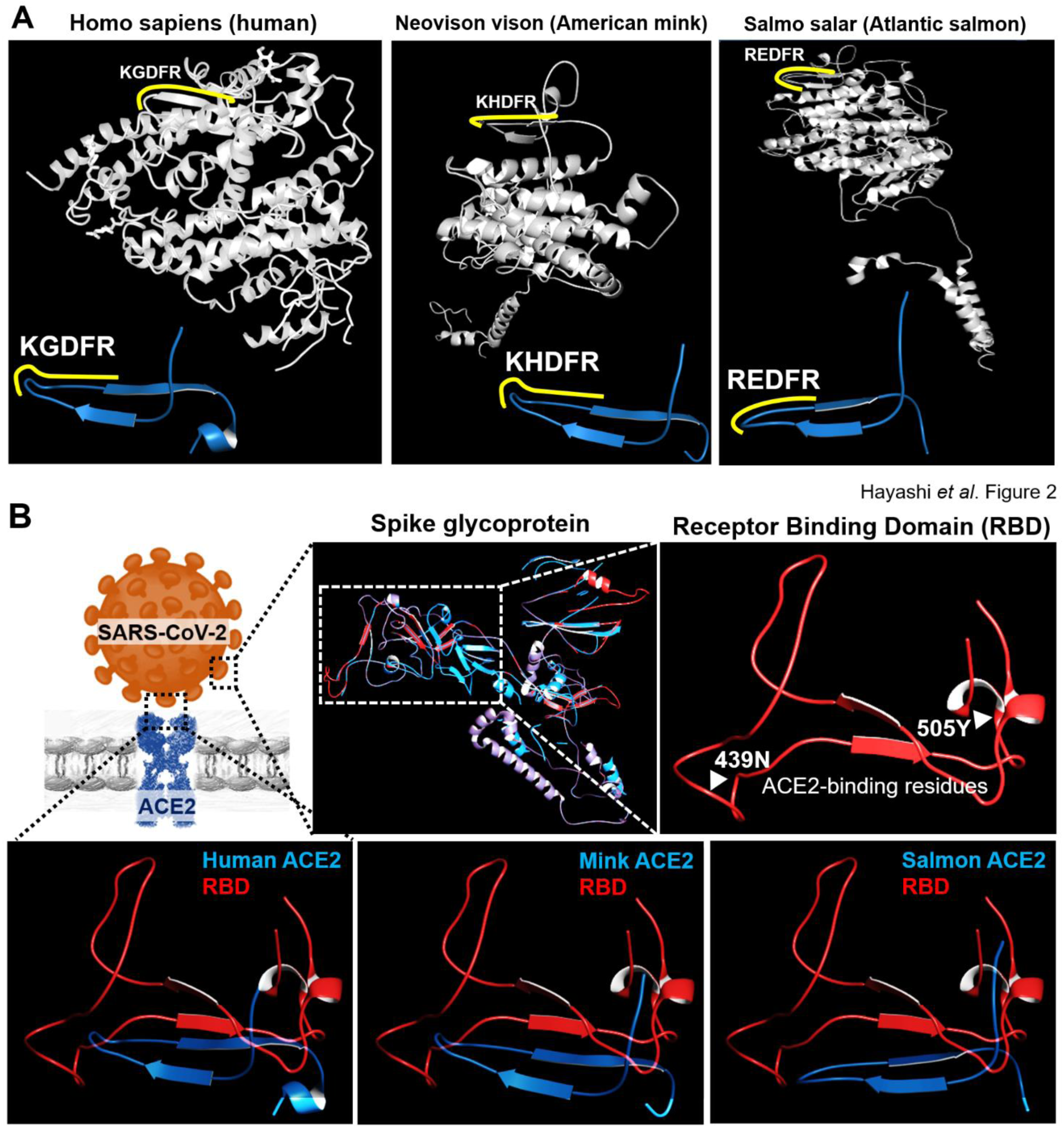
Examination of three-dimensional structure of binding region in mink and salmon ACE2 compared to human ACE2. Three-dimensional structures of the binding region for mink, salmon, and human ACE2s with the spike glycoprotein of SARS-CoV-2 using Spanner analysis, a structural homology modeling pipeline method (Supplementary Materials). **A**. The three-dimensional structure of the binding region containing five amino acids of mink ACE2 indicated by the yellow line for the spike glycoprotein of SARS-CoV-2 is conserved in comparison with the structure of human ACE2. The three-dimensional structure of the binding region containing five amino acids of salmon ACE2 indicated by the yellow line for the spike glycoprotein of SARS-CoV-2 is incompletely conserved in comparison with the structure of human ACE2. The ribbon diagram highlights the native ACE2 with the secondary structure and also the two subdomains (I and II) that form the two sides of the active site cleft. The two subdomains are defined as follows: The N terminus- and zinc-containing subdomain I, composed of residues 19–102, 290–397, and 417–430; and the C terminus-containing subdomain II, composed of residues 103–289, 398–416, and 431–615. Protein ID numbers NP_001358344.1 for human ACE2, CCP86723.1 for mink ACE2, and XP_014062928.1 for salmon ACE2 were used to analyze three-dimensional structures for mink, salmon, and human ACE2 molecules. **B**. Three-dimensional structures of the spike glycoprotein of SARS-CoV-2 using Spanner analysis (Supplementary Materials). Three-dimensional structures of receptor binding domain (RBD) of the spike glycoprotein of SARS-CoV-2 is displayed by the red line at the top right panel in B. Predictions of the complex structure of RBD (red line) with binding region including the five amino acids of human ACE2, mink ACE2, and salmon ACE2 (blue line) are shown under the panels in B. The results revealed the three-dimensional structure of the binding region of salmon ACE2 to be an imperfect match for the structure of RBD of the spike glycoprotein of SARS-CoV-2. Details of the amino acid sequence of mink, salmon, and human ACE2s are shown in the supplementary materials section.

**Figure 3.**
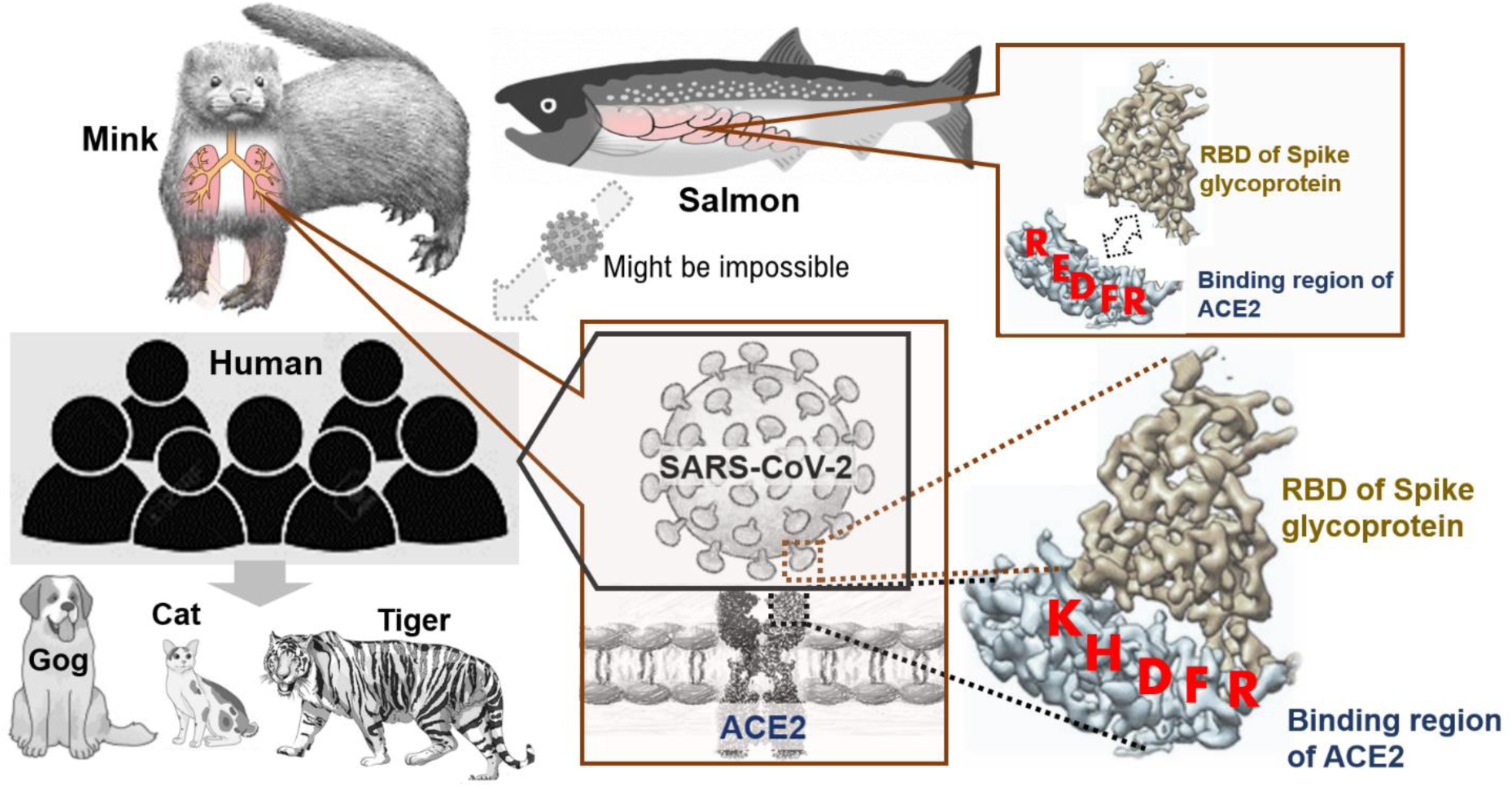
SARS-CoV-2 infection from mink or salmon to humans in zoonotic transmission. Currently, the exact route of zoonotic transmission of SARS-CoV-2 into the human population remains unknown. The three-dimensional structure of the binding region containing five amino acids of mink ACE2 for the spike glycoprotein of SARS-CoV-2 was highly conserved in comparison with the structure of human ACE2. However, the three-dimensional structure of the binding region containing five amino acids of salmon ACE2 for the spike glycoprotein of SARS-CoV-2 was incompletely conserved in comparison with the structure of human ACE2. From these research results, there is no medical and scientific evidence that SARS-CoV-2 infects salmon. Structural analysis suggests that multiple introductions of SARS-CoV-2 into the human population have occurred and both zoonotic transmission events and human-to-human transmission have driven the current COVID-19 outbreak.

## Discussion

A history of visits to the Beijing Fresh Fish Wholesale Market was confirmed in four patients with a confirmed diagnosis of SARS-CoV-2 infection on June 13, 2020. As a result, employees working at the market were voluntarily screened for SARS-CoV-2, with a positive reaction observed in 45 people.

Normally, SARS-CoV-2 infects the respiratory and intestinal tracts of homeotherms such as mammals and birds (7,8,9). The results of previous studies have not demonstrated that poikilotherms can be infected with the coronavirus. Fish are poikilotherms, and humans are homeotherms (10,11). In general, species that are closer in genetic classification are more susceptible to common infectious diseases. Due to the thermoregulatory mechanism of the species, the probability of transmission of a common infectious disease among homeotherms is higher than the rate of infection between homeotherms and poikilotherms animals (10,11). On the genetic classification, the difference in genetic background between humans and other poikilotherms such as fish, snakes, turtles, and soft-shelled turtles is much larger than that between humans and homeothermic animals such as birds and beasts (12).

However, in March 2020, the possibility of coronavirus infection in fish was reported. Sequences of coronavirus genes were confirmed in two separate cDNA pools (13). The initial cDNA sample pool was obtained from the *Carassius auratus* cell line (13). The second cDNA pool was obtained from *Ctenopharyngodon idella* kidney tissue (13). BLAST analysis revealed a DNA sequence very similar to SARS-CoV-2 in these cDNA pools (13), though these sequencing results might also be artifacts of the bioinformatics pipeline used in this study. However, it is possible that the aquatic habitat is an environmental pathogen of common coronaviruses such as SARS.

Amino acid loci similar to ACE2, which is a host-side receptor for mammalian SARS-CoV-2, have been found in poikilotherms animals. However, amphibians, reptiles, and mammals are not very closely genetically related. Our findings indicate that the homology of ACE2 between fish and humans is not high enough for coronavirus to infect fish (Figure 1). On the other hand, the homology of ACE2 between mammals including mink and humans is high, and cross-infection of coronavirus between different species is observed (Figure 1). Generally, it is considered that infectious diseases do not cross between terrestrial organisms and aquatic organisms (14). Therefore, it is considered that SARS-CoV-2 is unlikely to infect salmon.

It is possible that the surfaces of marine products and other foods were contaminated with SARS-CoV-2 in the new Beijing outbreak. In other words, it is possible that workers in food factories infected with SARS-CoV-2 were in contact with the marine products, allowing SARS-CoV-2 to adhere to their surface. It is known that virus can survive for long periods in a low-temperature environment, with previous studies having shown that coronaviruses can survive for years at minus 60 degrees (15,16). Therefore, SARS-CoV-2 may have been transported to the seafood market via the cold chain.

All cases of SARS-CoV-2 animal infections reported to date worldwide have been in mammals. Moreover, the transmission route of SARS-CoV-2 from humans to animals has been confirmed. In the United States, cases of infection from human to 5 dogs and lions three animals and pets 2 cats of the tiger have been reported (17-22). In Hong Kong, cases of infection of SARS-CoV-2 from human to two dogs and one cat have been reported. Medical evidence has been reported to suggest that COVID-19 is a zoonotic disease.

From the experiments to date, there has been no evidence to show the transmission of SARS-CoV-2 to salmon. Further research is needed to obtain medical evidence of this.

## Footnote

### ACE2-Angiotensin-Converting Enzyme 2

The protein encoded by this gene belongs to the angiotensin-converting enzyme family of dipeptidyl carboxypeptidases and has considerable homology with the human angiotensin-converting enzyme 1. This secreted protein catalyzes the cleavage of angiotensin I into angiotensin, and angiotensin II into the vasodilator angiotensin. The organ- and cell-specific expression of this gene suggests that it may play a role in the regulation of cardiovascular and renal functions, as well as fertility. In addition, the encoded protein is a functional receptor for the spike glycoprotein of the human coronaviruses SARS and HCoV-NL63. [Provided by RefSeq, Jul 2008: www.ncbi.nlm.nih.gov/gene/59272/ortholog/?scope=7776].

## Abbreviations

ACE2: Angiotensin-converting enzyme 2
COVID-19: Coronavirus disease 2019
NCBI: National Center for Biotechnology Information
SARS-CoV-2: Severe acute respiratory syndrome coronavirus 2

## Data Sharing

Data are available on various websites and have been made publicly available (more information can be found in the first paragraph of the Results section).

## Acknowledgments

We thank Professor Richard A. Young (Whitehead Institute for Biomedical Research, Massachusetts Institute of Technology, Cambridge, MA) for his research assistance. This study was supported in part by grants from the Japan Ministry of Education, Culture, Science and Technology (No. 24592510, No. 15K1079, and No. 19K09840), the Foundation of Osaka Cancer Research, The Ichiro Kanehara Foundation for the Promotion of Medical Science and Medical Care, the Foundation for Promotion of Cancer Research, the Kanzawa Medical Research Foundation, The Shinshu Medical Foundation, and the Takeda Foundation for Medical Science.

## Author Contributions

T.H. performed most of the experiments and coordinated the project. T.H. and M.M. conceived the study and wrote the manuscript. N.Y. and I.K. gave information on clinical medicine and oversaw the entire study.

## Transparency document

The transparency document associated with this article can be found in the online version at http://

