## supplemental materials for "Molecular analysis of binding region of an ACE2 as a receptor for SARS-CoV-2 between humans and mammals"

### Zoonotic disease: A highly conserved binding region of ACE2 as receptor for SARS-CoV-2 between human and mammals

#### METHODS

##### Phylogenetic Analysis and Annotation

Reference genomes and amino acids of human, dog, cat, tiger, bat, pangolin, and snake ACE2s were obtained from the National Center for Biotechnology Information (NCBI) Orthologs of the National Library of Medicine. Amino acid homological analysis was performed using Align Sequences Protein BLAST (algorithm protein–protein BLAST) with the protein accession numbers of ACE2s listed in the NCBI Reference Sequence Database in order to determine the whole amino acid homology of ACE2 between humans and other animals. Phylogenetic analyses of the complete protein and major coding regions were performed with RAxML software (version 8.2.9) with 1000 bootstrap replicates using the general time reversible nucleotide substitution model. Details of the protein accession numbers of ACE2s are available in the supplementary materials.

##### Amino Acid Homology Analysis of the Binding Region of ACE2 for Interaction with SARS-CoV-2 Spike Glycoprotein between Humans and Other Animals

The binding region for interaction with the SARS-CoV-2 spike glycoprotein (82.aa**-MYP-**84.aa, 353**.aa-KGDFR-**357.aa) of the verified genome amino acid sequences of human ACE2 was predicted using the NCBI protein database and Geneious software (version 11.1.5; Auckland, New Zealand), and was annotated using the NCBI Conserved Domain Database. Amino acid homological analysis was performed using Align Sequences Protein BLAST (algorithm protein–protein BLAST) with the protein accession numbers of human and individual animal ACE2s listed in the NCBI Reference Sequence Database. Details of the protein accession numbers of ACE2s are available in the supplementary materials.

**Analyzes the three-dimensional structure of the binding site between mink, Salmon and human ACE2**

Spanner is a structural homology modeling pipeline that threads a query amino-acid sequence onto a template protein structure. Spanner is unique in that it handles gaps by spanning the region of interest using fragments of known structures.

To create a model, you must provide a template structure, as well as an alignment of the query sequence you wish to model onto the template sequence. Spanner will replace mismatched residues, and fill any gaps caused by insertions or deletions.

For users that are unable to create an alignment a method for building a model starting only from sequence is also available. During this process a template search is conducted and an alignment is built dynamically using FORTE before being passed through to the main part of the pipeline.

Spanner consists of several modules written in the Go programming language. For Spanner jobs which build a model only from sequence, the first step is a search of the PDB for possible templates using BLAST. These possible templates are then aligned and scored with FORTE.

The next step involves defining the start and end points of fragments corresponding to insertions or deletions. The start and end points are referred to as anchors because they must be equivalent in both the template and any candidate fragment. The margin parameter determines how far from the edge of a gap the fragment begins or ends. For example a margin of 0 would mean that the anchors begins at the very edge of a gap. This is usually not a good idea, and the default margin is set to 1.

A representative set of protein chains was prepared using CD-HIT at 100% sequence identity.[3] All continuous fragments were then extracted from this set of chains and stored in a relational database, indexed by the internal coordinates of the fragment endpoints. A separate database is prepared for each fragment length. Currently, fragments of length 8-40, including the 8 anchor residues, are stored in the database.

**angiotensin-converting enzyme 2 precursor [Homo sapiens]**
NCBI Reference Sequence: NP_001358344.1

```
  1 mssssswllls lvavtaaqst ieeqaktfld kfnheaedlf yqsslaswny ntniteenvq
 61 nmnnagdkws aflkeqstla qmyplqeiqn ltvklqlqal qqngssvlse dkskrlntil
121 ntmstiystg kvcnpdnpqe clllepglne imansldyne rlwaweswrs evgkqlrply
181 eeyvvlknem aranhyedyg dywrgdyevn gvdgydysrg qliedvehtf eeikplyehl
241 hayvraklmn aypsyispig clpahllgdm wgrfwtnlys ltvpfgqkpn idvtdamvdq
301 awdaqrifke aekffvsvgl pnmtqgfwen smltdpgnvq kavchptawd lgkgdfrilm
361 ctkvtmddfl tahhemghiq ydmayaaqpf llrnganegf heavgeimsl saatpkhlks
421 igllspdfqe dneteinfll kqaltivgtl pftymlekwr wmvfkgeipk dqwmkkwwem
481 kreivgvvep vphdetycdp aslfhvsndy sfiryytrtl yqfqfqealc qaakhegplh
541 kcdisnstea gqklfnmlrl gksepwtlal envvgaknmn vrpllnyfep lftwlkdqnk
601 nsfvgwstdw spyadqsikv rislksalgd kayewndnem ylfrssvaya mrqyflkvkn
661 qmilfgeedv rvanlkpris fnffvtapkn vsdiiprtev ekairmsrsr indafrlndn
721 sleflgiqpt lgppnqppvs iwlivfgvvm gvivvgivil iftgirdrkk knkarsgenp
781 yasidiskge nnpgfqntdd vqtsf
```

82..84 **myp**, 353..357 **kgdfr**
/region_name="Interaction with SARS-CoV spike glycoprotein"
/experiment="experimental evidence, no additional details recorded"
/note="propagated from UniProtKB/Swiss-Prot (Q9BYF1.2)"

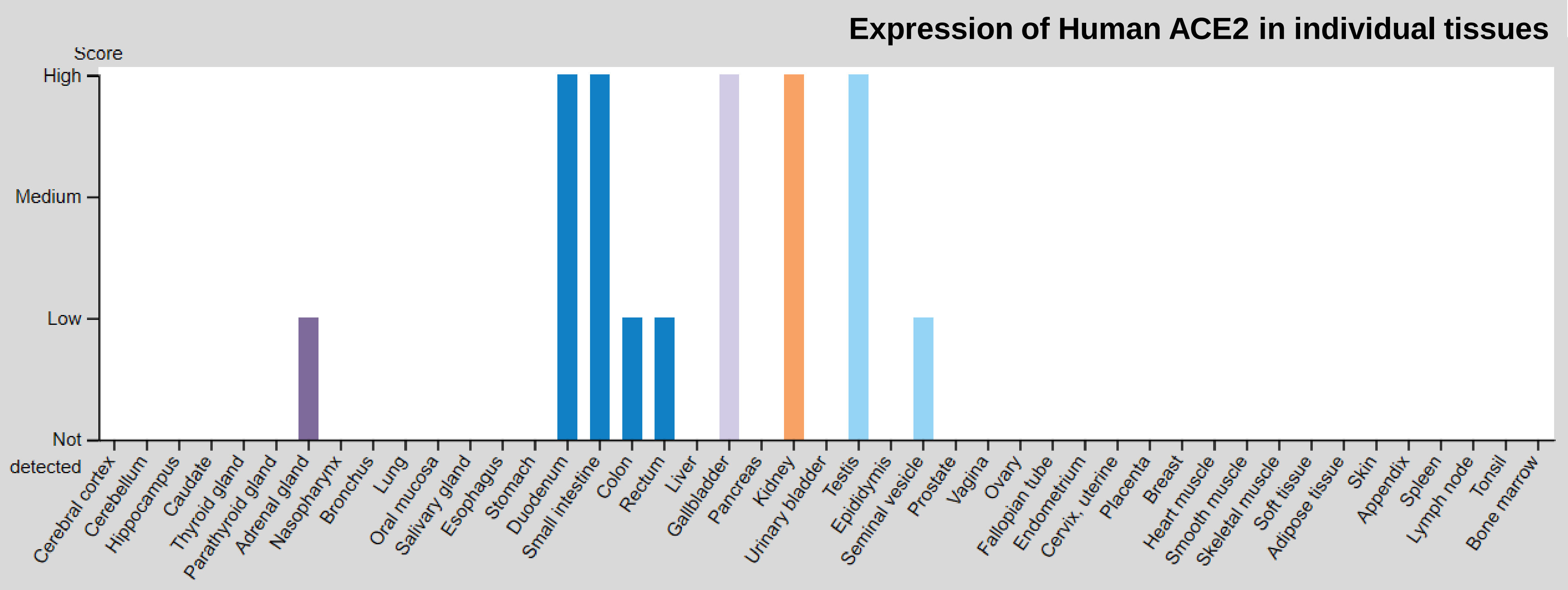


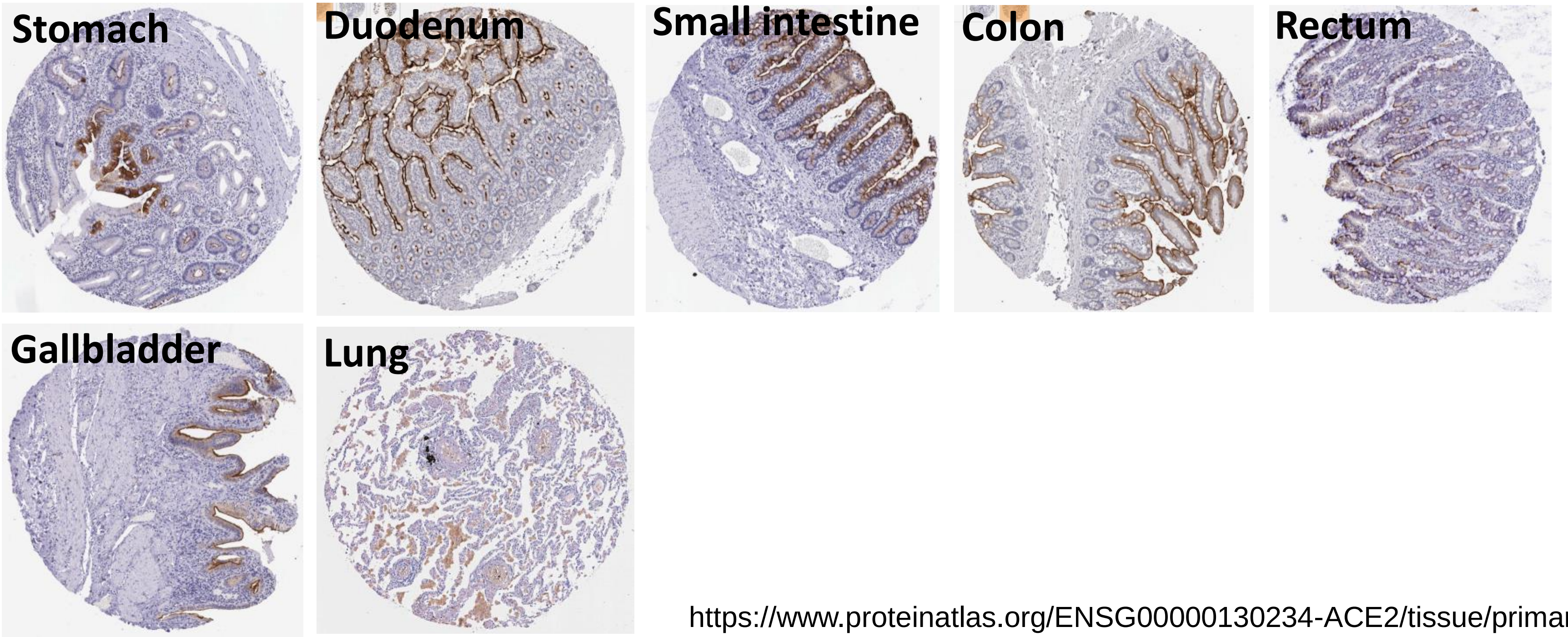


https://www.proteinatlas.org/ENSG00000130234-ACE2/tissue/primary+data

#### Global Alignment » results for RID-8B0TUUHX114 Human ACE2 vs Mammals ACE2

| Species | | | Protein ID | Homology |
|---|---|---|---|---|
| Human | [Homo sapiens] | NCBI Reference Sequence: | NP_001358344.1 | |
| Dog#1 | [Canis lupus familiaris] | NCBI Reference Sequence: | NP_001158732.1 | 91.0% |
| Dog#2 | [Canis lupus familiaris] | NCBI Reference Sequence: | XP_005641049.1 | 92.0% |
| Dog#3 | [Canis lupus familiaris] | NCBI Reference Sequence: | XP_013966804.1 | 92.0% |
| Dog#4 | [Canis lupus familiaris] | NCBI Reference Sequence: | XP_022271214.1 | 79.0% |
| Cat#1 | [Felis catus] | NCBI Reference Sequence: | XP_023104564.1 | 91.0% |
| Cat#2 | [Felis catus] | NCBI Reference Sequence: | NP_001034545.1 | 92.0% |
| Tiger#1 | [Panthera tigris altaica] | NCBI Reference Sequence: | XP_007090142.1 | 92.0% |
| Bat#1 | [Myotis brandtii] | NCBI Reference Sequence: | XP_014399780.1 | 88.0% |
| Bat#2 | [Myotis brandtii] | NCBI Reference Sequence: | XP_014399781.1 | 88.0% |
| Bat#3 | [Myotis brandtii] | NCBI Reference Sequence: | XP_014399782.1 | 89.0% |
| Bat#4 | [Myotis brandtii] | NCBI Reference Sequence: | XP_014399783.1 | 80.0% |
| Bat#5 | [Desmodus rotundus] | NCBI Reference Sequence: | XP_024425698.1 | 88.0% |
| Bat#6 | [Desmodus rotundus] | NCBI Reference Sequence: | XP_024425699.1 | 81.0% |
| Bat#7 | [Eptesicus fuscus] | NCBI Reference Sequence: | XP_008153150.1 | 88.0% |
| Bat#8 | [Eptesicus fuscus] | NCBI Reference Sequence: | XP_027986092.1 | 88.0% |
| Bat#9 | [Myotis lucifugus] | NCBI Reference Sequence: | XP_023609437.1 | 88.0% |
| Bat#10 | [Myotis lucifugus] | NCBI Reference Sequence: | XP_023609438.1 | 88.0% |
| Bat#11 | [Myotis lucifugus] | NCBI Reference Sequence: | XP_023609439.1 | 89.0% |
| Bat#12 | [Phyllostomus discolor] | NCBI Reference Sequence: | XP_028378317.1 | 87.0% |
| Bat#13 | [Hipposideros armiger] | NCBI Reference Sequence: | XP_019522936.1 | 89.0% |
| Bat#14 | [Hipposideros armiger] | NCBI Reference Sequence: | XP_019522943.1 | 89.0% |
| Bat#15 | [Hipposideros armiger] | NCBI Reference Sequence: | XP_019522954.1 | 89.0% |
| Pangolin#1 | [Manis javanica] | NCBI Reference Sequence: | XP_017505746.1 | 91.0% |
| Pangolin#2 | [Manis javanica] | NCBI Reference Sequence: | XP_017505752.1 | 91.0% |
| Snake#1 | [Notechis scutatus] | NCBI Reference Sequence: | XP_026530754.1 | 75.0% |
| Snake#2 | [Thamnophis elegans] | NCBI Reference Sequence: | XP_032082934.1 | 74.0% |

https://blast.ncbi.nlm.nih.gov/Blast.cgi

**angiotensin-converting enzyme 2 precursor [Homo sapiens]**

NCBI Reference Sequence: NP_001358344.1

**353..357 kgdfr /region_name="Interaction with SARS-CoV spike glycoprotein"**

**/note="propagated from UniProtKB/Swiss-Prot (Q9BYF1.2)"**

| | | | | | | | |
|---|---|---|---|---|---|---|---|
| **Homo.** | 301 | awdaqrifke | aekffvsvgl | pnmtqgfwen | smltdpgnvq | kavchptawd | lg**kgdfr**ilm |
| **Dog#1** | 301 | wdarkifkea | ekffvsvglp | nmtqefwgns | mltepsdsrk | vvchptawdl | g**kgdfr**ikmc |
| **Dog#2** | 301 | wdarkifkea | ekffvsvglp | nmtqefwens | mltepsdsrk | vvchptawdl | g**kgdfr**ikmc |
| **Dog#3** | 301 | wdarkifkea | ekffvsvglp | nmtqefwens | mltepsdsrk | vvchptawdl | g**kgdfr**ikmc |
| **Dog#4** | 181 | darkifkeae | kffvsvglpn | mtqefwensm | ltepsdsrkv | vchptawdlg | **kgdfr**ikmct |
| **Cat#1** | 301 | nqswdarrif | keaekffvsv | glpnmtqgfw | ensmltepgd | srkvvchpta | wdlg**kgdfr**i |
| **Cat#2** | 301 | swdarrifke | aekffvsvgl | pnmtqgfwen | smltepgdsr | kvvchptawd | lg**kgdfr**ikm |
| **Tiger#1** | 291 | nqswdarrif | keaekffvsv | glpnmtqgfw | ensmltepgn | sqkvvchpta | wdlg**kgdfr**i |
| **Bat#1** | 301 | wdaekifkea | ekfyisvglp | smtpgfwnns | mltepgdgrk | vvchptawdl | g**kgdfr**ikmc |
| **Bat#2** | 301 | wdaekifkea | ekfyisvglp | smtpgfwnns | mltepgdgrk | vvchptawdl | g**kgdfr**ikmc |
| **Bat#3** | 301 | wdaekifkea | ekfyisvglp | smtpgfwnns | mltepgdgrk | vvchptawdl | g**kgdfr**ikmc |
| **Bat#4** | 301 | wdaekifkea | ekfyisvglp | smtpgfwnns | mltepgdgrk | vvchptawdl | g**kgdfr**ikmc |
| **Bat#5** | 301 | wdaqrifkea | ekffksvglf | smtqgfwdns | mltkpddgre | vvchptawdl | gn**kdfr**ikmc |
| **Bat#6** | 231 | dqswdaqrif | keaekffksv | glfsmtqgfw | dnsmltkpdd | grevvchpta | wdlgn**kdfr**i |
| **Bat#7** | 301 | wdaekifkea | ekfymsvglp | smtpgfwnns | mltepgdgrk | vvchptawdl | g**kndfr**ikmc |
| **Bat#8** | 301 | wdaekifkea | ekfymsvglp | smtpgfwnns | mltepgdgrk | vvchptawdl | g**kndfr**ikmc |
| **Bat#9** | 301 | wdaekifkea | ekfyisvglp | smtpgfwnns | mltepgdgrk | vvchptawdl | g**kgdfr**ikmc |
| **Bat#10** | 301 | wdaekifkea | ekfyisvglp | smtpgfwnns | mltepgdgrk | vvchptawdl | g**kgdfr**ikmc |
| **Bat#11** | 301 | wdaekifkea | ekfyisvglp | smtpgfwnns | mltepgdgrk | vvchptawdl | g**kgdfr**ikmc |
| **Bat#12** | 301 | aqrifkeaek | ffvsvglfnm | tqgfwdnsml | tkpddgrevv | chptawdlg**k** | **kdfr**ikmctk |
| **Bat#13** | 301 | kwdakkifqe | aekffvsvgl | pnmtkgfwen | smltepgdgr | kvvchptawd | lg**kgdfr**ikm |
| **Bat#14** | 301 | kwdakkifqe | aekffvsvgl | pnmtkgfwen | smltepgdgr | kvvchptawd | lg**kgdfr**ikm |
| **Bat#15** | 301 | kwdakkifqe | aekffvsvgl | pnmtkgfwen | smltepgdgr | kvvchptawd | lg**kgdfr**ikm |
| **Pan#1** | 301 | twdanrifke | aekffvsvgl | pkmtqtfwen | smltepgdgr | kvvchptawd | lg**khdfr**ikm |
| **Pan#2** | 301 | twdanrifke | aekffvsvgl | pkmtqtfwen | smltepgdgr | kvvchptawd | lg**khdfr**ikm |
| **Sna#1** | 361 | ekkwtvdsif | kaaehffisi | glfnmtesfw | knsmleepkd | grkvvchpta | wdmg**kedyr**i |
| **Sna#2** | 321 | tkkwtvnsif | kaaeqfftsi | glfpmtdnfw | nnsmleepkd | grkvvchpta | wdmg**kkdyr**i |

**angiotensin-converting enzyme 2 precursor [Homo sapiens]**

NCBI Reference Sequence: **NP_001358344.1**

**353..357 kgdfr /region_name="Interaction with SARS-CoV spike glycoprotein"**

**/note="propagated from UniProtKB/Swiss-Prot (Q9BYF1.2)"**

```
Homo.    301 awdaqrifke aekffvsvgl pnmtqgfwen smltdpgnvq kavchptawd lgkgdfrilm
mink       1                    gl pnmtegfwqn smltepgdnr kvvchptawd lgkhdfrikm
salmon   181 aqkwpkdrlfqeaekffmsvg ykmfdnfwkd smlekptdgr kvvchptawdmgnredfrikm
```

#### angiotensin-converting enzyme 2 precursor [Canis lupus familiaris]

Sequence ID: NP_001158732.1 Length: 804 Number of Matches: 1

See 1 more title(s)

angiotensin I converting enzyme 2 [Canis lupus familiaris]
Sequence ID: ACT66277.1

**Human vs Dog**

**Range 1: 1 to 804** GenPept Graphics Next Match

| NW Score | Identities | Positives | Gaps |
|---|---|---|---|
| 3632 | 672/805(83%) | 739/805(91%) | 1/805(0%) |

```
Query  1    MSSSSWLLLSLVAVTAAQSTIEEQAKTFLDKFNHEAEDLFYQSSLASWNYNTNITEENVQ  60
            MS SSWLLLSL A+TAAQST E+  KTFL+KFN+EAE+L YQSSLASWNYN NIT+ENVQ
Sbjct  1    MSGSSWLLLSLAALTAAQST-EDLVKTFLEKFNYEAEELSYQSSLASWNYNINITDENVQ  59

Query  61   NMNNAGDKWSAFLKEQSTLAQMYPLQEIQNLTVKLQLQALQQNGSSVLSEDKSKRLNTIL  120
             MNNAG KWSAF +EQS LA+ YPL+EIQ+ TVK QL+ALQ +GSSVLS DK++RLNTIL
Sbjct  60   KMNNAGAKWSAFYEEQSKLAKTYPLEEIQDSTVKRQLRALQHSGSSVLSADKNQRLNTIL  119

Query  121  NTMSTIYSTGKVCNPDNPQECLLLEPGLNEIMANSLDYNERLWAWESWRSEVGKQLRPLY  180
            N+MST+YSTGK CNP NPQECLLLEPGL++IM NS DYNERLWAWE WRSEVGKQLRPLY
Sbjct  120  NSMSTVYSTGKACNPSNPQECLLLEPGLDDIMENSKDYNERLWAWEGWRSEVGKQLRPLY  179

Query  181  EEYVVLKNEMARANHYEDYGDYWRGDYEVNGVDGYDYSRGQLIEDVEHTFEEIKPLYEHL  240
            EEYV LKNEMARAN+YEDYGDYWRGDYE    +GY+YSR QLI+DVE TF +I PLY+HL
Sbjct  180  EEYVALKNEMARANNYEDYGDYWRGDYEEEWENGYNYSRNQLIDDVELTFTQIMPLYQHL  239

Query  241  HAYVRAKLMNAYPSYISPIGCLPAHLLGDMWGRFWTNLYSLTVPFGQKPNIDVTDAMVDQ  300
            HAYVR KLM+ YPSYISP GCLPAHLLGDMWGRFWTNLY LTVPFGQKPNIDVT+AMV+Q
Sbjct  240  HAYVRTKLMDTYPSYISPTGCLPAHLLGDMWGRFWTNLYPLTVPFGQKPNIDVTNAMVNQ  299

Query  301  AWDAQRIFKEAEKFFVSVGLPNMTQGFWENSMLTDPGNVQKAVCHPTAWDLGKGDFRILM  360
            +WDA++IFKEAEKFFVSVGLPNMTQ FW NSMLT+P + +K VCHPTAWDLGKGDFRI M
Sbjct  300  SWDARKIFKEAEKFFVSVGLPNMTQEFWGNSMLTEPSDSRKVVCHPTAWDLGKGDFRIKM  359

Query  361  CTKVTMDDFLTAHHEMGHIQYDMAYAAQPFLLRNGANEGFHEAVGEIMSLSAATPKHLKS  420
            CTKVTMDDFLTAHHEMGHIQYDMAYAAQPFLLRNGANEGFHEAVGEIMSLSAATP HLK+
Sbjct  360  CTKVTMDDFLTAHHEMGHIQYDMAYAAQPFLLRNGANEGFHEAVGEIMSLSAATPNHLKN  419

Query  421  IGLLSPDFQEDNETEINFLLKQALTIVGTLPFTYMLEKWRWMVFKGEIPKDQWMKKWWEM  480
            IGLL P F ED+ETEINFLLKQALTIVGTLPFTYMLEKWRWMVFKGEIPKDQWMK WWEM
Sbjct  420  IGLLPPSFFEDSETEINFLLKQALTIVGTLPFTYMLEKWRWMVFKGEIPKDQWMKTWWEM  479

Query  481  KREIVGVVEPVPHDETYCDPASLFHVSNDYSFIRYYTRTLYQFQFQEALCQAAKHEGPLH  540
            KR IVGVVEPVPHDETYCDPASLFHV+NDYSFIRYYTRT+YQFQFQEALCQ AKHEGPLH
Sbjct  480  KRNIVGVVEPVPHDETYCDPASLFHVANDYSFIRYYTRTIYQFQFQEALCQIAKHEGPLH  539

Query  541  KCDISNSTEAGQKLFNMLRLGKSEPWTLALENVVGAKNMNVRPLLNYFEPLFTWLKDQNK  600
            KCDISNS+EAGQKL  ML+LGKS+PWT ALE VVGAKNM+VRPLLNYFEPLFTWLK+QN+
Sbjct  540  KCDISNSSEAGQKLLEMLKLGKSKPWTYALEIVVGAKNMDVRPLLNYFEPLFTWLKEQNR  599

Query  601  NSFVGWSTDWSPYADQSIKVRISLKSALGDKAYEWNDNEMYLFRSSVAYAMRQYFLKVKN  660
            NSFVGW+TDWSPYADQSIKVRISLKSALG+KAYEWN+NEMYLFRSS+AYAMRQYF +VKN
Sbjct  600  NSFVGWNTDWSPYADQSIKVRISLKSALGEKAYEWNNNEMYLFRSSIAYAMRQYFSEVKN  659

Query  661  QMILFGEEDVRVANLKPRISFNFFVTAPKNVSDIIPRTEVEKAIRMSRSRINDAFRLNDN  720
            Q I F E++V V++LKPRISFNF VT+P NVSDIIPRTEVE+AIRM RSRIND FRL+DN
Sbjct  660  QTIPFVEDNVWVSDLKPRISFNFSVTSPGNVSDIIPRTEVEEAIRMYRSRINDVFRLDDN  719

Query  721  SLEFLGIQPTLGPPNQPPVSIWLIVFGVVMGVIVVGIVILIFTGIRDRKKKNKARSGENP  780
            SLEFLGIQPT GPP +PPV+IWLIVFGVVMGV+VVGIV+LIF+GIR+R+K ++AR  ENP
Sbjct  720  SLEFLGIQPTPGPPYEPPVTIWLIVFGVVMGVVVVGIVLLIFSGIRNRRKNDQARGEENP  779

Query  781  YASIDISKGENNPGFQNTDDVQTSF  805
            YAS+D+SKGENNPGFQ+ DDVQTSF
Sbjct  780  YASVDLSKGENNPGFQSGDDVQTSF  804
```

#### angiotensin-converting enzyme 2 isoform X1 [Canis lupus familiaris]

Sequence ID: XP_005641049.1 Length: 804 Number of Matches: 1

See 1 more title(s)

angiotensin-converting enzyme 2 isoform X1 [Canis lupus familiaris]
Sequence ID: XP_013966804.1

**Human vs Dog**

**Range 1: 1 to 804** GenPept Graphics Next Match

| NW Score | Identities | Positives | Gaps |
|---|---|---|---|
| 3669 | 677/805(84%) | 742/805(92%) | 1/805(0%) |

```
Query  1    MSSSSWLLLSLVAVTAAQSTIEEQAKTFLDKFNHEAEDLFYQSSLASWNYNTNITEENVQ  60
            MS SSWLLLSL A+TAAQST E+  KTFL+KFN+EAE+L YQSSLASWNYN NIT+ENVQ
Sbjct  1    MSGSSWLLLSLAALTAAQST-EDLVKTFLEKFNYEAEELSYQSSLASWNYNINITDENVQ  59

Query  61   NMNNAGDKWSAFLKEQSTLAQMYPLQEIQNLTVKLQLQALQQNGSSVLSEDKSKRLNTIL  120
             MNNAG KWSAF +EQS LA+ YPL+EIQ+ TVK QL+ALQ +GSSVLS DK++RLNTIL
Sbjct  60   KMNNAGAKWSAFYEEQSKLAKTYPLEEIQDSTVKRQLRALQHSGSSVLSADKNQRLNTIL  119

Query  121  NTMSTIYSTGKVCNPDNPQECLLLEPGLNEIMANSLDYNERLWAWESWRSEVGKQLRPLY  180
            N+MSTIYSTGK CNP NPQECLLLEPGL++IM NS DYNERLWAWE WRSEVGKQLRPLY
Sbjct  120  NSMSTIYSTGKACNPSNPQECLLLEPGLDDIMENSKDYNERLWAWEGWRSEVGKQLRPLY  179

Query  181  EEYVVLKNEMARANHYEDYGDYWRGDYEVNGVDGYDYSRGQLIEDVEHTFEEIKPLYEHL  240
            EEYV LKNEMARAN+YEDYGDYWRGDYE    +GY+YSR QLI+DVEHTF +I PLY+HL
Sbjct  180  EEYVALKNEMARANNYEDYGDYWRGDYEEEWENGYNYSRNQLIDDVEHTFTQIMPLYQHL  239

Query  241  HAYVRAKLMNAYPSYISPIGCLPAHLLGDMWGRFWTNLYSLTVPFGQKPNIDVTDAMVDQ  300
            HAYVR KLM+ YPSYISP GCLPAHLLGDMWGRFWTNLY LTVPFGQKPNIDVT+AMV+Q
Sbjct  240  HAYVRTKLMDTYPSYISPTGCLPAHLLGDMWGRFWTNLYPLTVPFGQKPNIDVTNAMVNQ  299

Query  301  AWDAQRIFKEAEKFFVSVGLPNMTQGFWENSMLTDPGNVQKAVCHPTAWDLGKGDFRILM  360
            +WDA++IFKEAEKFFVSVGLPNMTQ FWENSMLT+P + +K VCHPTAWDLGKGDFRI M
Sbjct  300  SWDARKIFKEAEKFFVSVGLPNMTQEFWENSMLTEPSDSRKVVCHPTAWDLGKGDFRIKM  359

Query  361  CTKVTMDDFLTAHHEMGHIQYDMAYAAQPFLLRNGANEGFHEAVGEIMSLSAATPKHLKS  420
            CTKVTMDDFLTAHHEMGHIQYDMAYAAQPFLLRNGANEGFHEAVGEIMSLSAATP HLK+
Sbjct  360  CTKVTMDDFLTAHHEMGHIQYDMAYAAQPFLLRNGANEGFHEAVGEIMSLSAATPNHLKN  419

Query  421  IGLLSPDFQEDNETEINFLLKQALTIVGTLPFTYMLEKWRWMVFKGEIPKDQWMKKWWEM  480
            IGLL P F ED+ETEINFLLKQALTIVGTLPFTYMLEKWRWMVFKGEIPKDQWMK WWEM
Sbjct  420  IGLLPPSFFEDSETEINFLLKQALTIVGTLPFTYMLEKWRWMVFKGEIPKDQWMKTWWEM  479

Query  481  KREIVGVVEPVPHDETYCDPASLFHVSNDYSFIRYYTRTLYQFQFQEALCQAAKHEGPLH  540
            KR IVGVVEPVPHDETYCDPASLFHV+NDYSFIRYYTRT+YQFQFQEALCQ AKHEGPLH
Sbjct  480  KRNIVGVVEPVPHDETYCDPASLFHVANDYSFIRYYTRTIYQFQFQEALCQIAKHEGPLH  539

Query  541  KCDISNSTEAGQKLFNMLRLGKSEPWTLALENVVGAKNMNVRPLLNYFEPLFTWLKDQNK  600
            KCDISNS+EAGQKL  ML+LGKS+PWT ALE VVGAKNM+VRPLLNYFEPLFTWLK+QN+
Sbjct  540  KCDISNSSEAGQKLLEMLKLGKSKPWTYALEIVVGAKNMDVRPLLNYFEPLFTWLKEQNR  599

Query  601  NSFVGWSTDWSPYADQSIKVRISLKSALGDKAYEWNDNEMYLFRSSVAYAMRQYFLKVKN  660
            NSFVGW+TDWSPYADQSIKVRISLKSALG+KAYEWN+NEMYLFRSS+AYAMRQYF +VKN
Sbjct  600  NSFVGWNTDWSPYADQSIKVRISLKSALGEKAYEWNNNEMYLFRSSIAYAMRQYFSEVKN  659

Query  661  QMILFGEEDVRVANLKPRISFNFFVTAPKNVSDIIPRTEVEKAIRMSRSRINDAFRLNDN  720
            Q I F E++V V++LKPRISFNFFVT+P NVSDIIPRTEVE+AIRM RSRIND FRL+DN
Sbjct  660  QTIPFVEDNVWVSDLKPRISFNFFVTSPGNVSDIIPRTEVEEAIRMYRSRINDVFRLDDN  719

Query  721  SLEFLGIQPTLGPPNQPPVSIWLIVFGVVMGVIVVGIVILIFTGIRDRKKKNKARSGENP  780
            SLEFLGIQPTLGPP +PPV+IWLIVFGVVMGV+VVGIV+LIF+GIR+R+K ++AR  ENP
Sbjct  720  SLEFLGIQPTLGPPYEPPVTIWLIVFGVVMGVVVVGIVLLIFSGIRNRRKNDQARGEENP  779

Query  781  YASIDISKGENNPGFQNTDDVQTSF  805
            YAS+D+SKGENNPGFQN DD QTSF
Sbjct  780  YASVDLSKGENNPGFQNVDDAQTSF  804
```

**angiotensin-converting enzyme 2 isoform X1 [Canis lupus familiaris]**

Sequence ID: XP_005641049.1 Length: 804 Number of Matches: 1

See 1 more title(s)

angiotensin-converting enzyme 2 isoform X1 [Canis lupus familiaris]
Sequence ID: XP_013966804.1

#### Human vs Dog

Range 1: 1 to 804 GenPept Graphics Next Match

| NW Score | Identities | Positives | Gaps |
|---|---|---|---|
| 3669 | 677/805(84%) | 742/805(92%) | 1/805(0%) |

```
Query  1    MSSSSWLLLSLVAVTAAQSTIEEQAKTFLDKFNHEAEDLFYQSSLASWNYNTNITEENVQ  60
            MS SSWLLLSL A+TAAQST E+  KTFL+KFN+EAE+L YQSSLASWNYN NIT+ENVQ
Sbjct  1    MSGSSWLLLSLAALTAAQST-EDLVKTFLEKFNYEAEELSYQSSLASWNYNINITDENVQ  59

Query  61   NMNNAGDKWSAFLKEQSTLAQMYPLQEIQNLTVKLQLQALQQNGSSVLSEDKSKRLNTIL  120
             MNNAG KWSAF +EQS LA+ YPL+EIQ+ TVK QL+ALQ +GSSVLS DK++RLNTIL
Sbjct  60   KMNNAGAKWSAFYEEQSKLAKTYPLEEIQDSTVKRQLRALQHSGSSVLSADKNQRLNTIL  119

Query  121  NTMSTIYSTGKVCNPDNPQECLLLEPGLNEIMANSLDYNERLWAWESWRSEVGKQLRPLY  180
            N+MSTIYSTGK CNP NPQECLLLEPGL++IM NS DYNERLWAWE WRSEVGKQLRPLY
Sbjct  120  NSMSTIYSTGKACNPSNPQECLLLEPGLDDIMENSKDYNERLWAWEGWRSEVGKQLRPLY  179

Query  181  EEYVVLKNEMARANHYEDYGDYWRGDYEVNGVDGYDYSRGQLIEDVEHTFEEIKPLYEHL  240
            EEYV LKNEMARAN+YEDYGDYWRGDYE    +GY+YSR QLI+DVEHTF +I PLY+HL
Sbjct  180  EEYVALKNEMARANNYEDYGDYWRGDYEEEWENGYNYSRNQLIDDVEHTFTQIMPLYQHL  239

Query  241  HAYVRAKLMNAYPSYISPIGCLPAHLLGDMWGRFWTNLYSLTVPFGQKPNIDVTDAMVDQ  300
            HAYVR KLM+ YPSYISP GCLPAHLLGDMWGRFWTNLY LTVPFGQKPNIDVT+AMV+Q
Sbjct  240  HAYVRTKLMDTYPSYISPTGCLPAHLLGDMWGRFWTNLYPLTVPFGQKPNIDVTNAMVNQ  299

Query  301  AWDAQRIFKEAEKFFVSVGLPNMTQGFWENSMLTDPGNVQKAVCHPTAWDLGKGDFRILM  360
            +WDA++IFKEAEKFFVSVGLPNMTQ FWENSMLT+P + +K VCHPTAWDLGKGDFRI M
Sbjct  300  SWDARKIFKEAEKFFVSVGLPNMTQEFWENSMLTEPSDSRKVVCHPTAWDLGKGDFRIKM  359

Query  361  CTKVTMDDFLTAHHEMGHIQYDMAYAAQPFLLRNGANEGFHEAVGEIMSLSAATPKHLKS  420
            CTKVTMDDFLTAHHEMGHIQYDMAYAAQPFLLRNGANEGFHEAVGEIMSLSAATP HLK+
Sbjct  360  CTKVTMDDFLTAHHEMGHIQYDMAYAAQPFLLRNGANEGFHEAVGEIMSLSAATPNHLKN  419

Query  421  IGLLSPDFQEDNETEINFLLKQALTIVGTLPFTYMLEKWRWMVFKGEIPKDQWMKKWWEM  480
            IGLL P F ED+ETEINFLLKQALTIVGTLPFTYMLEKWRWMVFKGEIPKDQWMK WWEM
Sbjct  420  IGLLPPSFFEDSETEINFLLKQALTIVGTLPFTYMLEKWRWMVFKGEIPKDQWMKTWWEM  479

Query  481  KREIVGVVEPVPHDETYCDPASLFHVSNDYSFIRYYTRTLYQFQFQEALCQAAKHEGPLH  540
            KR IVGVVEPVPHDETYCDPASLFHV+NDYSFIRYYTRT+YQFQFQEALCQ AKHEGPLH
Sbjct  480  KRNIVGVVEPVPHDETYCDPASLFHVANDYSFIRYYTRTIYQFQFQEALCQIAKHEGPLH  539

Query  541  KCDISNSTEAGQKLFNMLRLGKSEPWTLALENVVGAKNMNVRPLLNYFEPLFTWLKDQNK  600
            KCDISNS+EAGQKL  ML+LGKS+PWT ALE VVGAKNM+VRPLLNYFEPLFTWLK+QN+
Sbjct  540  KCDISNSSEAGQKLLEMLKLGKSKPWTYALEIVVGAKNMDVRPLLNYFEPLFTWLKEQNR  599

Query  601  NSFVGWSTDWSPYADQSIKVRISLKSALGDKAYEWNDNEMYLFRSSVAYAMRQYFLKVKN  660
            NSFVGW+TDWSPYADQSIKVRISLKSALG+KAYEWN+NEMYLFRSS+AYAMRQYF +VKN
Sbjct  600  NSFVGWNTDWSPYADQSIKVRISLKSALGEKAYEWNNNEMYLFRSSIAYAMRQYFSEVKN  659

Query  661  QMILFGEEDVRVANLKPRISFNFFVTAPKNVSDIIPRTEVEKAIRMSRSRINDAFRLNDN  720
            Q I F E++V V++LKPRISFNFFVT+P NVSDIIPRTEVE+AIRM RSRIND FRL+DN
Sbjct  660  QTIPFVEDNVWVSDLKPRISFNFFVTSPGNVSDIIPRTEVEEAIRMYRSRINDVFRLDDN  719

Query  721  SLEFLGIQPTLGPPNQPPVSIWLIVFGVVMGVIVVGIVILIFTGIRDRKKKNKARSGENP  780
            SLEFLGIQPTLGPP +PPV+IWLIVFGVVMGV+VVGIV+LIF+GIR+R+K ++AR  ENP
Sbjct  720  SLEFLGIQPTLGPPYEPPVTIWLIVFGVVMGVVVVGIVLLIFSGIRNRRKNDQARGEENP  779

Query  781  YASIDISKGENNPGFQNTDDVQTSF  805
            YAS+D+SKGENNPGFQN DD QTSF
Sbjct  780  YASVDLSKGENNPGFQNVDDAQTSF  804
```

**angiotensin-converting enzyme 2 isoform X2 [Canis lupus familiaris]**

Sequence ID: XP_022271214.1 Length: 683 Number of Matches: 1

See 1 more title(s)

angiotensin-converting enzyme 2 isoform X2 [Canis lupus dingo]
Sequence ID: XP_025292937.1

#### Human vs Dog

Range 1: 1 to 683 GenPept Graphics Next Match

| NW Score | Identities | Positives | Gaps |
|---|---|---|---|
| 3088 | 586/805(73%) | 636/805(79%) | 122/805(15%) |

```
Query  1    MSSSSWLLLSLVAVTAAQSTIEEQAKTFLDKFNHEAEDLFYQSSLASWNYNTNITEENVQ  60
            MS
Sbjct  1    MS----------------------------------------------------------  2

Query  61   NMNNAGDKWSAFLKEQSTLAQMYPLQEIQNLTVKLQLQALQQNGSSVLSEDKSKRLNTIL  120

Sbjct       ------------------------------------------------------------

Query  121  NTMSTIYSTGKVCNPDNPQECLLLEPGLNEIMANSLDYNERLWAWESWRSEVGKQLRPLY  180
                TIYSTGK CNP NPQECLLLEPGL++IM NS DYNERLWAWE WRSEVGKQLRPLY
Sbjct  3    ----TIYSTGKACNPSNPQECLLLEPGLDDIMENSKDYNERLWAWEGWRSEVGKQLRPLY  58

Query  181  EEYVVLKNEMARANHYEDYGDYWRGDYEVNGVDGYDYSRGQLIEDVEHTFEEIKPLYEHL  240
            EEYV LKNEMARAN+YEDYGDYWRGDYE    +GY+YSR QLI+DVEHTF +I PLY+HL
Sbjct  59   EEYVALKNEMARANNYEDYGDYWRGDYEEEWENGYNYSRNQLIDDVEHTFTQIMPLYQHL  118

Query  241  HAYVRAKLMNAYPSYISPIGCLPAHLLGDMWGRFWTNLYSLTVPFGQKPNIDVTDAMVDQ  300
            HAYVR KLM+ YPSYISP GCLPAHLLGDMWGRFWTNLY LTVPFGQKPNIDVT+AMV+Q
Sbjct  119  HAYVRTKLMDTYPSYISPTGCLPAHLLGDMWGRFWTNLYPLTVPFGQKPNIDVTNAMVNQ  178

Query  301  AWDAQRIFKEAEKFFVSVGLPNMTQGFWENSMLTDPGNVQKAVCHPTAWDLGKGDFRILM  360
            +WDA++IFKEAEKFFVSVGLPNMTQ FWENSMLT+P + +K VCHPTAWDLGKGDFRI M
Sbjct  179  SWDARKIFKEAEKFFVSVGLPNMTQEFWENSMLTEPSDSRKVVCHPTAWDLGKGDFRIKM  238

Query  361  CTKVTMDDFLTAHHEMGHIQYDMAYAAQPFLLRNGANEGFHEAVGEIMSLSAATPKHLKS  420
            CTKVTMDDFLTAHHEMGHIQYDMAYAAQPFLLRNGANEGFHEAVGEIMSLSAATP HLK+
Sbjct  239  CTKVTMDDFLTAHHEMGHIQYDMAYAAQPFLLRNGANEGFHEAVGEIMSLSAATPNHLKN  298

Query  421  IGLLSPDFQEDNETEINFLLKQALTIVGTLPFTYMLEKWRWMVFKGEIPKDQWMKKWWEM  480
            IGLL P F ED+ETEINFLLKQALTIVGTLPFTYMLEKWRWMVFKGEIPKDQWMK WWEM
Sbjct  299  IGLLPPSFFEDSETEINFLLKQALTIVGTLPFTYMLEKWRWMVFKGEIPKDQWMKTWWEM  358

Query  481  KREIVGVVEPVPHDETYCDPASLFHVSNDYSFIRYYTRTLYQFQFQEALCQAAKHEGPLH  540
            KR IVGVVEPVPHDETYCDPASLFHV+NDYSFIRYYTRT+YQFQFQEALCQ AKHEGPLH
Sbjct  359  KRNIVGVVEPVPHDETYCDPASLFHVANDYSFIRYYTRTIYQFQFQEALCQIAKHEGPLH  418

Query  541  KCDISNSTEAGQKLFNMLRLGKSEPWTLALENVVGAKNMNVRPLLNYFEPLFTWLKDQNK  600
            KCDISNS+EAGQKL  ML+LGKS+PWT ALE VVGAKNM+VRPLLNYFEPLFTWLK+QN+
Sbjct  419  KCDISNSSEAGQKLLEMLKLGKSKPWTYALEIVVGAKNMDVRPLLNYFEPLFTWLKEQNR  478

Query  601  NSFVGWSTDWSPYADQSIKVRISLKSALGDKAYEWNDNEMYLFRSSVAYAMRQYFLKVKN  660
            NSFVGW+TDWSPYADQSIKVRISLKSALG+KAYEWN+NEMYLFRSS+AYAMRQYF +VKN
Sbjct  479  NSFVGWNTDWSPYADQSIKVRISLKSALGEKAYEWNNNEMYLFRSSIAYAMRQYFSEVKN  538

Query  661  QMILFGEEDVRVANLKPRISFNFFVTAPKNVSDIIPRTEVEKAIRMSRSRINDAFRLNDN  720
            Q I F E++V V++LKPRISFNFFVT+P NVSDIIPRTEVE+AIRM RSRIND FRL+DN
Sbjct  539  QTIPFVEDNVWVSDLKPRISFNFFVTSPGNVSDIIPRTEVEEAIRMYRSRINDVFRLDDN  598

Query  721  SLEFLGIQPTLGPPNQPPVSIWLIVFGVVMGVIVVGIVILIFTGIRDRKKKNKARSGENP  780
            SLEFLGIQPTLGPP +PPV+IWLIVFGVVMGV+VVGIV+LIF+GIR+R+K ++AR  ENP
Sbjct  599  SLEFLGIQPTLGPPYEPPVTIWLIVFGVVMGVVVVGIVLLIFSGIRNRRKNDQARGEENP  658

Query  781  YASIDISKGENNPGFQNTDDVQTSF  805
            YAS+D+SKGENNPGFQN DD QTSF
Sbjct  659  YASVDLSKGENNPGFQNVDDAQTSF  683
```

#### angiotensin-converting enzyme 2 isoform X1 [Felis catus]

Sequence ID: XP_023104564.1 Length: 807 Number of Matches: 1

**Human vs Cat**

Range 1: 1 to 807 GenPept Graphics ▼ Next Match

| NW Score | Identities | Positives | Gaps |
|---|---|---|---|
| 3692 | 683/807(85%) | 742/807(91%) | 2/807(0%) |

```
Query  1    MSSSSWLLLSLVAVTAAQSTIEEQAKTFLDKFNHEAEDLFYQSSLASWNYNTNITEENVQ  60
            MS S WLLLS  A+TAAQST EE AKTFL+KFNHEAE+L YQSSLASWNYNTNIT+ENVQ
Sbjct  1    MSGSFWLLLSFAALTAAQSTTEELAKTFLEKFNHEAEELSYQSSLASWNYNTNITDENVQ  60

Query  61   NMNNAGDKWSAFLKEQSTLAQMYPLQEIQNLTVKLQLQALQQNGSSVLSEDKSKRLNTIL  120
             MN AG KWSAF +EQS LA+ YPL EI N TVK QLQALQQ+GSSVLS DKS+RLNTIL
Sbjct  61   KMNEAGAKWSAFYEEQSKLAKTYPLAEIHNTTVKRQLQALQQSGSSVLSADKSQRLNTIL  120

Query  121  NTMSTIYSTGKVCNPDNPQECLLLEPGLNEIMANSLDYNERLWAWESWRSEVGKQLRPLY  180
            N MSTIYSTGK CNP+NPQECLLLEPGL++IM NS DYNERLWAWE WR+EVGKQLRPLY
Sbjct  121  NAMSTIYSTGKACNPNNPQECLLLEPGLDDIMENSKDYNERLWAWEGWRAEVGKQLRPLY  180

Query  181  EEYVVLKNEMARAN--HYEDYGDYWRGDYEVNGVDGYDYSRGQLIEDVEHTFEEIKPLYE  238
            EEYV LKNEMA++   +YEDYGDYWRGDYE    DGY+YSR QLI+DVEHTF +IKPLY+
Sbjct  181  EEYVALKNEMAKSKQVNYEDYGDYWRGDYEEEWTDGYNYSRSQLIKDVEHTFTQIKPLYQ  240

Query  239  HLHAYVRAKLMNAYPSYISPIGCLPAHLLGDMWGRFWTNLYSLTVPFGQKPNIDVTDAMV  298
            HLHAYVRAKLM+ YPS ISP GCLPAHLLGDMWGRFWTNLY LTVPFGQKPNIDVTDAMV
Sbjct  241  HLHAYVRAKLMDTYPSRISPTGCLPAHLLGDMWGRFWTNLYPLTVPFGQKPNIDVTDAMV  300

Query  299  DQAWDAQRIFKEAEKFFVSVGLPNMTQGFWENSMLTDPGNVQKAVCHPTAWDLGKGDFRI  358
            +Q+WDA+RIFKEAEKFFVSVGLPNMTQGFWENSMLT+PG+ +K VCHPTAWDLGKGDFRI
Sbjct  301  NQSWDARRIFKEAEKFFVSVGLPNMTQGFWENSMLTEPGDSRKVVCHPTAWDLGKGDFRI  360

Query  359  LMCTKVTMDDFLTAHHEMGHIQYDMAYAAQPFLLRNGANEGFHEAVGEIMSLSAATPKHL  418
             MCTKVTMDDFLTAHHEMGHIQYDMAYA QPFLLRNGANEGFHEAVGEIMSLSAATP HL
Sbjct  361  KMCTKVTMDDFLTAHHEMGHIQYDMAYAVQPFLLRNGANEGFHEAVGEIMSLSAATPNHL  420

Query  419  KSIGLLSPDFQEDNETEINFLLKQALTIVGTLPFTYMLEKWRWMVFKGEIPKDQWMKKWW  478
            K+IGLLSP F ED+ETEINFLLKQALTIVGTLPFTYMLEKWRWMVFKGEIPK+QWM+KWW
Sbjct  421  KTIGLLSPGFSEDSETEINFLLKQALTIVGTLPFTYMLEKWRWMVFKGEIPKEQWMQKWW  480

Query  479  EMKREIVGVVEPVPHDETYCDPASLFHVSNDYSFIRYYTRTLYQFQFQEALCQAAKHEGP  538
            EMKREIVGVVEPVPHDETYCDPASLFHV+NDYSFIRYYTRT+YQFQFQEALC+ AKHEGP
Sbjct  481  EMKREIVGVVEPVPHDETYCDPASLFHVANDYSFIRYYTRTIYQFQFQEALCRIAKHEGP  540

Query  539  LHKCDISNSTEAGQKLFNMLRLGKSEPWTLALENVVGAKNMNVRPLLNYFEPLFTWLKDQ  598
            LHKCDISNS+EAG+KL  ML LGKS+PWTLALE+VVG K MNV PLL YFEPLFTWLK+Q
Sbjct  541  LHKCDISNSSEAGKKLLQMLTLGKSKPWTLALEHVVGEKKMNVTPLLKYFEPLFTWLKEQ  600

Query  599  NKNSFVGWSTDWSPYADQSIKVRISLKSALGDKAYEWNDNEMYLFRSSVAYAMRQYFLKV  658
            N+NSFVGW+TDW PYADQSIKVRISLKSALGD+AYEWNDNEMYLFRSSVAYAMR+YF KV
Sbjct  601  NRNSFVGWNTDWRPYADQSIKVRISLKSALGDEAYEWNDNEMYLFRSSVAYAMREYFSKV  660

Query  659  KNQMILFGEEDVRVANLKPRISFNFFVTAPKNVSDIIPRTEVEKAIRMSRSRINDAFRLN  718
            KNQ I F E++V V+NLKPRISFNFFVTA KNVSD+IPR+EVE+AIRMSRSRINDAFRL+
Sbjct  661  KNQTIPFVEDNVWVSNLKPRISFNFFVTASKNVSDVIPRSEVEEAIRMSRSRINDAFRLD  720

Query  719  DNSLEFLGIQPTLGPPNQPPVSIWLIVFGVVMGVIVVGIVILIFTGIRDRKKKNKARSGE  778
            DNSLEFLGIQPTL PP QPPV+IWLIVFGVVMGV+VVGIV+LI +GIR+R+K N+ARS E
Sbjct  721  DNSLEFLGIQPTLSPPYQPPVTIWLIVFGVVMGVVVVGIVLLIVSGIRNRRKNNQARSEE  780

Query  779  NPYASIDISKGENNPGFQNTDDVQTSF  805
            NPYAS+D+SKGENNPGFQ+ DDVQTSF
Sbjct  781  NPYASVDLSKGENNPGFQHADDVQTSF  807
```

#### angiotensin-converting enzyme 2 precursor [Felis catus]

Sequence ID: NP_001034545.1 Length: 805 Number of Matches: 1

See 3 more title(s) ⌄

**Human vs Cat**

Range 1: 1 to 805 GenPept Graphics ▼ Next Match

| NW Score | Identities | Positives | Gaps |
|---|---|---|---|
| 3717 | 686/805(85%) | 743/805(92%) | 0/805(0%) |

```
Query  1    MSSSSWLLLSLVAVTAAQSTIEEQAKTFLDKFNHEAEDLFYQSSLASWNYNTNITEENVQ  60
            MS S WLLLS  A+TAAQST EE AKTFL+KFNHEAE+L YQSSLASWNYNTNIT+ENVQ
Sbjct  1    MSGSFWLLLSFAALTAAQSTTEELAKTFLEKFNHEAEELSYQSSLASWNYNTNITDENVQ  60

Query  61   NMNNAGDKWSAFLKEQSTLAQMYPLQEIQNLTVKLQLQALQQNGSSVLSEDKSKRLNTIL  120
             MN AG KWSAF +EQS LA+ YPL EI N TVK QLQALQQ+GSSVLS DKS+RLNTIL
Sbjct  61   KMNEAGAKWSAFYEEQSKLAKTYPLAEIHNTTVKRQLQALQQSGSSVLSADKSQRLNTIL  120

Query  121  NTMSTIYSTGKVCNPDNPQECLLLEPGLNEIMANSLDYNERLWAWESWRSEVGKQLRPLY  180
            N MSTIYSTGK CNP+NPQECLLLEPGL++IM NS DYNERLWAWE WR+EVGKQLRPLY
Sbjct  121  NAMSTIYSTGKACNPNNPQECLLLEPGLDDIMENSKDYNERLWAWEGWRAEVGKQLRPLY  180

Query  181  EEYVVLKNEMARANHYEDYGDYWRGDYEVNGVDGYDYSRGQLIEDVEHTFEEIKPLYEHL  240
            EEYV LKNEMARAN+YEDYGDYWRGDYE    DGY+YSR QLI+DVEHTF +IKPLY+HL
Sbjct  181  EEYVALKNEMARANNYEDYGDYWRGDYEEEWTDGYNYSRSQLIKDVEHTFTQIKPLYQHL  240

Query  241  HAYVRAKLMNAYPSYISPIGCLPAHLLGDMWGRFWTNLYSLTVPFGQKPNIDVTDAMVDQ  300
            HAYVRAKLM+ YPS ISP GCLPAHLLGDMWGRFWTNLY LTVPFGQKPNIDVTDAMV+Q
Sbjct  241  HAYVRAKLMDTYPSRISPTGCLPAHLLGDMWGRFWTNLYPLTVPFGQKPNIDVTDAMVNQ  300

Query  301  AWDAQRIFKEAEKFFVSVGLPNMTQGFWENSMLTDPGNVQKAVCHPTAWDLGKGDFRILM  360
            +WDA+RIFKEAEKFFVSVGLPNMTQGFWENSMLT+PG+ +K VCHPTAWDLGKGDFRI M
Sbjct  301  SWDARRIFKEAEKFFVSVGLPNMTQGFWENSMLTEPGDSRKVVCHPTAWDLGKGDFRIKM  360

Query  361  CTKVTMDDFLTAHHEMGHIQYDMAYAAQPFLLRNGANEGFHEAVGEIMSLSAATPKHLKS  420
            CTKVTMDDFLTAHHEMGHIQYDMAYA QPFLLRNGANEGFHEAVGEIMSLSAATP HLK+
Sbjct  361  CTKVTMDDFLTAHHEMGHIQYDMAYAVQPFLLRNGANEGFHEAVGEIMSLSAATPNHLKT  420

Query  421  IGLLSPDFQEDNETEINFLLKQALTIVGTLPFTYMLEKWRWMVFKGEIPKDQWMKKWWEM  480
            IGLLSP F ED+ETEINFLLKQALTIVGTLPFTYMLEKWRWMVFKGEIPK+QWM+KWWEM
Sbjct  421  IGLLSPGFSEDSETEINFLLKQALTIVGTLPFTYMLEKWRWMVFKGEIPKEQWMQKWWEM  480

Query  481  KREIVGVVEPVPHDETYCDPASLFHVSNDYSFIRYYTRTLYQFQFQEALCQAAKHEGPLH  540
            KREIVGVVEPVPHDETYCDPASLFHV+NDYSFIRYYTRT+YQFQFQEALC+ AKHEGPLH
Sbjct  481  KREIVGVVEPVPHDETYCDPASLFHVANDYSFIRYYTRTIYQFQFQEALCRIAKHEGPLH  540

Query  541  KCDISNSTEAGQKLFNMLRLGKSEPWTLALENVVGAKNMNVRPLLNYFEPLFTWLKDQNK  600
            KCDISNS+EAG+KL  ML LGKS+PWTLALE+VVG K MNV PLL YFEPLFTWLK+QN+
Sbjct  541  KCDISNSSEAGKKLLQMLTLGKSKPWTLALEHVVGEKKMNVTPLLKYFEPLFTWLKEQNR  600

Query  601  NSFVGWSTDWSPYADQSIKVRISLKSALGDKAYEWNDNEMYLFRSSVAYAMRQYFLKVKN  660
            NSFVGW+TDW PYADQSIKVRISLKSALGD+AYEWNDNEMYLFRSSVAYAMR+YF KVKN
Sbjct  601  NSFVGWNTDWRPYADQSIKVRISLKSALGDEAYEWNDNEMYLFRSSVAYAMREYFSKVKN  660

Query  661  QMILFGEEDVRVANLKPRISFNFFVTAPKNVSDIIPRTEVEKAIRMSRSRINDAFRLNDN  720
            Q I F E++V V+NLKPRISFNFFVTA KNVSD+IPR+EVE+AIRMSRSRINDAFRL+DN
Sbjct  661  QTIPFVEDNVWVSNLKPRISFNFFVTASKNVSDVIPRSEVEEAIRMSRSRINDAFRLDDN  720

Query  721  SLEFLGIQPTLGPPNQPPVSIWLIVFGVVMGVIVVGIVILIFTGIRDRKKKNKARSGENP  780
            SLEFLGIQPTL PP QPPV+IWLIVFGVVMGV+VVGIV+LI +GIR+R+K N+ARS ENP
Sbjct  721  SLEFLGIQPTLSPPYQPPVTIWLIVFGVVMGVVVVGIVLLIVSGIRNRRKNNQARSEENP  780

Query  781  YASIDISKGENNPGFQNTDDVQTSF  805
            YAS+D+SKGENNPGFQ+ DDVQTSF
```

**PREDICTED: angiotensin-converting enzyme 2, partial [Panthera tigris altaica]**

Sequence ID: XP_007090142.1 Length: 797 Number of Matches: 1

#### Human vs Tiger

Range 1: 1 to 797 GenPept Graphics ▼ Next Match

| Score | Expect | Method | Identities | Positives | Gaps |
|---|---|---|---|---|---|
| 1415 bits(3662) | 0.0 | Compositional matrix adjust. | 683/797(86%) | 737/797(92%) | 0/797(0[illegible] |

```
Query  9    LSLVAVTAAQSTIEEQAKTFLDKFNHEAEDLFYQSSLASWNYNTNITEENVQNMNNAGDK  68
            LS  A+TAAQST EE AKTFL+KFNHEAE+L YQSSLASWNYNTNIT+ENVQ MN AG K
Sbjct  1    LSFAALTAAQSTTEELAKTFLEKFNHEAEELSYQSSLASWNYNTNITDENVQKMNEAGAK  60

Query  69   WSAFLKEQSTLAQMYPLQEIQNLTVKLQLQALQQNGSSVLSEDKSKRLNTILNTMSTIYS  128
            WSAF +EQS LA+ YPL EI N TVK QLQALQQ+GSSVLS DKS+RLNTILN MSTIYS
Sbjct  61   WSAFYEEQSKLAETYPLAEIHNTTVKRQLQALQQSGSSVLSADKSQRLNTILNAMSTIYS  120

Query  129  TGKVCNPDNPQECLLLEPGLNEIMANSLDYNERLWAWESWRSEVGKQLRPLYEEYVVLKN  188
            TGK CNP+NPQECLLLEPGL++IM NS DYNERLWAWE WR+EVGKQLRPLYEEYV LKN
Sbjct  121  TGKACNPNNPQECLLLEPGLDDIMENSKDYNERLWAWEGWRAEVGKQLRPLYEEYVALKN  180

Query  189  EMARANHYEDYGDYWRGDYEVNGVDGYDYSRGQLIEDVEHTFEEIKPLYEHLHAYVRAKL  248
            EMARAN+YEDYGDYWRGDYE    DGY+YSR QLI+DVEHTF +IKPLY+HLHAYVRAKL
Sbjct  181  EMARANNYEDYGDYWRGDYEEEWTDGYNYSRSQLIKDVEHTFTQIKPLYQHLHAYVRAKL  240

Query  249  MNAYPSYISPIGCLPAHLLGDMWGRFWTNLYSLTVPFGQKPNIDVTDAMVDQAWDAQRIF  308
            M++YPS ISP GCLPAHLLGDMWGRFWTNLY LTVPFGQKPNIDVTDAMV+Q+WDA+RIF
Sbjct  241  MDSYPSRISPTGCLPAHLLGDMWGRFWTNLYPLTVPFGQKPNIDVTDAMVNQSWDARRIF  300

Query  309  KEAEKFFVSVGLPNMTQGFWENSMLTDPGNVQKAVCHPTAWDLGKGDFRILMCTKVTMDD  368
            KEAEKFFVSVGLPNMTQGFWENSMLT+PGN QK VCHPTAWDLGKGDFRI MCTKVTMDD
Sbjct  301  KEAEKFFVSVGLPNMTQGFWENSMLTEPGNSQKVVCHPTAWDLGKGDFRIKMCTKVTMDD  360

Query  369  FLTAHHEMGHIQYDMAYAAQPFLLRNGANEGFHEAVGEIMSLSAATPKHLKSIGLLSPDF  428
            FLTAHHEMGHIQYDMAYA QPFLLRNGANEGFHEAVGEIMSLSAATP HLK+IGLL P F
Sbjct  361  FLTAHHEMGHIQYDMAYAVQPFLLRNGANEGFHEAVGEIMSLSAATPNHLKTIGLLPPGF  420

Query  429  QEDNETEINFLLKQALTIVGTLPFTYMLEKWRWMVFKGEIPKDQWMKKWWEMKREIVGVV  488
             ED+ETEINFLLKQALTIVGTLPFTYMLEKWRWMVFKGEIPK+QWM+KWWEMKREIVGVV
Sbjct  421  SEDSETEINFLLKQALTIVGTLPFTYMLEKWRWMVFKGEIPKEQWMQKWWEMKREIVGVV  480

Query  489  EPVPHDETYCDPASLFHVSNDYSFIRYYTRTLYQFQFQEALCQAAKHEGPLHKCDISNST  548
            EPVPHDETYCDPASLFHV+NDYSFIRYYTRT+YQFQFQEALC+ AKHEGPLHKCDISNS+
Sbjct  481  EPVPHDETYCDPASLFHVANDYSFIRYYTRTIYQFQFQEALCRIAKHEGPLHKCDISNSS  540

Query  549  EAGQKLFNMLRLGKSEPWTLALENVVGAKNMNVRPLLNYFEPLFTWLKDQNKNSFVGWST  608
            EAG+KL  ML LGKS+PWTLALE+VVG KNMNV PLL YFEPLFTWLK+QN+NSFVGW+T
Sbjct  541  EAGKKLLQMLTLGKSKPWTLALEHVVGEKNMNVTPLLKYFEPLFTWLKEQNRNSFVGWNT  600

Query  609  DWSPYADQSIKVRISLKSALGDKAYEWNDNEMYLFRSSVAYAMRQYFLKVKNQMILFGEE  668
            DW PYADQSIKVRISLKSALGDKAYEWNDNEMYLFRSSVAYAMR+YF KVKNQ I F E+
Sbjct  601  DWRPYADQSIKVRISLKSALGDKAYEWNDNEMYLFRSSVAYAMREYFSKVKNQTIPFVED  660

Query  669  DVRVANLKPRISFNFFVTAPKNVSDIIPRTEVEKAIRMSRSRINDAFRLNDNSLEFLGIQ  728
            +V V+NLKPRISFNFFVTA KNVSD+IPR EVE+AIRMSRSRINDAFRL+DNSLEFLGIQ
Sbjct  661  NVWVSNLKPRISFNFFVTASKNVSDVIPRREVEEAIRMSRSRINDAFRLDDNSLEFLGIQ  720

Query  729  PTLGPPNQPPVSIWLIVFGVVMGVIVVGIVILIFTGIRDRKKKNKARSGENPYASIDISK  788
            PTL PP QPPV+IWLIVFGVVMGV+VVGIV+LI +GIR+R+K N+ARS ENPYAS+D+SK
Sbjct  721  PTLSPPYQPPVTIWLIVFGVVMGVVVVGIVLLIVSGIRNRRKNNQARSEENPYASVDLSK  780

Query  789  GENNPGFQNTDDVQTSF  805
            GENNPGFQ+ DDVQTSF
Sbjct  781  GENNPGFQHADDVQTSF  797
```

**PREDICTED: angiotensin-converting enzyme 2 isoform X1 [Myotis brandtii]**

Sequence ID: XP_014399780.1 Length: 819 Number of Matches: 1

See 2 more title(s)

#### Human vs Bat

Range 1: 1 to 819 GenPept Graphics Next Match

| NW Score | Identities | Positives | Gaps |
|---|---|---|---|
| 3529 | 651/820(79%) | 724/820(88%) | 16/820(1%) |

```
Query  1    MSSSSWLLLSLVAVTAAQSTIEEQAKTFLDKFNHEAEDLFYQSSLASWNYNTNITEENVQ  60
            MS SSWL LSLVAV AAQS+ EE+AK FL+ FN +AEDL ++S+LASWNYNTNIT+ENVQ
Sbjct  1    MSGSSWLFLSLVAVAAAQSSTEEKAKIFLENFNSKAEDLSHESALASWNYNTNITDENVQ  60

Query  61   NMNNAGDKWSAFLKEQSTLAQMYPLQEIQNLTVKLQLQALQQNGSSVLSEDKSKRLNTIL  120
             MN A   KWSAF ++QS LAQ YPLQEIQNLT+K QLQ LQQNGSSVLS DKSKRLNTIL
Sbjct  61   KMNEADSKWSAFYEQQSKLAQTYPLQEIQNLTIKRQLQVLQQNGSSVLSADKSKRLNTIL  120

Query  121  NTMSTIYSTGKVCNPDNPQECLLLEPGLNEIMANSLDYNERLWAWESWRSEVGKQLRPLY  180
             TMSTIYSTGKVCNP+NPQEC  L  GL +IM  S DYN+RLW WE WRSEVGKQLRPLY
Sbjct  121  TTMSTIYSTGKVCNPNNPQECFTLA-GLEDIMEKSKDYNQRLWVWEGWRSEVGKQLRPLY  179

Query  181  EEYVVLKNEMARANHYEDYGDYWRGDYEVNGVDGYDYSRGQLIEDVEHTFEEIKPLYEHL  240
            EEYV LKNEMAR N+YEDYGDYWRGDYE  G DGY+YSR QL EDVE  F EIKPLYEHL
Sbjct  180  EEYVDLKNEMARGNNYEDYGDYWRGDYETEGEDGYNYSRNQLTEDVERIFLEIKPLYEHL  239

Query  241  HAYVRAKLMNAYPSYISPIGCLPAHLLGDMWGRFWTNLYSLTVPFGQKPNIDVTDAMVDQ  300
            HAYVRAKL+NAYPS ISP G LPAHLLGDMWGRFWTNLY+LTVPF QKPNIDVT AMV+Q
Sbjct  240  HAYVRAKLVNAYPSRISPTGYLPAHLLGDMWGRFWTNLYNLTVPFEQKPNIDVTGAMVEQ  299

Query  301  AWDAQRIFKEAEKFFVSVGLPNMTQGFWENSMLTDPGNVQKAVCHPTAWDLGKGDFRILM  360
            +WDA++IFKEAEKF++SVGLP+MT GFW NSMLT+PG+ +K VCHPTAWDLGKGDFRI M
Sbjct  300  SWDAEKIFKEAEKFYISVGLPSMTPGFWNNSMLTEPGDGRKVVCHPTAWDLGKGDFRIKM  359

Query  361  CTKVTMDDFLTAHHEMGHIQYDMAYAAQPFLLRNGANEGFHEAVGEIMSLSAATPKHLKS  420
            CTKVTMDDFLTAHHEMGHIQYDMAYA QP+LLRNGANEGFHEAVGE+MSLS ATPKHLK
Sbjct  360  CTKVTMDDFLTAHHEMGHIQYDMAYATQPYLLRNGANEGFHEAVGEVMSLSVATPKHLKV  419

Query  421  IGLLSPDFQEDNETEINFLLKQALTIVGTLPFTYMLEKWRWMVFKGEIPKDQWMKKWWEM  480
            +GLL PDF EDNETEINFLLKQAL IVGTLPFTYMLEKWRWMVFKGEIPK+QWMKKWWEM
Sbjct  420  MGLLPPDFSEDNETEINFLLKQALNIVGTLPFTYMLEKWRWMVFKGEIPKEQWMKKWWEM  479

Query  481  KREIVGVVEPVPHDETYCDPASLFHVSNDYSFIRYYTRTLYQFQFQEALCQAAKHEGPLH  540
            KREIVGV+EP+PHDETYCDPASLFHV+NDYSFIRY+TRT+++FQFQEALCQ AKH+GPLH
Sbjct  480  KREIVGVMEPLPHDETYCDPASLFHVANDYSFIRYFTRTIFEFQFQEALCQIAKHQGPLH  539

Query  541  KCDISNSTEAGQKLFNMLRLGKSEPWTLALENVVGAKNMNVRPLLNYFEPLFTWLKDQNK  600
            KCDISNS EAG KL  ML+LGKSEPWTLALE +VG K M+ +PLLNYFEPLFTWLK+QN
Sbjct  540  KCDISNSKEAGNKLLEMLKLGKSEPWTLALEKIVGTKKMDAKPLLNYFEPLFTWLKEQNG  599

Query  601  NSF---------------VGWSTDWSPYADQSIKVRISLKSALGDKAYEWNDNEMYLFRS  645
            NS                VGW +DWSPYA+QSIKVRISLKSALG+KAY+WN+NEMYLF+S
Sbjct  600  NSVGWNSGNSVESRSGNSVGWFSDWSPYAEQSIKVRISLKSALGEKAYKWNENEMYLFQS  659

Query  646  SVAYAMRQYFLKVKNQMILFGEEDVRVANLKPRISFNFFVTAPKNVSDIIPRTEVEKAIR  705
            SVAYAMR+YFLK KNQ I FG E+V V ++KPRISF FFVT+P+N+S +IPR+EVE AIR
Sbjct  660  SVAYAMREYFLKEKNQTIPFGVENVMVNDVKPRISFKFFVTSPENISVVIPRSEVEDAIR  719

Query  706  MSRSRINDAFRLNDNSLEFLGIQPTLGPPNQPPVSIWLIVFGVVMGVIVVGIVILIFTGI  765
            MSRSRINDAFRL+DN+LEFLGIQPTLGPPNQPPV+IWLIVFGVVMGV+V+GI +LIFTGI
Sbjct  720  MSRSRINDAFRLDDNTLEFLGIQPTLGPPNQPPVTIWLIVFGVVMGVVVIGIAVLIFTGI  779

Query  766  RDRKKKNKARSGENPYASIDISKGENNPGFQNTDDVQTSF  805
            RDRKKK +A + ENPY+S+++SKGENNPGFQ+ DDVQTSF
Sbjct  780  RDRKKKQAGNEENPYSSVNLSKGENNPGFQSGDDVQTSF  819
```


**PREDICTED: angiotensin-converting enzyme 2 isoform X1 [Myotis brandtii]**

Sequence ID: XP_014399780.1 Length: 819 Number of Matches: 1

See 2 more title(s)

#### Human vs Bat

Range 1: 1 to 819 GenPept Graphics Next Match

| NW Score | Identities | Positives | Gaps |
|---|---|---|---|
| 3529 | 651/820(79%) | 724/820(88%) | 16/820(1%) |

```
Query  1    MSSSSWLLLSLVAVTAAQSTIEEQAKTFLDKFNHEAEDLFYQSSLASWNYNTNITEENVQ  60
            MS SSWL LSLVAV AAQS+ EE+AK FL+ FN +AEDL ++S+LASWNYNTNIT+ENVQ
Sbjct  1    MSGSSWLFLSLVAVAAAQSSTEEKAKIFLENFNSKAEDLSHESALASWNYNTNITDENVQ  60

Query  61   NMNNAGDKWSAFLKEQSTLAQMYPLQEIQNLTVKLQLQALQQNGSSVLSEDKSKRLNTIL  120
             MN A   KWSAF ++QS LAQ YPLQEIQNLT+K QLQ LQQNGSSVLS DKSKRLNTIL
Sbjct  61   KMNEADSKWSAFYEQQSKLAQTYPLQEIQNLTIKRQLQVLQQNGSSVLSADKSKRLNTIL  120

Query  121  NTMSTIYSTGKVCNPDNPQECLLLEPGLNEIMANSLDYNERLWAWESWRSEVGKQLRPLY  180
             TMSTIYSTGKVCNP+NPQEC  L  GL +IM  S DYN+RLW WE WRSEVGKQLRPLY
Sbjct  121  TTMSTIYSTGKVCNPNNPQECFTLA-GLEDIMEKSKDYNQRLWVWEGWRSEVGKQLRPLY  179

Query  181  EEYVVLKNEMARANHYEDYGDYWRGDYEVNGVDGYDYSRGQLIEDVEHTFEEIKPLYEHL  240
            EEYV LKNEMAR N+YEDYGDYWRGDYE  G DGY+YSR QL EDVE  F EIKPLYEHL
Sbjct  180  EEYVDLKNEMARGNNYEDYGDYWRGDYETEGEDGYNYSRNQLTEDVERIFLEIKPLYEHL  239

Query  241  HAYVRAKLMNAYPSYISPIGCLPAHLLGDMWGRFWTNLYSLTVPFGQKPNIDVTDAMVDQ  300
            HAYVRAKL+NAYPS ISP G LPAHLLGDMWGRFWTNLY+LTVPF QKPNIDVT AMV+Q
Sbjct  240  HAYVRAKLVNAYPSRISPTGYLPAHLLGDMWGRFWTNLYNLTVPFEQKPNIDVTGAMVEQ  299

Query  301  AWDAQRIFKEAEKFFVSVGLPNMTQGFWENSMLTDPGNVQKAVCHPTAWDLGKGDFRILM  360
            +WDA++IFKEAEKF++SVGLP+MT GFW NSMLT+PG+ +K VCHPTAWDLGKGDFRI M
Sbjct  300  SWDAEKIFKEAEKFYISVGLPSMTPGFWNNSMLTEPGDGRKVVCHPTAWDLGKGDFRIKM  359

Query  361  CTKVTMDDFLTAHHEMGHIQYDMAYAAQPFLLRNGANEGFHEAVGEIMSLSAATPKHLKS  420
            CTKVTMDDFLTAHHEMGHIQYDMAYA QP+LLRNGANEGFHEAVGE+MSLS ATPKHLK
Sbjct  360  CTKVTMDDFLTAHHEMGHIQYDMAYATQPYLLRNGANEGFHEAVGEVMSLSVATPKHLKV  419

Query  421  IGLLSPDFQEDNETEINFLLKQALTIVGTLPFTYMLEKWRWMVFKGEIPKDQWMKKWWEM  480
            +GLL PDF EDNETEINFLLKQAL IVGTLPFTYMLEKWRWMVFKGEIPK+QWMKKWWEM
Sbjct  420  MGLLPPDFSEDNETEINFLLKQALNIVGTLPFTYMLEKWRWMVFKGEIPKEQWMKKWWEM  479

Query  481  KREIVGVVEPVPHDETYCDPASLFHVSNDYSFIRYYTRTLYQFQFQEALCQAAKHEGPLH  540
            KREIVGV+EP+PHDETYCDPASLFHV+NDYSFIRY+TRT+++FQFQEALCQ AKH+GPLH
Sbjct  480  KREIVGVMEPLPHDETYCDPASLFHVANDYSFIRYFTRTIFEFQFQEALCQIAKHQGPLH  539

Query  541  KCDISNSTEAGQKLFNMLRLGKSEPWTLALENVVGAKNMNVRPLLNYFEPLFTWLKDQNK  600
            KCDISNS EAG KL  ML+LGKSEPWTLALE +VG K M+ +PLLNYFEPLFTWLK+QN
Sbjct  540  KCDISNSKEAGNKLLEMLKLGKSEPWTLALEKIVGTKKMDAKPLLNYFEPLFTWLKEQNG  599

Query  601  NSF---------------VGWSTDWSPYADQSIKVRISLKSALGDKAYEWNDNEMYLFRS  645
            NS                VGW +DWSPYA+QSIKVRISLKSALG+KAY+WN+NEMYLF+S
Sbjct  600  NSVGWNSGNSVESRSGNSVGWFSDWSPYAEQSIKVRISLKSALGEKAYKWNENEMYLFQS  659

Query  646  SVAYAMRQYFLKVKNQMILFGEEDVRVANLKPRISFNFFVTAPKNVSDIIPRTEVEKAIR  705
            SVAYAMR+YFLK KNQ I FG E+V V ++KPRISF FFVT+P+N+S +IPR+EVE AIR
Sbjct  660  SVAYAMREYFLKEKNQTIPFGVENVMVNDVKPRISFKFFVTSPENISVVIPRSEVEDAIR  719

Query  706  MSRSRINDAFRLNDNSLEFLGIQPTLGPPNQPPVSIWLIVFGVVMGVIVVGIVILIFTGI  765
            MSRSRINDAFRL+DN+LEFLGIQPTLGPPNQPPV+IWLIVFGVVMGV+V+GI +LIFTGI
Sbjct  720  MSRSRINDAFRLDDNTLEFLGIQPTLGPPNQPPVTIWLIVFGVVMGVVVIGIAVLIFTGI  779

Query  766  RDRKKKNKARSGENPYASIDISKGENNPGFQNTDDVQTSF  805
            RDRKKK +A + ENPY+S+++SKGENNPGFQ+ DDVQTSF
Sbjct  780  RDRKKKQAGNEENPYSSVNLSKGENNPGFQSGDDVQTSF  819
```

**PREDICTED: angiotensin-converting enzyme 2 isoform X2 [Myotis brandtii]**

Sequence ID: XP_014399782.1 Length: 799 Number of Matches: 1

**Human vs Bat**

Range 1: 1 to 799 GenPept Graphics ▼ Next Match

| NW Score | Identities | Positives | Gaps |
| --- | --- | --- | --- |
| 3503 | 647/805(80%) | 721/805(89%) | 6/805(0%) |

```
Query  1    MSSSSWLLLSLVAVTAAQSTIEEQAKTFLDKFNHEAEDLFYQSSLASWNYNTNITEENVQ  60
            MS SSWL LSLVAV AAQS+ EE+AK FL+ FN +AEDL ++S+LASWNYNTNIT+ENVQ
Sbjct  1    MSGSSWLFLSLVAVAAAQSSTEEKAKIFLENFNSKAEDLSHESALASWNYNTNITDENVQ  60

Query  61   NMNNAGDKWSAFLKEQSTLAQMYPLQEIQNLTVKLQLQALQQNGSSVLSEDKSKRLNTIL  120
             MN A  KWSAF ++QS LAQ YPLQEIQNLT+K QLQ LQQNGSSVLS DKSKRLNTIL
Sbjct  61   KMNEADSKWSAFYEQQSKLAQTYPLQEIQNLTIKRQLQVLQQNGSSVLSADKSKRLNTIL  120

Query  121  NTMSTIYSTGKVCNPDNPQECLLLEPGLNEIMANSLDYNERLWAWESWRSEVGKQLRPLY  180
             TMSTIYSTGKVCNP+NPQEC  L  GL +IM  S DYN+RLW WE WRSEVGKQLRPLY
Sbjct  121  TTMSTIYSTGKVCNPNNPQECFTLA-GLEDIMEKSKDYNQRLWVWEGWRSEVGKQLRPLY  179

Query  181  EEYVVLKNEMARANHYEDYGDYWRGDYEVNGVDGYDYSRGQLIEDVEHTFEEIKPLYEHL  240
            EEYV LKNEMAR N+YEDYGDYWRGDYE  G DGY+YSR QL EDVE  F EIKPLYEHL
Sbjct  180  EEYVDLKNEMARGNNYEDYGDYWRGDYETEGEDGYNYSRNQLTEDVERIFLEIKPLYEHL  239

Query  241  HAYVRAKLMNAYPSYISPIGCLPAHLLGDMWGRFWTNLYSLTVPFGQKPNIDVTDAMVDQ  300
            HAYVRAKL+NAYPS ISP G LPAHLLGDMWGRFWTNLY+LTVPF QKPNIDVT AMV+Q
Sbjct  240  HAYVRAKLVNAYPSRISPTGYLPAHLLGDMWGRFWTNLYNLTVPFEQKPNIDVTGAMVEQ  299

Query  301  AWDAQRIFKEAEKFFVSVGLPNMTQGFWENSMLTDPGNVQKAVCHPTAWDLGKGDFRILM  360
            +WDA++IFKEAEKF++SVGLP+MT GFW NSMLT+PG+ +K VCHPTAWDLGKGDFRI M
Sbjct  300  SWDAEKIFKEAEKFYISVGLPSMTPGFWNNSMLTEPGDGRKVVCHPTAWDLGKGDFRIKM  359

Query  361  CTKVTMDDFLTAHHEMGHIQYDMAYAAQPFLLRNGANEGFHEAVGEIMSLSAATPKHLKS  420
            CTKVTMDDFLTAHHEMGHIQYDMAYA QP+LLRNGANEGFHEAVGE+MSLS ATPKHLK
Sbjct  360  CTKVTMDDFLTAHHEMGHIQYDMAYATQPYLLRNGANEGFHEAVGEVMSLSVATPKHLKV  419

Query  421  IGLLSPDFQEDNETEINFLLKQALTIVGTLPFTYMLEKWRWMVFKGEIPKDQWMKKWWEM  480
            +GLL PDF EDNETEINFLLKQAL IVGTLPFTYMLEKWRWMVFKGEIPK+QWMKKWWEM
Sbjct  420  MGLLPPDFSEDNETEINFLLKQALNIVGTLPFTYMLEKWRWMVFKGEIPKEQWMKKWWEM  479

Query  481  KREIVGVVEPVPHDETYCDPASLFHVSNDYSFIRYYTRTLYQFQFQEALCQAAKHEGPLH  540
            KREIVGV+EP+PHDETYCDPASLFHV+NDYSFIRY+TRT+++FQFQEALCQ AKH+GPLH
Sbjct  480  KREIVGVMEPLPHDETYCDPASLFHVANDYSFIRYFTRTIFEFQFQEALCQIAKHQGPLH  539

Query  541  KCDISNSTEAGQKLFNMLRLGKSEPWTLALENVVGAKNMNVRPLLNYFEPLFTWLKDQNK  600
            KCDISNS EAG KL  ML+LGKSEPWTLALE +VG K M+ +PLLNYFEPLFTWLK+QN
Sbjct  540  KCDISNSKEAGNKLLEMLKLGKSEPWTLALEKIVGTKKMDAKPLLNYFEPLFTWLKEQNG  599

Query  601  NSFVGWSTDWSPYADQSIKVRISLKSALGDKAYEWNDNEMYLFRSSVAYAMRQYFLKVKN  660
            NS VGW++D    A+QSIKVRISLKSALG+KAY+WN+NEMYLF+SSVAYAMR+YFLK KN
Sbjct  600  NS-VGWNSD----AEQSIKVRISLKSALGEKAYKWNENEMYLFQSSVAYAMREYFLKEKN  654

Query  661  QMILFGEEDVRVANLKPRISFNFFVTAPKNVSDIIPRTEVEKAIRMSRSRINDAFRLNDN  720
            Q I FG E+V V ++KPRISF FFVT+P+N+S +IPR+EVE AIRMSRSRINDAFRL+DN
Sbjct  655  QTIPFGVENVMVNDVKPRISFKFFVTSPENISVVIPRSEVEDAIRMSRSRINDAFRLDDN  714

Query  721  SLEFLGIQPTLGPPNQPPVSIWLIVFGVVMGVIVVGIVILIFTGIRDRKKKNKARSGENP  780
            +LEFLGIQPTLGPPNQPPV+IWLIVFGVVMGV+V+GI +LIFTGIRDRKKK +A + ENP
Sbjct  715  TLEFLGIQPTLGPPNQPPVTIWLIVFGVVMGVVVIGIAVLIFTGIRDRKKKKQAGNEENP  774

Query  781  YASIDISKGENNPGFQNTDDVQTSF  805
            Y+S+++SKGENNPGFQ+ DDVQTSF
Sbjct  775  YSSVNLSKGENNPGFQSGDDVQTSF  799
```

**PREDICTED: angiotensin-converting enzyme 2 isoform X3 [Myotis brandtii]**

Sequence ID: XP_014399783.1 Length: 754 Number of Matches: 1

**Human vs Bat**

Range 1: 1 to 754 GenPept Graphics ▼ Next Matc

| NW Score | Identities | Positives | Gaps |
| --- | --- | --- | --- |
| 3141 | 593/820(72%) | 660/820(80%) | 81/820(9%) |

```
Query  1    MSSSSWLLLSLVAVTAAQSTIEEQAKTFLDKFNHEAEDLFYQSSLASWNYNTNITEENVQ  60
            MS SSWL LSLVAV AAQS+ EE+AK FL+ FN +AEDL ++S+LASWNYNTNIT+ENVQ
Sbjct  1    MSGSSWLFLSLVAVAAAQSSTEEKAKIFLENFNSKAEDLSHESALASWNYNTNITDENVQ  60

Query  61   NMNNAGDKWSAFLKEQSTLAQMYPLQEIQNLTVKLQLQALQQNGSSVLSEDKSKRLNTIL  120
             MN A  KWSAF ++QS LAQ YPLQEIQNLT+K QLQ LQQNGSSVLS DKSKRLNTIL
Sbjct  61   KMNEADSKWSAFYEQQSKLAQTYPLQEIQNLTIKRQLQVLQQNGSSVLSADKSKRLNTIL  120

Query  121  NTMSTIYSTGKVCNPDNPQECLLLEPGLNEIMANSLDYNERLWAWESWRSEVGKQLRPLY  180
             TMSTIYSTGKVCNP+NPQEC  L  GL +IM  S DYN+RLW WE WRSEVGKQLRPLY
Sbjct  121  TTMSTIYSTGKVCNPNNPQECFTLA-GLEDIMEKSKDYNQRLWVWEGWRSEVGKQLRPLY  179

Query  181  EEYVVLKNEMARANHYEDYGDYWRGDYEVNGVDGYDYSRGQLIEDVEHTFEEIKPLYEHL  240
            EEYV LKNEMAR N+YEDYGDYWRGDYE  G DGY+YSR QL EDVE  F EIKPLYEHL
Sbjct  180  EEYVDLKNEMARGNNYEDYGDYWRGDYETEGEDGYNYSRNQLTEDVERIFLEIKPLYEHL  239

Query  241  HAYVRAKLMNAYPSYISPIGCLPAHLLGDMWGRFWTNLYSLTVPFGQKPNIDVTDAMVDQ  300
            HAYVRAKL+NAYPS ISP G LPAHLLGDMWGRFWTNLY+LTVPF QKPNIDVT AMV+Q
Sbjct  240  HAYVRAKLVNAYPSRISPTGYLPAHLLGDMWGRFWTNLYNLTVPFEQKPNIDVTGAMVEQ  299

Query  301  AWDAQRIFKEAEKFFVSVGLPNMTQGFWENSMLTDPGNVQKAVCHPTAWDLGKGDFRILM  360
            +WDA++IFKEAEKF++SVGLP+MT GFW NSMLT+PG+ +K VCHPTAWDLGKGDFRI M
Sbjct  300  SWDAEKIFKEAEKFYISVGLPSMTPGFWNNSMLTEPGDGRKVVCHPTAWDLGKGDFRIKM  359

Query  361  CTKVTMDDFLTAHHEMGHIQYDMAYAAQPFLLRNGANEGFHEAVGEIMSLSAATPKHLKS  420
            CTKVTMDDFLTAHHEMGHIQYDMAYA QP+LLRNGANEGFHEAVGE+MSLS ATPKHLK
Sbjct  360  CTKVTMDDFLTAHHEMGHIQYDMAYATQPYLLRNGANEGFHEAVGEVMSLSVATPKHLKV  419

Query  421  IGLLSPDFQEDNETEINFLLKQALTIVGTLPFTYMLEKWRWMVFKGEIPKDQWMKKWWEM  480
            +GLL PDF EDNETEINFLLKQAL IVGTLPFTYMLEKWRWMVFKGEIPK+QWMKKWWEM
Sbjct  420  MGLLPPDFSEDNETEINFLLKQALNIVGTLPFTYMLEKWRWMVFKGEIPKEQWMKKWWEM  479

Query  481  KREIVGVVEPVPHDETYCDPASLFHVSNDYSFIRYYTRTLYQFQFQEALCQAAKHEGPLH  540
            KREIVGV+EP+PHDETYCDPASLFHV+NDYSFIRY+TRT+++FQFQEALCQ AKH+GPLH
Sbjct  480  KREIVGVMEPLPHDETYCDPASLFHVANDYSFIRYFTRTIFEFQFQEALCQIAKHQGPLH  539

Query  541  KCDISNSTEAGQKLFNMLRLGKSEPWTLALENVVGAKNMNVRPLLNYFEPLFTWLKDQNK  600
            KCDISNS EAG KL  ML+LGKSEPWTLALE +VG K M+ +PLLNYFEPLFTWLK+QN
Sbjct  540  KCDISNSKEAGNKLLEMLKLGKSEPWTLALEKIVGTKKMDAKPLLNYFEPLFTWLKEQNG  599

Query  601  NSF---------------VGWSTDWSPYADQSIKVRISLKSALGDKAYEWNDNEMYLFRS  645
            NS                VGW +DWSPYA+QSIKVRISLKSALG+KAY+WN+NEMYLF+S
Sbjct  600  NSVGWNSGNSVESRSGNSVGWFSDWSPYAEQSIKVRISLKSALGEKAYKWNENEMYLFQS  659

Query  646  SVAYAMRQYFLKVKNQMILFGEEDVRVANLKPRISFNFFVTAPKNVSDIIPRTEVEKAIR  705
            SVAYAMR+YFLK KNQ I FG E+V V ++KPRISF FFVT+P+N+S +IPR+EVE AIR
Sbjct  660  SVAYAMREYFLKEKNQTIPFGVENVMVNDVKPRISFKFFVTSPENISVVIPRSEVEDAIR  719

Query  706  MSRSRINDAFRLNDNSLEFLGIQPTLGPPNQPPVSIWLIVFGVVMGVIVVGIVILIFTGI  765

Sbjct       ------------------------------------------------------------

Query  766  RDRKKKNKARSGENPYASIDISKGENNPGFQNTDDVQTSF  805
                 K +A + ENPY+S+++SKGENNPGFQ+ DDVQTSF
Sbjct  720  -----KKQAGNEENPYSSVNLSKGENNPGFQSGDDVQTSF  754
```

**angiotensin-converting enzyme 2 isoform X1 [Desmodus rotundus]**

Sequence ID: XP_024425698.1 Length: 804 Number of Matches: 1

**Human vs Bat**

Range 1: 1 to 804 GenPept Graphics ▼ Next Match

| NW Score | Identities | Positives | Gaps |
|---|---|---|---|
| 3465 | 641/807(79%) | 711/807(88%) | 5/807(0%) |

```
Query  1    MSSSSWLLLSLVAVTAAQSTIEEQAKTFLDKFNHEAEDLFYQSSLASWNYNTNITEENVQ  60
            MS SSWL LSLVAV AAQ+  EE+A+TFL+ FN EAE+ FYQ+SLASWNYNTNIT+ENVQ
Sbjct  1    MSGSSWLFLSLVAVAAAQTPTEEEARTFLENFNTEAEEWFYQNSLASWNYNTNITDENVQ  60

Query  61   NMNNAGDKWSAFLKEQSTLAQMYPLQEIQNLTVKLQLQALQQNGSSVLSEDKSKRL--NT  118
             MN A   WS F +  S +A+ YPL+ I+++ VK QLQALQQNG   L EDK K+L  N
Sbjct  61   KMNEAEQMWSTFYERNSNIAKTYPLETIKDVNVKRQLQALQQNG---LLEDKDKQLQLNA  117

Query  119  ILNTMSTIYSTGKVCNPDNPQECLLLEPGLNEIMANSLDYNERLWAWESWRSEVGKQLRP  178
            ILNTMSTIYSTGKVC P+NPQEC LL   GL +IM +S DYNERLWAWE WRS+VGKQLRP
Sbjct  118  ILNTMSTIYSTGKVCKPNNPQECYLLATGLEDIMQDSKDYNERLWAWEGWRSKVGKQLRP  177

Query  179  LYEEYVVLKNEMARANHYEDYGDYWRGDYEVNGVDGYDYSRGQLIEDVEHTFEEIKPLYE  238
            LYEEYVVLKNEMAR  +YEDYGDYWRGDYE  G  GY+YSR QLIEDVE+TF EIKPLYE
Sbjct  178  LYEEYVVLKNEMAREKNYEDYGDYWRGDYETEGSSGYEYSRNQLIEDVENTFAEIKPLYE  237

Query  239  HLHAYVRAKLMNAYPSYISPIGCLPAHLLGDMWGRFWTNLYSLTVPFGQKPNIDVTDAMV  298
            HLHAYVRAKLM+ YPS+ISP GCLPAHLLGDMWGRFWTNLY+LT PFG+KP IDVT AMV
Sbjct  238  HLHAYVRAKLMDTYPSHISPTGCLPAHLLGDMWGRFWTNLYNLTAPFGEKPTIDVTAAMV  297

Query  299  DQAWDAQRIFKEAEKFFVSVGLPNMTQGFWENSMLTDPGNVQKAVCHPTAWDLGKGDFRI  358
            DQ+WDAQRIFKEAEKFF SVGL +MTQGFW+NSMLT P + ++ VCHPTAWDLG  DFRI
Sbjct  298  DQSWDAQRIFKEAEKFFKSVGLFSMTQGFWDNSMLTKPDDGREVVCHPTAWDLGNKDFRI  357

Query  359  LMCTKVTMDDFLTAHHEMGHIQYDMAYAAQPFLLRNGANEGFHEAVGEIMSLSAATPKHL  418
             MCTKVTMDDFLTAHHEMGHIQYDMAYA Q FLLRNGANEGFHEAVGEIMSLS ATPKHL
Sbjct  358  KMCTKVTMDDFLTAHHEMGHIQYDMAYANQSFLLRNGANEGFHEAVGEIMSLSVATPKHL  417

Query  419  KSIGLLSPDFQEDNETEINFLLKQALTIVGTLPFTYMLEKWRWMVFKGEIPKDQWMKKWW  478
            K +GLL PDF EDNET+INFLLKQAL IVGTLPFTYMLEKWRWMVFKGEIPK+QWMKKWW
Sbjct  418  KVLGLLPPDFHEDNETDINFLLKQALNIVGTLPFTYMLEKWRWMVFKGEIPKEQWMKKWW  477

Query  479  EMKREIVGVVEPVPHDETYCDPASLFHVSNDYSFIRYYTRTLYQFQFQEALCQAAKHEGP  538
            EMKREIVGVVEPVPHDETYCDPA+LFHV+NDYSFIRYYTRT++QFQFQEALCQ A+HEGP
Sbjct  478  EMKREIVGVVEPVPHDETYCDPATLFHVANDYSFIRYYTRTIFQFQFQEALCQTAQHEGP  537

Query  539  LHKCDISNSTEAGQKLFNMLRLGKSEPWTLALENVVGAKNMNVRPLLNYFEPLFTWLKDQ  598
            LHKCDISNST AG+KL  ML+LGKSEPWT ALEN+VG K M+VRPLLNYFEPLFTWLK+Q
Sbjct  538  LHKCDISNSTAAGEKLLQMLKLGKSEPWTRALENIVGKKQMDVRPLLNYFEPLFTWLKEQ  597

Query  599  NKNSFVGWSTDWSPYADQSIKVRISLKSALGDKAYEWNDNEMYLFRSSVAYAMRQYFLKV  658
            N+NSFVGW TDWSPYA +SIKVRISLKSALGDKAYEWNDNEMY FRSS+AYAMR+YF
Sbjct  598  NRNSFVGWLTDWSPYAAESIKVRISLKSALGDKAYEWNDNEMYFFRSSIAYAMREYFSNF  657

Query  659  KNQMILFGEEDVRVANLKPRISFNFFVTAPKNVSDIIPRTEVEKAIRMSRSRINDAFRLN  718
            KNQ I F  EDV V++LKPR+SFNFFVT+P +VSDIIPR+EVE+AIR SRSRINDAFRL+
Sbjct  658  KNQTIPFRAEDVWVSDLKPRVSFNFFVTSPNSVSDIIPRSEVEEAIRKSRSRINDAFRLD  717

Query  719  DNSLEFLGIQPTLGPPNQPPVSIWLIVFGVVMGVIVVGIVILIFTGIRDRKKKNKARSGE  778
            DNSLEFLGIQPTL PP QP V+IWLI FGVVMG++VVGI +LIFTGIR+RK+K++  S E
Sbjct  718  DNSLEFLGIQPTLEPPYQPAVTIWLIAFGVVMGLVVVGIGVLIFTGIRERKRKSQETSEE  777

Query  779  NPYASIDISKGENNPGFQNTDDVQTSF  805
            NPY+S+++SKGE+NPGFQN DDVQTSF
Sbjct  778  NPYSSMNLSKGESNPGFQNGDDVQTSF  804
```

**angiotensin-converting enzyme 2 isoform X2 [Desmodus rotundus]**

Sequence ID: XP_024425699.1 Length: 737 Number of Matches: 1

**Human vs Bat**

Range 1: 1 to 737 GenPept Graphics ▼ Next Match

| NW Score | Identities | Positives | Gaps |
|---|---|---|---|
| 3146 | 594/807(74%) | 657/807(81%) | 72/807(8%) |

```
Query  1    MSSSSWLLLSLVAVTAAQSTIEEQAKTFLDKFNHEAEDLFYQSSLASWNYNTNITEENVQ  60
            M
Sbjct       M-----------------------------------------------------------

Query  61   NMNNAGDKWSAFLKEQSTLAQMYPLQEIQNLTVKLQLQALQQNGSSVLSEDKSKRL--NT  118
                    WS F +  S +A+ YPL+ I+++ VK QLQALQQNG   L EDK K+L  N
Sbjct  2    --------WSTFYERNSNIAKTYPLETIKDVNVKRQLQALQQNG---LLEDKDKQLQLNA  50

Query  119  ILNTMSTIYSTGKVCNPDNPQECLLLEPGLNEIMANSLDYNERLWAWESWRSEVGKQLRP  178
            ILNTMSTIYSTGKVC P+NPQEC LL   GL +IM +S DYNERLWAWE WRS+VGKQLRP
Sbjct  51   ILNTMSTIYSTGKVCKPNNPQECYLLATGLEDIMQDSKDYNERLWAWEGWRSKVGKQLRP  110

Query  179  LYEEYVVLKNEMARANHYEDYGDYWRGDYEVNGVDGYDYSRGQLIEDVEHTFEEIKPLYE  238
            LYEEYVVLKNEMAR  +YEDYGDYWRGDYE  G  GY+YSR QLIEDVE+TF EIKPLYE
Sbjct  111  LYEEYVVLKNEMAREKNYEDYGDYWRGDYETEGSSGYEYSRNQLIEDVENTFAEIKPLYE  170

Query  239  HLHAYVRAKLMNAYPSYISPIGCLPAHLLGDMWGRFWTNLYSLTVPFGQKPNIDVTDAMV  298
            HLHAYVRAKLM+ YPS+ISP GCLPAHLLGDMWGRFWTNLY+LT PFG+KP IDVT AMV
Sbjct  171  HLHAYVRAKLMDTYPSHISPTGCLPAHLLGDMWGRFWTNLYNLTAPFGEKPTIDVTAAMV  230

Query  299  DQAWDAQRIFKEAEKFFVSVGLPNMTQGFWENSMLTDPGNVQKAVCHPTAWDLGKGDFRI  358
            DQ+WDAQRIFKEAEKFF SVGL +MTQGFW+NSMLT P + ++ VCHPTAWDLG  DFRI
Sbjct  231  DQSWDAQRIFKEAEKFFKSVGLFSMTQGFWDNSMLTKPDDGREVVCHPTAWDLGNKDFRI  290

Query  359  LMCTKVTMDDFLTAHHEMGHIQYDMAYAAQPFLLRNGANEGFHEAVGEIMSLSAATPKHL  418
             MCTKVTMDDFLTAHHEMGHIQYDMAYA Q FLLRNGANEGFHEAVGEIMSLS ATPKHL
Sbjct  291  KMCTKVTMDDFLTAHHEMGHIQYDMAYANQSFLLRNGANEGFHEAVGEIMSLSVATPKHL  350

Query  419  KSIGLLSPDFQEDNETEINFLLKQALTIVGTLPFTYMLEKWRWMVFKGEIPKDQWMKKWW  478
            K +GLL PDF EDNET+INFLLKQAL IVGTLPFTYMLEKWRWMVFKGEIPK+QWMKKWW
Sbjct  351  KVLGLLPPDFHEDNETDINFLLKQALNIVGTLPFTYMLEKWRWMVFKGEIPKEQWMKKWW  410

Query  479  EMKREIVGVVEPVPHDETYCDPASLFHVSNDYSFIRYYTRTLYQFQFQEALCQAAKHEGP  538
            EMKREIVGVVEPVPHDETYCDPA+LFHV+NDYSFIRYYTRT++QFQFQEALCQ A+HEGP
Sbjct  411  EMKREIVGVVEPVPHDETYCDPATLFHVANDYSFIRYYTRTIFQFQFQEALCQTAQHEGP  470

Query  539  LHKCDISNSTEAGQKLFNMLRLGKSEPWTLALENVVGAKNMNVRPLLNYFEPLFTWLKDQ  598
            LHKCDISNST AG+KL  ML+LGKSEPWT ALEN+VG K M+VRPLLNYFEPLFTWLK+Q
Sbjct  471  LHKCDISNSTAAGEKLLQMLKLGKSEPWTRALENIVGKKQMDVRPLLNYFEPLFTWLKEQ  530

Query  599  NKNSFVGWSTDWSPYADQSIKVRISLKSALGDKAYEWNDNEMYLFRSSVAYAMRQYFLKV  658
            N+NSFVGW TDWSPYA +SIKVRISLKSALGDKAYEWNDNEMY FRSS+AYAMR+YF
Sbjct  531  NRNSFVGWLTDWSPYAAESIKVRISLKSALGDKAYEWNDNEMYFFRSSIAYAMREYFSNF  590

Query  659  KNQMILFGEEDVRVANLKPRISFNFFVTAPKNVSDIIPRTEVEKAIRMSRSRINDAFRLN  718
            KNQ I F  EDV V++LKPR+SFNFFVT+P +VSDIIPR+EVE+AIR SRSRINDAFRL+
Sbjct  591  KNQTIPFRAEDVWVSDLKPRVSFNFFVTSPNSVSDIIPRSEVEEAIRKSRSRINDAFRLD  650

Query  719  DNSLEFLGIQPTLGPPNQPPVSIWLIVFGVVMGVIVVGIVILIFTGIRDRKKKNKARSGE  778
            DNSLEFLGIQPTL PP QP V+IWLI FGVVMG++VVGI +LIFTGIR+RK+K++  S E
Sbjct  651  DNSLEFLGIQPTLEPPYQPAVTIWLIAFGVVMGLVVVGIGVLIFTGIRERKRKSQETSEE  710

Query  779  NPYASIDISKGENNPGFQNTDDVQTSF  805
            NPY+S+++SKGE+NPGFQN DDVQTSF
Sbjct  711  NPYSSMNLSKGESNPGFQNGDDVQTSF  737
```

#### angiotensin-converting enzyme 2 isoform X1 [Eptesicus fuscus]

Sequence ID: XP_008153150.1 Length: 811 Number of Matches: 1

**Human vs Bat**

Range 1: 1 to 811 GenPept Graphics ▼ Next Match

| NW Score | Identities | Positives | Gaps |
|---|---|---|---|
| 3544 | 653/812(80%) | 722/812(88%) | 8/812(0%) |

```
Query  1    MSSSSWLLLSLVAVTAAQSTIEEQAKTFLDKFNHEAEDLFYQSSLASWNYNTNITEENVQ  60
            MS SSWL LSLVAVTAAQST E+ A  FL+ FN EAEDL ++S+LASWNYNTNIT+EN Q
Sbjct  1    MSGSSWLFLSLVAVTAAQSTTEKNATIFLENFNSEAEDLSHESALASWNYNTNITDENAQ  60

Query  61   NMNNAGDKWSAFLKEQSTLAQMYPLQEIQNLTVKLQLQALQQNGSSVLSEDKSKRLNTIL  120
             MN A  KWSAF ++QS LAQ YPLQEIQNLT+KLQLQ LQQNGSSVL+ DKSKRL+TIL
Sbjct  61   KMNEADSKWSAFYEKQSKLAQTYPLQEIQNLTIKLQLQVLQQNGSSVLTADKSKRLSTIL  120

Query  121  NTMSTIYSTGKVCNPDNPQECLLLEPGLNEIMANSLDYNERLWAWESWRSEVGKQLRPLY  180
             TMSTIYSTGKVCNP+NPQECL L  GL +IM  S DYN+RLW WE WRSEVGKQLRPLY
Sbjct  121  TTMSTIYSTGKVCNPNNPQECLTLS-GLEDIMEKSKDYNQRLWVWEGWRSEVGKQLRPLY  179

Query  181  EEYVVLKNEMARANHYEDYGDYWRGDYEVNGVDGYDYSRGQLIEDVEHTFEEIKPLYEHL  240
            EEYVVLKNEMAR N+YEDYGDYWRGDYE  G +GY+YSR QL EDV+  F EIKPLYEHL
Sbjct  180  EEYVVLKNEMARGNNYEDYGDYWRGDYETEGENGYNYSRSQLTEDVDRIFLEIKPLYEHL  239

Query  241  HAYVRAKLMNAYPSYISPIGCLPAHLLGDMWGRFWTNLYSLTVPFGQKPNIDVTDAMVDQ  300
            HAYVRAKLM+ YPS ISP GCLPAHLLGDMWGRFWTNLY+LTVPF QKPNIDVTDAM +Q
Sbjct  240  HAYVRAKLMDTYPSRISPTGCLPAHLLGDMWGRFWTNLYNLTVPFEQKPNIDVTDAMKEQ  299

Query  301  AWDAQRIFKEAEKFFVSVGLPNMTQGFWENSMLTDPGNVQKAVCHPTAWDLGKGDFRILM  360
            +WDA++IFKEAEKF++SVGLP+MT GFW NSMLT+PG+ +K VCHPTAWDLGK DFRI M
Sbjct  300  SWDAEKIFKEAEKFYMSVGLPSMTPGFWNNSMLTEPGDGRKVVCHPTAWDLGKNDFRIKM  359

Query  361  CTKVTMDDFLTAHHEMGHIQYDMAYAAQPFLLRNGANEGFHEAVGEIMSLSAATPKHLKS  420
            CTKVTMDDFLTAHHEMGHIQYDMAYA QP+LLRNGANEGFHEAVGE+MSLS ATPKHLK
Sbjct  360  CTKVTMDDFLTAHHEMGHIQYDMAYATQPYLLRNGANEGFHEAVGEVMSLSVATPKHLKG  419

Query  421  IGLLSPDFQEDNETEINFLLKQALTIVGTLPFTYMLEKWRWMVFKGEIPKDQWMKKWWEM  480
            +GLL  DF EDNETEINFLLKQAL IVGTLPFTYMLEKWRWMVFKGEIPK+QWMKKWWEM
Sbjct  420  MGLLPSDFSEDNETEINFLLKQALNIVGTLPFTYMLEKWRWMVFKGEIPKEQWMKKWWEM  479

Query  481  KREIVGVVEPVPHDETYCDPASLFHVSNDYSFIRYYTRTLYQFQFQEALCQAAKHEGPLH  540
            KREIVGVVEP+PHDETYCDPASLFHV+NDYSFIRY+TRT+++FQFQEALCQ AKH+GPLH
Sbjct  480  KREIVGVVEPLPHDETYCDPASLFHVANDYSFIRYFTRTIFEFQFQEALCQIAKHQGPLH  539

Query  541  KCDISNSTEAGQKLFNMLRLGKSEPWTLALENVVGAKNMNVRPLLNYFEPLFTWLKDQNK  600
            KCDISNSTEAG KL  ML+LGKS+PWT ALE + G K M+ +PLLNYFEPLFTWLK+QN
Sbjct  540  KCDISNSTEAGNKLLEMLKLGKSKPWTFALEKITGTKKMDAKPLLNYFEPLFTWLKEQNG  599

Query  601  NS-------FVGWSTDWSPYADQSIKVRISLKSALGDKAYEWNDNEMYLFRSSVAYAMRQ  653
            NS       +VGWS+DWSPYADQSIKVRISLKSALG+KAYEWNDNEMYLFRSSVAYAMR+
Sbjct  600  NSVGWHSGNYVGWSSDWSPYADQSIKVRISLKSALGEKAYEWNDNEMYLFRSSVAYAMRE  659

Query  654  YFLKVKNQMILFGEEDVRVANLKPRISFNFFVTAPKNVSDIIPRTEVEKAIRMSRSRIND  713
            YFLKVKNQ I F  EDV V  ++KPR+SF FFVT+P N+SDIIPR+EVE AIRMSRSRIN
Sbjct  660  YFLKVKNQTIPFRAEDVWVNDVKPRVSFKFFVTSPTNMSDIIPRSEVEDAIRMSRSRINA  719

Query  714  AFRLNDNSLEFLGIQPTLGPPNQPPVSIWLIVFGVVMGVIVVGIVILIFTGIRDRKKKNK  773
            AFRL+DNSLEFLGIQPTLGPP QPPV+IWLIVFGVVMGV+V+GI +LIFTGIRDRKKKN+
Sbjct  720  AFRLDDNSLEFLGIQPTLGPPYQPPVTIWLIVFGVVMGVVVIGIGVLIFTGIRDRKKKNQ  779

Query  774  ARSGENPYASIDISKGENNPGFQNTDDVQTSF  805
              + ENPY+S+++SKGENNPGFQ+ DDVQTSF
Sbjct  780  PGNEENPYSSVNLSKGENNPGFQSGDDVQTSF  811
```

#### angiotensin-converting enzyme 2 isoform X2 [Eptesicus fuscus]

Sequence ID: XP_027986092.1 Length: 799 Number of Matches: 1

**Human vs Bat**

Range 1: 1 to 799 GenPept Graphics ▼ Next Match

| NW Score | Identities | Positives | Gaps |
|---|---|---|---|
| 3498 | 648/805(80%) | 716/805(88%) | 6/805(0%) |

```
Query  1    MSSSSWLLLSLVAVTAAQSTIEEQAKTFLDKFNHEAEDLFYQSSLASWNYNTNITEENVQ  60
            MS SSWL LSLVAVTAAQST E+ A  FL+ FN EAEDL ++S+LASWNYNTNIT+EN Q
Sbjct  1    MSGSSWLFLSLVAVTAAQSTTEKNATIFLENFNSEAEDLSHESALASWNYNTNITDENAQ  60

Query  61   NMNNAGDKWSAFLKEQSTLAQMYPLQEIQNLTVKLQLQALQQNGSSVLSEDKSKRLNTIL  120
             MN A  KWSAF ++QS LAQ YPLQEIQNLT+KLQLQ LQQNGSSVL+ DKSKRL+TIL
Sbjct  61   KMNEADSKWSAFYEKQSKLAQTYPLQEIQNLTIKLQLQVLQQNGSSVLTADKSKRLSTIL  120

Query  121  NTMSTIYSTGKVCNPDNPQECLLLEPGLNEIMANSLDYNERLWAWESWRSEVGKQLRPLY  180
             TMSTIYSTGKVCNP+NPQECL L  GL +IM  S DYN+RLW WE WRSEVGKQLRPLY
Sbjct  121  TTMSTIYSTGKVCNPNNPQECLTLS-GLEDIMEKSKDYNQRLWVWEGWRSEVGKQLRPLY  179

Query  181  EEYVVLKNEMARANHYEDYGDYWRGDYEVNGVDGYDYSRGQLIEDVEHTFEEIKPLYEHL  240
            EEYVVLKNEMAR N+YEDYGDYWRGDYE  G +GY+YSR QL EDV+  F EIKPLYEHL
Sbjct  180  EEYVVLKNEMARGNNYEDYGDYWRGDYETEGENGYNYSRSQLTEDVDRIFLEIKPLYEHL  239

Query  241  HAYVRAKLMNAYPSYISPIGCLPAHLLGDMWGRFWTNLYSLTVPFGQKPNIDVTDAMVDQ  300
            HAYVRAKLM+ YPS ISP GCLPAHLLGDMWGRFWTNLY+LTVPF QKPNIDVTDAM +Q
Sbjct  240  HAYVRAKLMDTYPSRISPTGCLPAHLLGDMWGRFWTNLYNLTVPFEQKPNIDVTDAMKEQ  299

Query  301  AWDAQRIFKEAEKFFVSVGLPNMTQGFWENSMLTDPGNVQKAVCHPTAWDLGKGDFRILM  360
            +WDA++IFKEAEKF++SVGLP+MT GFW NSMLT+PG+ +K VCHPTAWDLGK DFRI M
Sbjct  300  SWDAEKIFKEAEKFYMSVGLPSMTPGFWNNSMLTEPGDGRKVVCHPTAWDLGKNDFRIKM  359

Query  361  CTKVTMDDFLTAHHEMGHIQYDMAYAAQPFLLRNGANEGFHEAVGEIMSLSAATPKHLKS  420
            CTKVTMDDFLTAHHEMGHIQYDMAYA QP+LLRNGANEGFHEAVGE+MSLS ATPKHLK
Sbjct  360  CTKVTMDDFLTAHHEMGHIQYDMAYATQPYLLRNGANEGFHEAVGEVMSLSVATPKHLKG  419

Query  421  IGLLSPDFQEDNETEINFLLKQALTIVGTLPFTYMLEKWRWMVFKGEIPKDQWMKKWWEM  480
            +GLL  DF EDNETEINFLLKQAL IVGTLPFTYMLEKWRWMVFKGEIPK+QWMKKWWEM
Sbjct  420  MGLLPSDFSEDNETEINFLLKQALNIVGTLPFTYMLEKWRWMVFKGEIPKEQWMKKWWEM  479

Query  481  KREIVGVVEPVPHDETYCDPASLFHVSNDYSFIRYYTRTLYQFQFQEALCQAAKHEGPLH  540
            KREIVGVVEP+PHDETYCDPASLFHV+NDYSFIRY+TRT+++FQFQEALCQ AKH+GPLH
Sbjct  480  KREIVGVVEPLPHDETYCDPASLFHVANDYSFIRYFTRTIFEFQFQEALCQIAKHQGPLH  539

Query  541  KCDISNSTEAGQKLFNMLRLGKSEPWTLALENVVGAKNMNVRPLLNYFEPLFTWLKDQNK  600
            KCDISNSTEAG KL  ML+LGKS+PWT ALE + G K M+ +PLLNYFEPLFTWLK+QN
Sbjct  540  KCDISNSTEAGNKLLEMLKLGKSKPWTFALEKITGTKKMDAKPLLNYFEPLFTWLKEQNG  599

Query  601  NSFVGWSTDWSPYADQSIKVRISLKSALGDKAYEWNDNEMYLFRSSVAYAMRQYFLKVKN  660
            NS VGW +D    ADQSIKVRISLKSALG+KAYEWNDNEMYLFRSSVAYAMR+YFLKVKN
Sbjct  600  NS-VGWHSD----ADQSIKVRISLKSALGEKAYEWNDNEMYLFRSSVAYAMREYFLKVKN  654

Query  661  QMILFGEEDVRVANLKPRISFNFFVTAPKNVSDIIPRTEVEKAIRMSRSRINDAFRLNDN  720
            Q I F  EDV V  ++KPR+SF FFVT+P N+SDIIPR+EVE AIRMSRSRIN AFRL+DN
Sbjct  655  QTIPFRAEDVWVNDVKPRVSFKFFVTSPTNMSDIIPRSEVEDAIRMSRSRINAAFRLDDN  714

Query  721  SLEFLGIQPTLGPPNQPPVSIWLIVFGVVMGVIVVGIVILIFTGIRDRKKKNKARSGENP  780
            SLEFLGIQPTLGPP QPPV+IWLIVFGVVMGV+V+GI +LIFTGIRDRKKKN+  + ENP
Sbjct  715  SLEFLGIQPTLGPPYQPPVTIWLIVFGVVMGVVVIGIGVLIFTGIRDRKKKNQPGNEENP  774

Query  781  YASIDISKGENNPGFQNTDDVQTSF  805
            Y+S+++SKGENNPGFQ+ DDVQTSF
Sbjct  775  YSSVNLSKGENNPGFQSGDDVQTSF  799
```

**angiotensin-converting enzyme 2 isoform X1 [Myotis lucifugus]**

Sequence ID: XP_023609437.1 Length: 819 Number of Matches: 1

See 1 more title(s)

#### Human vs Bat

Range 1: 1 to 819 GenPept Graphics ▼ Next Match

| NW Score | Identities | Positives | Gaps |
| --- | --- | --- | --- |
| 3528 | 650/820(79%) | 723/820(88%) | 16/820(1%) |

```
Query  1    MSSSSWLLLSLVAVTAAQSTIEEQAKTFLDKFNHEAEDLFYQSSLASWNYNTNITEENVQ  60
            MS SSWL LSLVAV AAQS+ EE+AK FL+ FN +AEDL ++S+LASWNYNTNIT+ENVQ
Sbjct  1    MSGSSWLFLSLVAVAAAQSSTEEKAKIFLENFNSKAEDLSHESALASWNYNTNITDENVQ  60

Query  61   NMNNAGDKWSAFLKEQSTLAQMYPLQEIQNLTVKLQLQALQQNGSSVLSEDKSKRLNTIL  120
             MN A  KWSAF ++QS LAQ YPLQEIQN T+K QLQ LQQNGSSVLS DKSKRLNTIL
Sbjct  61   KMNEADSKWSAFYEQQSKLAQTYPLQEIQNSTIKRQLQVLQQNGSSVLSADKSKRLNTIL  120

Query  121  NTMSTIYSTGKVCNPDNPQECLLLEPGLNEIMANSLDYNERLWAWESWRSEVGKQLRPLY  180
             TMSTIYSTGKVCNP+NPQEC   L   GL EIM   S DYN+RLW WE WRSEVGKQLRPLY
Sbjct  121  TTMSTIYSTGKVCNPNNPQECFTLA-GLEEIMEKSKDYNQRLWVWEGWRSEVGKQLRPLY  179

Query  181  EEYVVLKNEMARANHYEDYGDYWRGDYEVNGVDGYDYSRGQLIEDVEHTFEEIKPLYEHL  240
            EEYV LKNEMAR N+YEDYGDYWRGDYE   G DGY+YSR QL EDVE   F EIKPLYEHL
Sbjct  180  EEYVDLKNEMARGNNYEDYGDYWRGDYETEGEDGYNYSRNQLTEDVERIFLEIKPLYEHL  239

Query  241  HAYVRAKLMNAYPSYISPIGCLPAHLLGDMWGRFWTNLYSLTVPFGQKPNIDVTDAMVDQ  300
            HAYVRAKL+NAYPS ISP G LPAHLLGDMWGRFWTNLY+LTVPF QKPNIDVT AMV+Q
Sbjct  240  HAYVRAKLVNAYPSRISPTGYLPAHLLGDMWGRFWTNLYNLTVPFEQKPNIDVTGAMVEQ  299

Query  301  AWDAQRIFKEAEKFFVSVGLPNMTQGFWENSMLTDPGNVQKAVCHPTAWDLGKGDFRILM  360
            +WDA++IFKEAEKF++SVGLP+MT GFW NSMLT+PG+ +K VCHPTAWDLGKGDFRI M
Sbjct  300  SWDAEKIFKEAEKFYISVGLPSMTPGFWNNSMLTEPGDGRKVVCHPTAWDLGKGDFRIKM  359

Query  361  CTKVTMDDFLTAHHEMGHIQYDMAYAAQPFLLRNGANEGFHEAVGEIMSLSAATPKHLKS  420
            CTKVTMDDFLTAHHEMGHIQYDMAYA QP+LLRNGANEGFHEAVGE+MSLS ATPKHLK
Sbjct  360  CTKVTMDDFLTAHHEMGHIQYDMAYATQPYLLRNGANEGFHEAVGEVMSLSVATPKHLKG  419

Query  421  IGLLSPDFQEDNETEINFLLKQALTIVGTLPFTYMLEKWRWMVFKGEIPKDQWMKKWWEM  480
            +GLL PDF EDNETEINFLLKQAL IVGTLPFTYMLEKWRWMVFKGEIPK+QWMKKWWEM
Sbjct  420  MGLLPPDFSEDNETEINFLLKQALNIVGTLPFTYMLEKWRWMVFKGEIPKEQWMKKWWEM  479

Query  481  KREIVGVVEPVPHDETYCDPASLFHVSNDYSFIRYYTRTLYQFQFQEALCQAAKHEGPLH  540
            KR+IVGV+EP+PHDETYCDPASLFHV+NDYSFIRY+TRT+++FQFQEALCQ AKH+GPLH
Sbjct  480  KRDIVGVMEPLPHDETYCDPASLFHVANDYSFIRYFTRTIFEFQFQEALCQIAKHQGPLH  539

Query  541  KCDISNSTEAGQKLFNMLRLGKSEPWTLALENVVGAKNMNVRPLLNYFEPLFTWLKDQNK  600
            KCDISNS EAG KL  ML+LGKSEPWTLAL+ +VG K M+ +PLLNYFEPLFTWLK+QN
Sbjct  540  KCDISNSKEAGNKLLEMLKLGKSEPWTLALDKIVGTKKMDAKPLLNYFEPLFTWLKEQNG  599

Query  601  NSF---------------VGWSTDWSPYADQSIKVRISLKSALGDKAYEWNDNEMYLFRS  645
            NS                VGW +DWSPYA+QSIKVRISL+SALG+KAY+WNDNEMYLF+S
Sbjct  600  NSVGWNSGNSVESRSGNSVGWFSDWSPYAEQSIKVRISLQSALGEKAYKWNDNEMYLFQS  659

Query  646  SVAYAMRQYFLKVKNQMILFGEEDVRVANLKPRISFNFFVTAPKNVSDIIPRTEVEKAIR  705
            SVAYAMR+YFLK KNQ I FG E+VRV ++KPR+SF F+VT+PKN+S +IPR+EVE AIR
Sbjct  660  SVAYAMREYFLKEKNQTIPFGVENVRVNDIKPRVSFKFYVTSPKNMSVVIPRSEVEDAIR  719

Query  706  MSRSRINDAFRLNDNSLEFLGIQPTLGPPNQPPVSIWLIVFGVVMGVIVVGIVILIFTGI  765
            MSRSRINDAFRL+DN+LEFLGIQPTLGPPNQPPV+IWLIVFGVVMGV V+GI +LIFTGI
Sbjct  720  MSRSRINDAFRLDDNTLEFLGIQPTLGPPNQPPVTIWLIVFGVVMGVAVIGIAVLIFTGI  779

Query  766  RDRKKKNKARSGENPYASIDISKGENNPGFQNTDDVQTSF  805
            RDRKKK +A + ENPY+S+++SKGENNPGFQN DDVQTSF
Sbjct  780  RDRKKKQAGNEENPYSSVNLSKGENNPGFQNGDDVQTSF  819
```


**angiotensin-converting enzyme 2 isoform X1 [Myotis lucifugus]**

Sequence ID: XP_023609437.1 Length: 819 Number of Matches: 1

See 1 more title(s)

#### Human vs Bat

Range 1: 1 to 819 GenPept Graphics ▼ Next Matc

| NW Score | Identities | Positives | Gaps |
| --- | --- | --- | --- |
| 3528 | 650/820(79%) | 723/820(88%) | 16/820(1%) |

```
Query  1    MSSSSWLLLSLVAVTAAQSTIEEQAKTFLDKFNHEAEDLFYQSSLASWNYNTNITEENVQ  60
            MS SSWL LSLVAV AAQS+ EE+AK FL+ FN +AEDL ++S+LASWNYNTNIT+ENVQ
Sbjct  1    MSGSSWLFLSLVAVAAAQSSTEEKAKIFLENFNSKAEDLSHESALASWNYNTNITDENVQ  60

Query  61   NMNNAGDKWSAFLKEQSTLAQMYPLQEIQNLTVKLQLQALQQNGSSVLSEDKSKRLNTIL  120
             MN A  KWSAF ++QS LAQ YPLQEIQN T+K QLQ LQQNGSSVLS DKSKRLNTIL
Sbjct  61   KMNEADSKWSAFYEQQSKLAQTYPLQEIQNSTIKRQLQVLQQNGSSVLSADKSKRLNTIL  120

Query  121  NTMSTIYSTGKVCNPDNPQECLLLEPGLNEIMANSLDYNERLWAWESWRSEVGKQLRPLY  180
             TMSTIYSTGKVCNP+NPQEC   L   GL EIM   S DYN+RLW WE WRSEVGKQLRPLY
Sbjct  121  TTMSTIYSTGKVCNPNNPQECFTLA-GLEEIMEKSKDYNQRLWVWEGWRSEVGKQLRPLY  179

Query  181  EEYVVLKNEMARANHYEDYGDYWRGDYEVNGVDGYDYSRGQLIEDVEHTFEEIKPLYEHL  240
            EEYV LKNEMAR N+YEDYGDYWRGDYE   G DGY+YSR QL EDVE   F EIKPLYEHL
Sbjct  180  EEYVDLKNEMARGNNYEDYGDYWRGDYETEGEDGYNYSRNQLTEDVERIFLEIKPLYEHL  239

Query  241  HAYVRAKLMNAYPSYISPIGCLPAHLLGDMWGRFWTNLYSLTVPFGQKPNIDVTDAMVDQ  300
            HAYVRAKL+NAYPS ISP G LPAHLLGDMWGRFWTNLY+LTVPF QKPNIDVT AMV+Q
Sbjct  240  HAYVRAKLVNAYPSRISPTGYLPAHLLGDMWGRFWTNLYNLTVPFEQKPNIDVTGAMVEQ  299

Query  301  AWDAQRIFKEAEKFFVSVGLPNMTQGFWENSMLTDPGNVQKAVCHPTAWDLGKGDFRILM  360
            +WDA++IFKEAEKF++SVGLP+MT GFW NSMLT+PG+ +K VCHPTAWDLGKGDFRI M
Sbjct  300  SWDAEKIFKEAEKFYISVGLPSMTPGFWNNSMLTEPGDGRKVVCHPTAWDLGKGDFRIKM  359

Query  361  CTKVTMDDFLTAHHEMGHIQYDMAYAAQPFLLRNGANEGFHEAVGEIMSLSAATPKHLKS  420
            CTKVTMDDFLTAHHEMGHIQYDMAYA QP+LLRNGANEGFHEAVGE+MSLS ATPKHLK
Sbjct  360  CTKVTMDDFLTAHHEMGHIQYDMAYATQPYLLRNGANEGFHEAVGEVMSLSVATPKHLKG  419

Query  421  IGLLSPDFQEDNETEINFLLKQALTIVGTLPFTYMLEKWRWMVFKGEIPKDQWMKKWWEM  480
            +GLL PDF EDNETEINFLLKQAL IVGTLPFTYMLEKWRWMVFKGEIPK+QWMKKWWEM
Sbjct  420  MGLLPPDFSEDNETEINFLLKQALNIVGTLPFTYMLEKWRWMVFKGEIPKEQWMKKWWEM  479

Query  481  KREIVGVVEPVPHDETYCDPASLFHVSNDYSFIRYYTRTLYQFQFQEALCQAAKHEGPLH  540
            KR+IVGV+EP+PHDETYCDPASLFHV+NDYSFIRY+TRT+++FQFQEALCQ AKH+GPLH
Sbjct  480  KRDIVGVMEPLPHDETYCDPASLFHVANDYSFIRYFTRTIFEFQFQEALCQIAKHQGPLH  539

Query  541  KCDISNSTEAGQKLFNMLRLGKSEPWTLALENVVGAKNMNVRPLLNYFEPLFTWLKDQNK  600
            KCDISNS EAG KL  ML+LGKSEPWTLAL+ +VG K M+ +PLLNYFEPLFTWLK+QN
Sbjct  540  KCDISNSKEAGNKLLEMLKLGKSEPWTLALDKIVGTKKMDAKPLLNYFEPLFTWLKEQNG  599

Query  601  NSF---------------VGWSTDWSPYADQSIKVRISLKSALGDKAYEWNDNEMYLFRS  645
            NS                VGW +DWSPYA+QSIKVRISL+SALG+KAY+WNDNEMYLF+S
Sbjct  600  NSVGWNSGNSVESRSGNSVGWFSDWSPYAEQSIKVRISLQSALGEKAYKWNDNEMYLFQS  659

Query  646  SVAYAMRQYFLKVKNQMILFGEEDVRVANLKPRISFNFFVTAPKNVSDIIPRTEVEKAIR  705
            SVAYAMR+YFLK KNQ I FG E+VRV ++KPR+SF F+VT+PKN+S +IPR+EVE AIR
Sbjct  660  SVAYAMREYFLKEKNQTIPFGVENVRVNDIKPRVSFKFYVTSPKNMSVVIPRSEVEDAIR  719

Query  706  MSRSRINDAFRLNDNSLEFLGIQPTLGPPNQPPVSIWLIVFGVVMGVIVVGIVILIFTGI  765
            MSRSRINDAFRL+DN+LEFLGIQPTLGPPNQPPV+IWLIVFGVVMGV V+GI +LIFTGI
Sbjct  720  MSRSRINDAFRLDDNTLEFLGIQPTLGPPNQPPVTIWLIVFGVVMGVAVIGIAVLIFTGI  779

Query  766  RDRKKKNKARSGENPYASIDISKGENNPGFQNTDDVQTSF  805
            RDRKKK +A + ENPY+S+++SKGENNPGFQN DDVQTSF
Sbjct  780  RDRKKKQAGNEENPYSSVNLSKGENNPGFQNGDDVQTSF  819
```

#### angiotensin-converting enzyme 2 isoform X2 [Myotis lucifugus]

Sequence ID: XP_023609439.1 Length: 799 Number of Matches: 1

**Human vs Bat**

Range 1: 1 to 799 GenPept Graphics ▼ Next Matc

| NW Score | Identities | Positives | Gaps |
|---|---|---|---|
| 3502 | 646/805(80%) | 720/805(89%) | 6/805(0%) |

```
Query  1    MSSSSWLLLSLVAVTAAQSTIEEQAKTFLDKFNHEAEDLFYQSSLASWNYNTNITEENVQ  60
            MS SSWL LSLVAV AAQS+ EE+AK FL+ FN +AEDL ++S+LASWNYNTNIT+ENVQ
Sbjct  1    MSGSSWLFLSLVAVAAAQSSTEEKAKIFLENFNSKAEDLSHESALASWNYNTNITDENVQ  60

Query  61   NMNNAGDKWSAFLKEQSTLAQMYPLQEIQNLTVKLQLQALQQNGSSVLSEDKSKRLNTIL  120
             MN A  KWSAF ++QS LAQ YPLQEIQN T+K QLQ LQQNGSSVLS DKSKRLNTIL
Sbjct  61   KMNEADSKWSAFYEQQSKLAQTYPLQEIQNSTIKRQLQVLQQNGSSVLSADKSKRLNTIL  120

Query  121  NTMSTIYSTGKVCNPDNPQECLLLEPGLNEIMANSLDYNERLWAWESWRSEVGKQLRPLY  180
             TMSTIYSTGKVCNP+NPQEC  L  GL EIM  S DYN+RLW WE WRSEVGKQLRPLY
Sbjct  121  TTMSTIYSTGKVCNPNNPQECFTLA-GLEEIMEKSKDYNQRLWVWEGWRSEVGKQLRPLY  179

Query  181  EEYVVLKNEMARANHYEDYGDYWRGDYEVNGVDGYDYSRGQLIEDVEHTFEEIKPLYEHL  240
            EEYV LKNEMAR N+YEDYGDYWRGDYE  G DGY+YSR QL EDVE  F EIKPLYEHL
Sbjct  180  EEYVDLKNEMARGNNYEDYGDYWRGDYETEGEDGYNYSRNQLTEDVERIFLEIKPLYEHL  239

Query  241  HAYVRAKLMNAYPSYISPIGCLPAHLLGDMWGRFWTNLYSLTVPFGQKPNIDVTDAMVDQ  300
            HAYVRAKL+NAYPS ISP G LPAHLLGDMWGRFWTNLY+LTVPF QKPNIDVT AMV+Q
Sbjct  240  HAYVRAKLVNAYPSRISPTGYLPAHLLGDMWGRFWTNLYNLTVPFEQKPNIDVTGAMVEQ  299

Query  301  AWDAQRIFKEAEKFFVSVGLPNMTQGFWENSMLTDPGNVQKAVCHPTAWDLGKGDFRILM  360
            +WDA++IFKEAEKF++SVGLP+MT GFW NSMLT+PG+ +K VCHPTAWDLGKGDFRI M
Sbjct  300  SWDAEKIFKEAEKFYISVGLPSMTPGFWNNSMLTEPGDGRKVVCHPTAWDLGKGDFRIKM  359

Query  361  CTKVTMDDFLTAHHEMGHIQYDMAYAAQPFLLRNGANEGFHEAVGEIMSLSAATPKHLKS  420
            CTKVTMDDFLTAHHEMGHIQYDMAYA QP+LLRNGANEGFHEAVGE+MSLS ATPKHLK
Sbjct  360  CTKVTMDDFLTAHHEMGHIQYDMAYATQPYLLRNGANEGFHEAVGEVMSLSVATPKHLKG  419

Query  421  IGLLSPDFQEDNETEINFLLKQALTIVGTLPFTYMLEKWRWMVFKGEIPKDQWMKKWWEM  480
            +GLL PDF EDNETEINFLLKQAL IVGTLPFTYMLEKWRWMVFKGEIPK+QWMKKWWEM
Sbjct  420  MGLLPPDFSEDNETEINFLLKQALNIVGTLPFTYMLEKWRWMVFKGEIPKEQWMKKWWEM  479

Query  481  KREIVGVVEPVPHDETYCDPASLFHVSNDYSFIRYYTRTLYQFQFQEALCQAAKHEGPLH  540
            KR+IVGV+EP+PHDETYCDPASLFHV+NDYSFIRY+TRT+++FQFQEALCQ AKH+GPLH
Sbjct  480  KRDIVGVMEPLPHDETYCDPASLFHVANDYSFIRYFTRTIFEFQFQEALCQIAKHQGPLH  539

Query  541  KCDISNSTEAGQKLFNMLRLGKSEPWTLALENVVGAKNMNVRPLLNYFEPLFTWLKDQNK  600
            KCDISNS EAG KL  ML+LGKSEPWTLAL+ +VG K M+ +PLLNYFEPLFTWLK+QN
Sbjct  540  KCDISNSKEAGNKLLEMLKLGKSEPWTLALDKIVGTKKMDAKPLLNYFEPLFTWLKEQNG  599

Query  601  NSFVGWSTDWSPYADQSIKVRISLKSALGDKAYEWNDNEMYLFRSSVAYAMRQYFLKVKN  660
            NS VGW++D    A+QSIKVRISL+SALG+KAY+WNDNEMYLF+SSVAYAMR+YFLK KN
Sbjct  600  NS-VGWNSD----AEQSIKVRISLQSALGEKAYKWNDNEMYLFQSSVAYAMREYFLKEKN  654

Query  661  QMILFGEEDVRVANLKPRISFNFFVTAPKNVSDIIPRTEVEKAIRMSRSRINDAFRLNDN  720
            Q I FG E+VRV ++KPR+SF F+VT+PKN+S +IPR+EVE AIRMSRSRINDAFRL+DN
Sbjct  655  QTIPFGVENVRVNDIKPRVSFKFYVTSPKNMSVVIPRSEVEDAIRMSRSRINDAFRLDDN  714

Query  721  SLEFLGIQPTLGPPNQPPVSIWLIVFGVVMGVIVVGIVILIFTGIRDRKKKNKARSGENP  780
            +LEFLGIQPTLGPPNQPPV+IWLIVFGVVMGV V+GI +LIFTGIRDRKKK +A + ENP
Sbjct  715  TLEFLGIQPTLGPPNQPPVTIWLIVFGVVMGVAVIGIAVLIFTGIRDRKKKKQAGNEENP  774

Query  781  YASIDISKGENNPGFQNTDDVQTSF  805
            Y+S+++SKGENNPGFQN DDVQTSF
Sbjct  775  YSSVNLSKGENNPGFQNGDDVQTSF  799
```

#### angiotensin-converting enzyme 2 [Phyllostomus discolor]

Sequence ID: XP_028378317.1 Length: 804 Number of Matches: 1

**Human vs Bat**

Range 1: 1 to 804 GenPept Graphics ▼ Next Match

| NW Score | Identities | Positives | Gaps |
|---|---|---|---|
| 3447 | 642/807(80%) | 710/807(87%) | 5/807(0%) |

```
Query  1    MSSSSWLLLSLVAVTAAQSTIEEQAKTFLDKFNHEAEDLFYQSSLASWNYNTNITEENVQ  60
            MS SSWL LSLVAV AAQ+  EE A+ FL+ FN+EAE+L+YQSSLA+WNYNTNIT+ENVQ
Sbjct  1    MSGSSWLFLSLVAVAAAQTPTEEDARKFLENFNNEAEELYYQSSLAAWNYNTNITDENVQ  60

Query  61   NMNNAGDKWSAFLKEQSTLAQMYPLQEIQNLTVKLQLQALQQNGSSVLSEDKSKRLNTIL  120
             MN A  +WS F +EQS LA+ YPL  I ++TVK QLQALQQNG   L +DK KRLNTIL
Sbjct  61   KMNEADKRWSTFYEEQSRLAKNYPLATITDVTVKHQLQALQQNG---LLDDKDKRLNTIL  117

Query  121  NTMSTIYSTGKVCNPDNPQECLLLEPGLNEIMANSLDYNERLWAWESWRSEVGKQLRPLY  180
            NTMSTIYSTGKVC P+NPQECLLL  GL +IM NS DYNERLWAWE WRSEVGK+LRPLY
Sbjct  118  NTMSTIYSTGKVCKPNNPQECLLLATGLEDIMHNSKDYNERLWAWEGWRSEVGKRLRPLY  177

Query  181  EEYVVLKNEMARANHYEDYGDYWRGDYEVNGVDGYDYSRGQLIEDVEHTFEEIKPLYEHL  240
            EEYVVLKNEMAR  +YEDYGDYWRGDYE  G   Y YSR QLI+DVE TFEEIKPLYE+L
Sbjct  178  EEYVVLKNEMAREKNYEDYGDYWRGDYETEGTSDYGYSRNQLIKDVESTFEEIKPLYENL  237

Query  241  HAYVRAKLMNAYPSYISPIGCLPAHLLGDMWGRFWTNLYSLTVPFGQKPNIDVTDAMVDQ  300
            HAYVRAKLM+AYPS ISP GCLPAHLLGDMWGRFWTNLY LT PF +KP IDVT AMV Q
Sbjct  238  HAYVRAKLMDAYPSRISPTGCLPAHLLGDMWGRFWTNLYDLTAPFPEKPTIDVTSAMVAQ  297

Query  301  AWDAQRIFKEAEKFFVSVGLPNMTQGFWENSMLTDPGNVQKAVCHPTAWDLGKGDFRILM  360
            +WDAQRIFKEAEKFFVSVGL NMTQGFW+NSMLT P + ++ VCHPTAWDLGK DFRI M
Sbjct  298  SWDAQRIFKEAEKFFVSVGLFNMTQGFWDNSMLTKPDDGREVVCHPTAWDLGKKDFRIKM  357

Query  361  CTKVTMDDFLTAHHEMGHIQYDMAYAAQPFLLRNGANEGFHEAVGEIMSLSAATPKHLKS  420
            CTKVTMDDFLTAHHEMGHIQYDMAYA QPFLLRNGANEGFHEAVGEIMSLSAATPKHLK
Sbjct  358  CTKVTMDDFLTAHHEMGHIQYDMAYADQPFLLRNGANEGFHEAVGEIMSLSAATPKHLKV  417

Query  421  IGLLSPDFQEDNETEINFLLKQALTIVGTLPFTYMLEKWRWMVFKGEIPKDQWMKKWWEM  480
            +GLL  DF+EDNET+INFLLKQAL IVGTLPFTYMLEKWRWMVFKGEIPK+QWMKKWWEM
Sbjct  418  LGLLPSDFREDNETDINFLLKQALNIVGTLPFTYMLEKWRWMVFKGEIPKEQWMKKWWEM  477

Query  481  KREIVGVVEPVPHDETYCDPASLFHVSNDYSFIRYYTRTLYQFQFQEALCQAAKHEGPLH  540
            KREIVGVVEPVPH+ETYCDPA+LFHV+NDYSFIRYYTRT++QFQFQEALC+ A+HEGPLH
Sbjct  478  KREIVGVVEPVPHNETYCDPAALFHVANDYSFIRYYTRTIFQFQFQEALCRIAQHEGPLH  537

Query  541  KCDISNSTEAGQKLFNMLRLGKSEPWTLALENVVGAKNMNVRPLLNYFEPLFTWLKDQNK  600
            KCDISNST AGQKL  ML+LGKSEPWT ALE  VG K M+V+PLLNYFEPLFTWLKDQN+
Sbjct  538  KCDISNSTAAGQKLLEMLKLGKSEPWTRALETFVGKKQMDVKPLLNYFEPLFTWLKDQNR  597

Query  601  NSFVGWSTDWSPYAD--QSIKVRISLKSALGDKAYEWNDNEMYLFRSSVAYAMRQYFLKV  658
            NSFVGW T WSPYA   QSIKVRISLKSALGDKAYEWNDNEMY F+SS+AYAMR++F  +
Sbjct  598  NSFVGWRTTWSPYAAAAQSIKVRISLKSALGDKAYEWNDNEMYFFQSSIAYAMREHFSDL  657

Query  659  KNQMILFGEEDVRVANLKPRISFNFFVTAPKNVSDIIPRTEVEKAIRMSRSRINDAFRLN  718
            K Q+I F  EDV+V +LKPR+SFNFFVT+P + SDI+PR+EVE+AIR SRSRINDAFRL+
Sbjct  658  KKQVIPFRAEDVKVYDLKPRVSFNFFVTSPNDTSDIVPRSEVEEAIRKSRSRINDAFRLD  717

Query  719  DNSLEFLGIQPTLGPPNQPPVSIWLIVFGVVMGVIVVGIVILIFTGIRDRKKKNKARSGE  778
            DNSLEFLGIQPTL PP QP V+IWLIVFGVVMGV+VVGI +LIFTGIRDR+KK++    E
Sbjct  718  DNSLEFLGIQPTLEPPYQPAVTIWLIVFGVVMGVVVVGIAVLIFTGIRDRRKKDEPSIEE  777

Query  779  NPYASIDISKGENNPGFQNTDDVQTSF  805
            NPY+S+++SKGE+N GFQN DDVQTSF
Sbjct  778  NPYSSVNLSKGESNLGFQNGDDVQTSF  804
```

**PREDICTED: angiotensin-converting enzyme 2 [Hipposideros armiger]**

Sequence ID: XP_019522936.1 Length: 806 Number of Matches: 1

See 2 more title(s)

#### Human vs Bat

Range 1: 1 to 806 GenPept Graphics  Next Match

| NW Score | Identities | Positives | Gaps |
|---|---|---|---|
| 3519 | 649/806(81%) | 721/806(89%) | 1/806(0%) |

```
Query  1    MSSSSWLLLSLVAVTAAQSTIEEQAKTFLDKFNHEAEDLFYQSSLASWNYNTNITEENVQ  60
            MS SSWLLLSLVAV AAQS  E+ AK FLDKFN EAEDL + SSLASW+YNTNIT+ENVQ
Sbjct  1    MSGSSWLLLSLVAVAAAQSNSEDLAKEFLDKFNTEAEDLSHLSSLASWDYNTNITDENVQ  60

Query  61   NMNNAGDKWSAFLKEQSTLAQMYPLQEIQNLTVKLQLQALQQNGSSVLSEDKSKRLNTIL  120
             MN AG KWSAF +EQ    A+ Y  L++IQN T+K QLQ LQQ+ S VLSE+KSKRLNTIL
Sbjct  61   KMNEAGAKWSAFYEEQCKRAKDYRLEDIQNATIKRQLQILQQSASPVLSEEKSKRLNTIL  120

Query  121  NTMSTIYSTGKVCNPDNPQECLLLEPGLNEIMANSLDYNERLWAWESWRSEVGKQLRPLY  180
            NTMSTIYSTGKVC P+NP+ECLLLEPGL+ IMA+S DYNERLWAWE WRSEVGKQLRP Y
Sbjct  121  NTMSTIYSTGKVCKPNNPEECLLLEPGLDNIMASSTDYNERLWAWEGWRSEVGKQLRPFY  180

Query  181  EEYVVLKNEMARANHYEDYGDYWRGDYEVNGVDGYDYSRGQLIEDVEHTFEEIKPLYEHL  240
            EEYV LKNEMAR   YEDYGDYWR DYE   G  G+ YSR QL+ DVE  FEEIKPLY   L
Sbjct  181  EEYVALKNEMARGYQYEDYGDYWRSDYETEGTSGFQYSRDQLMRDVERIFEEIKPLYVQL  240

Query  241  HAYVRAKLMNAYPSYISPIGCLPAHLLGDMWGRFWTNLYSLTVPFGQKPNIDVTDAMVDQ  300
            HAYVR+KLM+ YPS+ISP G LPAHLLGDMWGRFWTNLY LTVP+GQKPNIDVTDAMV+Q
Sbjct  241  HAYVRSKLMDTYPSHISPTGGLPAHLLGDMWGRFWTNLYPLTVPYGQKPNIDVTDAMVNQ  300

Query  301  AWDAQRIFKEAEKFFVSVGLPNMTQGFWENSMLTDPGNVQKAVCHPTAWDLGKGDFRILM  360
             WDA++IF+EAEKFFVSVGLPNMT+GFWENSMLT+PG+ +K VCHPTAWDLGKGDFRI M
Sbjct  301  KWDAKKIFQEAEKFFVSVGLPNMTKGFWENSMLTEPGDGRKVVCHPTAWDLGKGDFRIKM  360

Query  361  CTKVTMDDFLTAHHEMGHIQYDMAYAAQPFLLRNGANEGFHEAVGEIMSLSAATPKHLKS  420
            CTKVTM+DFLTAHHEMGHIQYDMAYA QP+LLR+GANEGFHEAVGE+MSLS ATPKHLK+
Sbjct  361  CTKVTMEDFLTAHHEMGHIQYDMAYAIQPYLLRSGANEGFHEAVGEVMSLSVATPKHLKT  420

Query  421  IGLLSPDFQEDNETEINFLLKQALTIVGTLPFTYMLEKWRWMVFKGEIPKDQWMKKWWEM  480
            +GLL PDF EDNETEINFLLKQAL IV TLPFTYMLEKWRWMVF GE+PK++W KKWW+M
Sbjct  421  MGLLPPDFNEDNETEINFLLKQALNIVATLPFTYMLEKWRWMVFNGEVPKEEWTKKWWDM  480

Query  481  KREIVGVVEPVPHDETYCDPASLFHVSNDYSFIRYYTRTLYQFQFQEALCQAAKHEGPLH  540
            KREIVGVVEPV HDETYCDPASLFHV+NDYSFIRYYTRT+++FQFQEALC+ A+HEGPLH
Sbjct  481  KREIVGVVEPVSHDETYCDPASLFHVANDYSFIRYYTRTIFEFQFQEALCKIARHEGPLH  540

Query  541  KCDISNSTEAGQKLFNMLRLGKSEPWTLALENVVGAKNMNVRPLLNYFEPLFTWLKDQNK  600
            KCDISNSTEAG+KL  MLRLGKSEPWT ALE+VVG+ NM+V PLL YF+PLFTWLK+QN
Sbjct  541  KCDISNSTEAGKKLLEMLRLGKSEPWTYALESVVGSTNMDVGPLLRYFDPLFTWLKEQNS  600

Query  601  NSFVGWSTDWSPYA-DQSIKVRISLKSALGDKAYEWNDNEMYLFRSSVAYAMRQYFLKVK  659
            NS VGW+T WSPYA DQSIKVRISL SALG+KAYEWNDNEMYLF++SVAYAMR+YFLKVK
Sbjct  601  NSSVGWNTYWSPYAADQSIKVRISLISALGEKAYEWNDNEMYLFQASVAYAMREYFLKVK  660

Query  660  NQMILFGEEDVRVANLKPRISFNFFVTAPKNVSDIIPRTEVEKAIRMSRSRINDAFRLND  719
            NQ + FGEEDVRV++ KPR+SFNFFVT+P +VSDIIPRTEVE AIRMSRSRINDAFRL+D
Sbjct  661  NQTVHFGEEDVRVSDRKPRVSFNFFVTSPNHVSDIIPRTEVEAAIRMSRSRINDAFRLDD  720

Query  720  NSLEFLGIQPTLGPPNQPPVSIWLIVFGVVMGVIVVGIVILIFTGIRDRKKKNKARSGEN  779
            NSLEFLGIQPTLGPP QPPV+IWLIVFGVVMGV+VV I +LIFTGIRDR+KK++ R+ EN
Sbjct  721  NSLEFLGIQPTLGPPYQPPVTIWLIVFGVVMGVVVVAIGLLIFTGIRDRRKKDQERNEEN  780

Query  780  PYASIDISKGENNPGFQNTDDVQTSF  805
            PY S+D+SKGENNPGFQN DDVQTSF
Sbjct  781  PYPSVDLSKGENNPGFQNGDDVQTSF  806
```


**PREDICTED: angiotensin-converting enzyme 2 [Hipposideros armiger]**

Sequence ID: XP_019522936.1 Length: 806 Number of Matches: 1

See 2 more title(s)

Human vs Bat

Range 1: 1 to 806 GenPept Graphics  Next Match

| NW Score | Identities | Positives | Gaps |
|---|---|---|---|
| 3519 | 649/806(81%) | 721/806(89%) | 1/806(0%) |

```
Query  1    MSSSSWLLLSLVAVTAAQSTIEEQAKTFLDKFNHEAEDLFYQSSLASWNYNTNITEENVQ  60
            MS SSWLLLSLVAV AAQS  E+ AK FLDKFN EAEDL + SSLASW+YNTNIT+ENVQ
Sbjct  1    MSGSSWLLLSLVAVAAAQSNSEDLAKEFLDKFNTEAEDLSHLSSLASWDYNTNITDENVQ  60

Query  61   NMNNAGDKWSAFLKEQSTLAQMYPLQEIQNLTVKLQLQALQQNGSSVLSEDKSKRLNTIL  120
             MN AG KWSAF +EQ    A+ Y  L++IQN T+K QLQ LQQ+ S VLSE+KSKRLNTIL
Sbjct  61   KMNEAGAKWSAFYEEQCKRAKDYRLEDIQNATIKRQLQILQQSASPVLSEEKSKRLNTIL  120

Query  121  NTMSTIYSTGKVCNPDNPQECLLLEPGLNEIMANSLDYNERLWAWESWRSEVGKQLRPLY  180
            NTMSTIYSTGKVC P+NP+ECLLLEPGL+ IMA+S DYNERLWAWE WRSEVGKQLRP Y
Sbjct  121  NTMSTIYSTGKVCKPNNPEECLLLEPGLDNIMASSTDYNERLWAWEGWRSEVGKQLRPFY  180

Query  181  EEYVVLKNEMARANHYEDYGDYWRGDYEVNGVDGYDYSRGQLIEDVEHTFEEIKPLYEHL  240
            EEYV LKNEMAR   YEDYGDYWR DYE   G  G+ YSR QL+ DVE  FEEIKPLY   L
Sbjct  181  EEYVALKNEMARGYQYEDYGDYWRSDYETEGTSGFQYSRDQLMRDVERIFEEIKPLYVQL  240

Query  241  HAYVRAKLMNAYPSYISPIGCLPAHLLGDMWGRFWTNLYSLTVPFGQKPNIDVTDAMVDQ  300
            HAYVR+KLM+ YPS+ISP G LPAHLLGDMWGRFWTNLY LTVP+GQKPNIDVTDAMV+Q
Sbjct  241  HAYVRSKLMDTYPSHISPTGGLPAHLLGDMWGRFWTNLYPLTVPYGQKPNIDVTDAMVNQ  300

Query  301  AWDAQRIFKEAEKFFVSVGLPNMTQGFWENSMLTDPGNVQKAVCHPTAWDLGKGDFRILM  360
             WDA++IF+EAEKFFVSVGLPNMT+GFWENSMLT+PG+ +K VCHPTAWDLGKGDFRI M
Sbjct  301  KWDAKKIFQEAEKFFVSVGLPNMTKGFWENSMLTEPGDGRKVVCHPTAWDLGKGDFRIKM  360

Query  361  CTKVTMDDFLTAHHEMGHIQYDMAYAAQPFLLRNGANEGFHEAVGEIMSLSAATPKHLKS  420
            CTKVTM+DFLTAHHEMGHIQYDMAYA QP+LLR+GANEGFHEAVGE+MSLS ATPKHLK+
Sbjct  361  CTKVTMEDFLTAHHEMGHIQYDMAYAIQPYLLRSGANEGFHEAVGEVMSLSVATPKHLKT  420

Query  421  IGLLSPDFQEDNETEINFLLKQALTIVGTLPFTYMLEKWRWMVFKGEIPKDQWMKKWWEM  480
            +GLL PDF EDNETEINFLLKQAL IV TLPFTYMLEKWRWMVF GE+PK++W KKWW+M
Sbjct  421  MGLLPPDFNEDNETEINFLLKQALNIVATLPFTYMLEKWRWMVFNGEVPKEEWTKKWWDM  480

Query  481  KREIVGVVEPVPHDETYCDPASLFHVSNDYSFIRYYTRTLYQFQFQEALCQAAKHEGPLH  540
            KREIVGVVEPV HDETYCDPASLFHV+NDYSFIRYYTRT+++FQFQEALC+ A+HEGPLH
Sbjct  481  KREIVGVVEPVSHDETYCDPASLFHVANDYSFIRYYTRTIFEFQFQEALCKIARHEGPLH  540

Query  541  KCDISNSTEAGQKLFNMLRLGKSEPWTLALENVVGAKNMNVRPLLNYFEPLFTWLKDQNK  600
            KCDISNSTEAG+KL  MLRLGKSEPWT ALE+VVG+ NM+V PLL YF+PLFTWLK+QN
Sbjct  541  KCDISNSTEAGKKLLEMLRLGKSEPWTYALESVVGSTNMDVGPLLRYFDPLFTWLKEQNS  600

Query  601  NSFVGWSTDWSPYA-DQSIKVRISLKSALGDKAYEWNDNEMYLFRSSVAYAMRQYFLKVK  659
            NS VGW+T WSPYA DQSIKVRISL SALG+KAYEWNDNEMYLF++SVAYAMR+YFLKVK
Sbjct  601  NSSVGWNTYWSPYAADQSIKVRISLISALGEKAYEWNDNEMYLFQASVAYAMREYFLKVK  660

Query  660  NQMILFGEEDVRVANLKPRISFNFFVTAPKNVSDIIPRTEVEKAIRMSRSRINDAFRLND  719
            NQ + FGEEDVRV++ KPR+SFNFFVT+P +VSDIIPRTEVE AIRMSRSRINDAFRL+D
Sbjct  661  NQTVHFGEEDVRVSDRKPRVSFNFFVTSPNHVSDIIPRTEVEAAIRMSRSRINDAFRLDD  720

Query  720  NSLEFLGIQPTLGPPNQPPVSIWLIVFGVVMGVIVVGIVILIFTGIRDRKKKNKARSGEN  779
            NSLEFLGIQPTLGPP QPPV+IWLIVFGVVMGV+VV I +LIFTGIRDR+KK++ R+ EN
Sbjct  721  NSLEFLGIQPTLGPPYQPPVTIWLIVFGVVMGVVVVAIGLLIFTGIRDRRKKDQERNEEN  780

Query  780  PYASIDISKGENNPGFQNTDDVQTSF  805
            PY S+D+SKGENNPGFQN DDVQTSF
Sbjct  781  PYPSVDLSKGENNPGFQNGDDVQTSF  806
```

**PREDICTED: angiotensin-converting enzyme 2 [Hipposideros armiger]**

**Sequence ID:** XP_019522936.1 **Length:** 806 **Number of Matches:** 1

#### Human vs Bat

See 2 more title(s)

**Range 1: 1 to 806** GenPept Graphics Next Match

| NW Score | Identities | Positives | Gaps |
|---|---|---|---|
| 3519 | 649/806(81%) | 721/806(89%) | 1/806(0%) |

```
Query  1    MSSSSWLLLSLVAVTAAQSTIEEQAKTFLDKFNHEAEDLFYQSSLASWNYNTNITEENVQ  60
            MS SSWLLLSLVAV AAQS  E+ AK FLDKFN EAEDL + SSLASW+YNTNIT+ENVQ
Sbjct  1    MSGSSWLLLSLVAVAAAQSNSEDLAKEFLDKFNTEAEDLSHLSSLASWDYNTNITDENVQ  60

Query  61   NMNNAGDKWSAFLKEQSTLAQMYPLQEIQNLTVKLQLQALQQNGSSVLSEDKSKRLNTIL  120
             MN AG KWSAF +EQ   A+ Y L++IQN T+K QLQ LQQ+ S VLSE+KSKRLNTIL
Sbjct  61   KMNEAGAKWSAFYEEQCKRAKDYRLEDIQNATIKRQLQILQQSASPVLSEEKSKRLNTIL  120

Query  121  NTMSTIYSTGKVCNPDNPQECLLLEPGLNEIMANSLDYNERLWAWESWRSEVGKQLRPLY  180
            NTMSTIYSTGKVC P+NP+ECLLLEPGL+ IMA+S DYNERLWAWE WRSEVGKQLRP Y
Sbjct  121  NTMSTIYSTGKVCKPNNPEECLLLEPGLDNIMASSTDYNERLWAWEGWRSEVGKQLRPFY  180

Query  181  EEYVVLKNEMARANHYEDYGDYWRGDYEVNGVDGYDYSRGQLIEDVEHTFEEIKPLYEHL  240
            EEYV LKNEMAR   YEDYGDYWR DYE  G  G+ YSR QL+ DVE  FEEIKPLY  L
Sbjct  181  EEYVALKNEMARGYQYEDYGDYWRSDYETEGTSGFQYSRDQLMRDVERIFEEIKPLYVQL  240

Query  241  HAYVRAKLMNAYPSYISPIGCLPAHLLGDMWGRFWTNLYSLTVPFGQKPNIDVTDAMVDQ  300
            HAYVR+KLM+ YPS+ISP G LPAHLLGDMWGRFWTNLY LTVP+GQKPNIDVTDAMV+Q
Sbjct  241  HAYVRSKLMDTYPSHISPTGGLPAHLLGDMWGRFWTNLYPLTVPYGQKPNIDVTDAMVNQ  300

Query  301  AWDAQRIFKEAEKFFVSVGLPNMTQGFWENSMLTDPGNVQKAVCHPTAWDLGKGDFRILM  360
             WDA++IF+EAEKFFVSVGLPNMT+GFWENSMLT+PG+ +K VCHPTAWDLGKGDFRI M
Sbjct  301  KWDAKKIFQEAEKFFVSVGLPNMTKGFWENSMLTEPGDGRKVVCHPTAWDLGKGDFRIKM  360

Query  361  CTKVTMDDFLTAHHEMGHIQYDMAYAAQPFLLRNGANEGFHEAVGEIMSLSAATPKHLKS  420
            CTKVTM+DFLTAHHEMGHIQYDMAYA QP+LLR+GANEGFHEAVGE+MSLS ATPKHLK+
Sbjct  361  CTKVTMEDFLTAHHEMGHIQYDMAYAIQPYLLRSGANEGFHEAVGEVMSLSVATPKHLKT  420

Query  421  IGLLSPDFQEDNETEINFLLKQALTIVGTLPFTYMLEKWRWMVFKGEIPKDQWMKKWWEM  480
            +GLL PDF EDNETEINFLLKQAL IV TLPFTYMLEKWRWMVF GE+PK++W KKWW+M
Sbjct  421  MGLLPPDFNEDNETEINFLLKQALNIVATLPFTYMLEKWRWMVFNGEVPKEEWTKKWWDM  480

Query  481  KREIVGVVEPVPHDETYCDPASLFHVSNDYSFIRYYTRTLYQFQFQEALCQAAKHEGPLH  540
            KREIVGVVEPV HDETYCDPASLFHV+NDYSFIRYYTRT+++FQFQEALC+ A+HEGPLH
Sbjct  481  KREIVGVVEPVSHDETYCDPASLFHVANDYSFIRYYTRTIFEFQFQEALCKIARHEGPLH  540

Query  541  KCDISNSTEAGQKLFNMLRLGKSEPWTLALENVVGAKNMNVRPLLNYFEPLFTWLKDQNK  600
            KCDISNSTEAG+KL  MLRLGKSEPWT ALE+VVG+ NM+V PLL YF+PLFTWLK+QN
Sbjct  541  KCDISNSTEAGKKLLEMLRLGKSEPWTYALESVVGSTNMDVGPLLRYFDPLFTWLKEQNS  600

Query  601  NSFVGWSTDWSPYA-DQSIKVRISLKSALGDKAYEWNDNEMYLFRSSVAYAMRQYFLKVK  659
            NS VGW+T WSPYA DQSIKVRISL SALG+KAYEWNDNEMYLF++SVAYAMR+YFLKVK
Sbjct  601  NSSVGWNTYWSPYAADQSIKVRISLISALGEKAYEWNDNEMYLFQASVAYAMREYFLKVK  660

Query  660  NQMILFGEEDVRVANLKPRISFNFFVTAPKNVSDIIPRTEVEKAIRMSRSRINDAFRLND  719
            NQ + FGEEDVRV++ KPR+SFNFFVT+P +VSDIIPRTEVE AIRMSRSRINDAFRL+D
Sbjct  661  NQTVHFGEEDVRVSDRKPRVSFNFFVTSPNHVSDIIPRTEVEAAIRMSRSRINDAFRLDD  720

Query  720  NSLEFLGIQPTLGPPNQPPVSIWLIVFGVVMGVIVVGIVILIFTGIRDRKKKNKARSGEN  779
            NSLEFLGIQPTLGPP QPPV+IWLIVFGVVMGV+VV I +LIFTGIRDR+KK++ R+ EN
Sbjct  721  NSLEFLGIQPTLGPPYQPPVTIWLIVFGVVMGVVVVAIGLLIFTGIRDRRKKDQERNEEN  780

Query  780  PYASIDISKGENNPGFQNTDDVQTSF  805
            PY S+D+SKGENNPGFQN DDVQTSF
Sbjct  781  PYPSVDLSKGENNPGFQNGDDVQTSF  806
```

**PREDICTED: angiotensin-converting enzyme 2 [Manis javanica]**

Sequence ID: XP_017505746.1 Length: 805 Number of Matches: 1

**Human vs Pan.**

See 1 more title(s)

Range 1: 1 to 805 GenPept Graphics Next Match

| NW Score | Identities | Positives | Gaps |
|---|---|---|---|
| 3679 | 683/805(85%) | 735/805(91%) | 0/805(0%) |

```
Query  1    MSSSSWLLLSLVAVTAAQSTIEEQAKTFLDKFNHEAEDLFYQSSLASWNYNTNITEENVQ  60
            MS SSWLLLSLVAVTAAQST +E+AKTFL+KFN EAE+L YQSSLASWNYNTNIT+ENVQ
Sbjct  1    MSGSSWLLLSLVAVTAAQSTSDEEAKTFLEKFNSEAEELSYQSSLASWNYNTNITDENVQ  60

Query  61   NMNNAGDKWSAFLKEQSTLAQMYPLQEIQNLTVKLQLQALQQNGSSVLSEDKSKRLNTIL  120
             MN AG KWS F +EQS +A+ Y LQ IQN T+K QLQALQ +GSS LS DK++RLNTIL
Sbjct  61   KMNVAGAKWSTFYEEQSKIAKNYQLQNIQNDTIKRQLQALQLSGSSALSADKNQRLNTIL  120

Query  121  NTMSTIYSTGKVCNPDNPQECLLLEPGLNEIMANSLDYNERLWAWESWRSEVGKQLRPLY  180
            NTMSTIYSTGKVCNP NPQEC LLEPGL+ IM +S DYNERLWAWE WRSEVGKQLRPLY
Sbjct  121  NTMSTIYSTGKVCNPGNPQECSLLEPGLDNIMESSKDYNERLWAWEGWRSEVGKQLRPLY  180

Query  181  EEYVVLKNEMARANHYEDYGDYWRGDYEVNGVDGYDYSRGQLIEDVEHTFEEIKPLYEHL  240
            EEYVVLKNEMARANHYEDYGDYWRGDYE  G +GY+YSR  LIEDVEH F +IKPLYEHL
Sbjct  181  EEYVVLKNEMARANHYEDYGDYWRGDYEAEGANGYNYSRDHLIEDVEHIFTQIKPLYEHL  240

Query  241  HAYVRAKLMNAYPSYISPIGCLPAHLLGDMWGRFWTNLYSLTVPFGQKPNIDVTDAMVDQ  300
            HAYVRAKLM+ YPS+ISP GCLPAHLLGDMWGRFWTNLY LTVPF QKPNIDVTDAMV+Q
Sbjct  241  HAYVRAKLMDNYPSHISPTGCLPAHLLGDMWGRFWTNLYPLTVPFRQKPNIDVTDAMVNQ  300

Query  301  AWDAQRIFKEAEKFFVSVGLPNMTQGFWENSMLTDPGNVQKAVCHPTAWDLGKGDFRILM  360
             WDA RIFKEAEKFFVSVGLP MTQ FWENSMLT+PG+ +K VCHPTAWDLGK DFRI M
Sbjct  301  TWDANRIFKEAEKFFVSVGLPKMTQTFWENSMLTEPGDGRKVVCHPTAWDLGKHDFRIKM  360

Query  361  CTKVTMDDFLTAHHEMGHIQYDMAYAAQPFLLRNGANEGFHEAVGEIMSLSAATPKHLKS  420
            CTKVTMDDFLTAHHEMGHIQYDMAYA QP+LLRNGANEGFHEAVGEIMSLSAATPKHLK+
Sbjct  361  CTKVTMDDFLTAHHEMGHIQYDMAYAMQPYLLRNGANEGFHEAVGEIMSLSAATPKHLKN  420

Query  421  IGLLSPDFQEDNETEINFLLKQALTIVGTLPFTYMLEKWRWMVFKGEIPKDQWMKKWWEM  480
            IGLL PDF EDNETEINFLLKQALTIVGTLPFTYMLEKWRWMVF G+IPK+QWMKKWWEM
Sbjct  421  IGLLPPDFYEDNETEINFLLKQALTIVGTLPFTYMLEKWRWMVFSGQIPKEQWMKKWWEM  480

Query  481  KREIVGVVEPVPHDETYCDPASLFHVSNDYSFIRYYTRTLYQFQFQEALCQAAKHEGPLH  540
            KREIVGVVEPVPHDETYCDPASLFHV+NDYSFIRYYTRT+YQFQFQEALCQ AKHEGPLH
Sbjct  481  KREIVGVVEPVPHDETYCDPASLFHVANDYSFIRYYTRTIYQFQFQEALCQTAKHEGPLH  540

Query  541  KCDISNSTEAGQKLFNMLRLGKSEPWTLALENVVGAKNMNVRPLLNYFEPLFTWLKDQNK  600
            KCDISNS EAGQKL  ML LGKS+PWTLALE VVG KNM+VRPLLNYFEPL TWLK+QNK
Sbjct  541  KCDISNSAEAGQKLLQMLSLGKSKPWTLALERVVGTKNMDVRPLLNYFEPLLTWLKEQNK  600

Query  601  NSFVGWSTDWSPYADQSIKVRISLKSALGDKAYEWNDNEMYLFRSSVAYAMRQYFLKVKN  660
            NSFVGW+TDWSPYA QSIKVRISLKSALG+KAYEWND+EMYLFRSSVAYAMR+YF KVK
Sbjct  601  NSFVGWNTDWSPYAAQSIKVRISLKSALGEKAYEWNDSEMYLFRSSVAYAMREYFSKVKK  660

Query  661  QMILFGEEDVRVANLKPRISFNFFVTAPKNVSDIIPRTEVEKAIRMSRSRINDAFRLNDN  720
            Q I F +E VRV++LKPR+SF FFVT PKNVS +IPR EVE+AIR+SRSRINDAFRL+DN
Sbjct  661  QTIPFEDECVRVSDLKPRVSFIFFVTLPKNVSAVIPRAEVEEAIRISRSRINDAFRLDDN  720

Query  721  SLEFLGIQPTLGPPNQPPVSIWLIVFGVVMGVIVVGIVILIFTGIRDRKKKNKARSGENP  780
            SLEFLGIQPTL PP QPPV+IWLIVFGVVMGV+VVGIV+LIFTGIRDRKKK++ARS +NP
Sbjct  721  SLEFLGIQPTLQPPYQPPVTIWLIVFGVVMGVVVVGIVVLIFTGIRDRKKKDQARSEQNP  780

Query  781  YASIDISKGENNPGFQNTDDVQTSF  805
            YAS+D+SKGENNPGFQN DDVQTSF
Sbjct  781  YASVDLSKGENNPGFQNVDDVQTSF  805
```

**PREDICTED: angiotensin-converting enzyme 2 [Manis javanica]**

Sequence ID: XP_017505746.1 Length: 805 Number of Matches: 1

**Human vs Pan.**

See 1 more title(s)

**PREDICTED: angiotensin-converting enzyme 2 [Manis javanica]**

Sequence ID: XP_017505752.1

Range 1: 1 to 805 GenPept Graphics Next Match

| NW Score | Identities | Positives | Gaps |
|---|---|---|---|
| 3679 | 683/805(85%) | 735/805(91%) | 0/805(0%) |

```
Query  1    MSSSSWLLLSLVAVTAAQSTIEEQAKTFLDKFNHEAEDLFYQSSLASWNYNTNITEENVQ  60
            MS SSWLLLSLVAVTAAQST +E+AKTFL+KFN EAE+L YQSSLASWNYNTNIT+ENVQ
Sbjct  1    MSGSSWLLLSLVAVTAAQSTSDEEAKTFLEKFNSEAEELSYQSSLASWNYNTNITDENVQ  60

Query  61   NMNNAGDKWSAFLKEQSTLAQMYPLQEIQNLTVKLQLQALQQNGSSVLSEDKSKRLNTIL  120
             MN AG KWS F +EQS +A+ Y LQ IQN T+K QLQALQ +GSS LS DK++RLNTIL
Sbjct  61   KMNVAGAKWSTFYEEQSKIAKNYQLQNIQNDTIKRQLQALQLSGSSALSADKNQRLNTIL  120

Query  121  NTMSTIYSTGKVCNPDNPQECLLLEPGLNEIMANSLDYNERLWAWESWRSEVGKQLRPLY  180
            NTMSTIYSTGKVCNP NPQEC LLEPGL+ IM +S DYNERLWAWE WRSEVGKQLRPLY
Sbjct  121  NTMSTIYSTGKVCNPGNPQECSLLEPGLDNIMESSKDYNERLWAWEGWRSEVGKQLRPLY  180

Query  181  EEYVVLKNEMARANHYEDYGDYWRGDYEVNGVDGYDYSRGQLIEDVEHTFEEIKPLYEHL  240
            EEYVVLKNEMARANHYEDYGDYWRGDYE  G +GY+YSR  LIEDVEH F +IKPLYEHL
Sbjct  181  EEYVVLKNEMARANHYEDYGDYWRGDYEAEGANGYNYSRDHLIEDVEHIFTQIKPLYEHL  240

Query  241  HAYVRAKLMNAYPSYISPIGCLPAHLLGDMWGRFWTNLYSLTVPFGQKPNIDVTDAMVDQ  300
            HAYVRAKLM+ YPS+ISP GCLPAHLLGDMWGRFWTNLY LTVPF QKPNIDVTDAMV+Q
Sbjct  241  HAYVRAKLMDNYPSHISPTGCLPAHLLGDMWGRFWTNLYPLTVPFRQKPNIDVTDAMVNQ  300

Query  301  AWDAQRIFKEAEKFFVSVGLPNMTQGFWENSMLTDPGNVQKAVCHPTAWDLGKGDFRILM  360
             WDA RIFKEAEKFFVSVGLP MTQ FWENSMLT+PG+ +K VCHPTAWDLGK DFRI M
Sbjct  301  TWDANRIFKEAEKFFVSVGLPKMTQTFWENSMLTEPGDGRKVVCHPTAWDLGKHDFRIKM  360

Query  361  CTKVTMDDFLTAHHEMGHIQYDMAYAAQPFLLRNGANEGFHEAVGEIMSLSAATPKHLKS  420
            CTKVTMDDFLTAHHEMGHIQYDMAYA QP+LLRNGANEGFHEAVGEIMSLSAATPKHLK+
Sbjct  361  CTKVTMDDFLTAHHEMGHIQYDMAYAMQPYLLRNGANEGFHEAVGEIMSLSAATPKHLKN  420

Query  421  IGLLSPDFQEDNETEINFLLKQALTIVGTLPFTYMLEKWRWMVFKGEIPKDQWMKKWWEM  480
            IGLL PDF EDNETEINFLLKQALTIVGTLPFTYMLEKWRWMVF G+IPK+QWMKKWWEM
Sbjct  421  IGLLPPDFYEDNETEINFLLKQALTIVGTLPFTYMLEKWRWMVFSGQIPKEQWMKKWWEM  480

Query  481  KREIVGVVEPVPHDETYCDPASLFHVSNDYSFIRYYTRTLYQFQFQEALCQAAKHEGPLH  540
            KREIVGVVEPVPHDETYCDPASLFHV+NDYSFIRYYTRT+YQFQFQEALCQ AKHEGPLH
Sbjct  481  KREIVGVVEPVPHDETYCDPASLFHVANDYSFIRYYTRTIYQFQFQEALCQTAKHEGPLH  540

Query  541  KCDISNSTEAGQKLFNMLRLGKSEPWTLALENVVGAKNMNVRPLLNYFEPLFTWLKDQNK  600
            KCDISNS EAGQKL  ML LGKS+PWTLALE VVG KNM+VRPLLNYFEPL TWLK+QNK
Sbjct  541  KCDISNSAEAGQKLLQMLSLGKSKPWTLALERVVGTKNMDVRPLLNYFEPLLTWLKEQNK  600

Query  601  NSFVGWSTDWSPYADQSIKVRISLKSALGDKAYEWNDNEMYLFRSSVAYAMRQYFLKVKN  660
            NSFVGW+TDWSPYA QSIKVRISLKSALG+KAYEWND+EMYLFRSSVAYAMR+YF KVK
Sbjct  601  NSFVGWNTDWSPYAAQSIKVRISLKSALGEKAYEWNDSEMYLFRSSVAYAMREYFSKVKK  660

Query  661  QMILFGEEDVRVANLKPRISFNFFVTAPKNVSDIIPRTEVEKAIRMSRSRINDAFRLNDN  720
            Q I F +E VRV++LKPR+SF FFVT PKNVS +IPR EVE+AIR+SRSRINDAFRL+DN
Sbjct  661  QTIPFEDECVRVSDLKPRVSFIFFVTLPKNVSAVIPRAEVEEAIRISRSRINDAFRLDDN  720

Query  721  SLEFLGIQPTLGPPNQPPVSIWLIVFGVVMGVIVVGIVILIFTGIRDRKKKNKARSGENP  780
            SLEFLGIQPTL PP QPPV+IWLIVFGVVMGV+VVGIV+LIFTGIRDRKKK++ARS +NP
Sbjct  721  SLEFLGIQPTLQPPYQPPVTIWLIVFGVVMGVVVVGIVVLIFTGIRDRKKKDQARSEQNP  780

Query  781  YASIDISKGENNPGFQNTDDVQTSF  805
            YAS+D+SKGENNPGFQN DDVQTSF
Sbjct  781  YASVDLSKGENNPGFQNVDDVQTSF  805
```

#### LOW QUALITY PROTEIN: angiotensin-converting enzyme 2 [Notechis scutatus]

Sequence ID: XP_026530754.1 Length: 828 Number of Matches: 1

**Human vs Snake**

Range 1: 1 to 828 GenPept Graphics Next Match

| NW Score | Identities | Positives | Gaps |
|---|---|---|---|
| 2608 | 484/832(58%) | 627/832(75%) | 31/832(3%) |

```
Query  1    MSSS-----------------------SWLLL--SLVAVTAAQSTIEEQAKTFLDKFNHE  35
            M  +                       SWL L  SLV +   AQ    ++ A+ FL +F+
Sbjct  1    MKQALVRKPSSRSFTHPAFSDXKGNMLSWLCLTWSLVVLALAQDETQKAAE-FLKQFDIR  59

Query  36   AEDLFYQSSLASWNYNTNITEENVQNMNNAGDKWSAFLKEQSTLAQMYPLQEIQNLTVKL  95
            A DL+Y +S+ASWNYNTN+TEEN + M+      +S F   E S  A M+ + +I N T+KL
Sbjct  60   AVDLYYNASIASWNYNTNLTEENAKIMHEKDSIFSRFYDEASRNASMFNVNQISNETIKL  119

Query  96   QLQALQQNGSSVLSEDKSKRLNTILNTMSTIYSTGKVCNPDNPQECLLLEPGLNEIMANS  155
            Q++ LQ    +   ++D+   L+T+L  MST+YSTG VC  D+P  CL LEPGL+ IMAN+
Sbjct  120  QIRLLQNGPTDSSTKDQ---LDTVLRKMSTLYSTGTVCKQDDPFNCLPLEPGLDHIMANN  176

Query  156  LDYNERLWAWESWRSEVGKQLRPLYEEYVVLKNEMARANHYEDYGDYWRGDYEVNGVDGY  215
             +Y+ERLWAWESWR++VGK++RPLYE YV LKN+ AR   Y+DYGDYWR +YEV+     +
Sbjct  177  WNYSERLWAWESWRADVGKKMRPLYETYVELKNKYARLRGYDDYGDYWRANYEVDLPGKF  236

Query  216  DYSRGQLIEDVEHTFEEIKPLYEHLHAYVRAKLMNAY-PSYISPIGCLPAHLLGDMWGRF  274
             Y R QLI DVE+TF++I PLYE LHAYVR  L   Y P  I+P G +PAHLLGDMWGRF
Sbjct  237  QYQRAQLITDVENTFKQILPLYEQLHAYVRRHLYKRYGPELINPKGAIPAHLLGDMWGRF  296

Query  275  WTNLYSLTVPFGQKPNIDVTDAMVDQAWDAQRIFKEAEKFFVSVGLPNMTQGFWENSMLT  334
            WTNLY L VP+  K +IDV+ AMV++ W    IFK AE FF+S+GL NMT+ FW+NSML
Sbjct  297  WTNLYPLMVPYPNKTSIDVSSAMVEKKWTVDSIFKAAEHFFISIGLFNMTESFWKNSMLE  356

Query  335  DPGNVQKAVCHPTAWDLGKGDFRILMCTKVTMDDFLTAHHEMGHIQYDMAYAAQPFLLRN  394
            +P + +K VCHPTAWD+GK D+RI MCTK+ M+DFLTAHHEMGHI+YDMAYA QPFLLRN
Sbjct  357  EPKDGRKVVCHPTAWDMGKEDYRIKMCTKINMEDFLTAHHEMGHIEYDMAYADQPFLLRN  416

Query  395  GANEGFHEAVGEIMSLSAATPKHLKSIGLLSPDFQEDNETEINFLLKQALTIVGTLPFTY  454
            GANEGFHEAVGEIMSLSAATPK+L+S+GLL   FQED ET+INFLL+QALTIVGT+PFTY
Sbjct  417  GANEGFHEAVGEIMSLSAATPKYLQSLGLLESTFQEDAETDINFLLRQALTIVGTMPFTY  476

Query  455  MLEKWRWMVFKGEIPKDQWMKKWWEMKREIVGVVEPVPHDETYCDPASLFHVSNDYSFIR  514
            MLEKWRWMVF  +IPKDQWMKKWWEMKREIVGVVEP+PH+E YCDPA+LFHV+NDYSFIR
Sbjct  477  MLEKWRWMVFAEQIPKDQWMKKWWEMKREIVGVVEPLPHNEEYCDPAALFHVANDYSFIR  536

Query  515  YYTRTLYQFQFQEALCQAAKHEGPLHKCDISNSTEAGQKLFNMLRLGKSEPWTLALENVV  574
            YYTRT+YQFQFQEALC+AA H   L+KCDISNST AG+ L +ML LG S+PWT ALE++
Sbjct  537  YYTRTIYQFQFQEALCKAAGHTEELYKCDISNSTNAGRILKDMLALGSSQPWTKALESIT  596

Query  575  GAKNMNVRPLLNYFEPLFTWLKDQNKNSFVGWSTDWSPYADQSIKVRISLKSALGDKAYE  634
            G++ M+ +P   YF+PL  WL+  N N  VGW+ +W+PY+  +IKVRISLK+ALGD AY
Sbjct  597  GSQKMDAKPFCQYFDPLLKWLEKANSNENVGWNVNWTPYSKDAIKVRISLKTALGDDAYN  656

Query  635  WNDNEMYLFRSSVAYAMRQYFLKVKNQMILFGEEDVRVANLKPRISFNFFVTAPKNVSDI  694
            W+++EMYLF+S++AYAM++YFL+VKN+ +LF  ++V V+++  RISF F V+ P N+S++
Sbjct  657  WDESEMYLFKSTIAYAMQKYFLEVKNKTVLFQTDNVHVSDMTVRISFYFTVSMPNNISEL  716

Query  695  IPRTEVEKAIRMSRSRINDAFRLNDNSLEFLGIQPTLGPPNQPPVSIWLIVFGVVMGVIV  754
            +P++EVE+AI +SR RIN+AF+L D +LEF+G+ PTL PP + P+++WLI FGVV+G++V
Sbjct  717  VPKSEVEEAISLSRDRINEAFKLTDQTLEFVGLLPTLAPPYESPITVWLIAFGVVIGLVV  776

Query  755  VGIVILIFTGIRDRKKKNKA-RSGENPYASIDISKGENNPGFQNTDDVQTSF  805
            +GI+ L+  G +DRKKK +A ++     A+I    G++N  FQ  +   T+F
Sbjct  777  IGIITLLIIGQKDRKKKKRAAKTNSMETAAIHEDCGQSNSTFQLDEAATTTF  828
```

#### angiotensin-converting enzyme 2 [Thamnophis elegans]

Sequence ID: XP_032082934.1 Length: 828 Number of Matches: 1

**Human vs Snake**

Range 1: 1 to 828 GenPept Graphics Next Match

| NW Score | Identities | Positives | Gaps |
|---|---|---|---|
| 2592 | 485/832(58%) | 620/832(74%) | 31/832(3%) |

```
Query  1    MSSS-----------------------SWLLL--SLVAVTAAQSTIEEQAKTFLDKFNHE  35
            M  +                       SWL    SLV +   AQ   + +QA  FL +F+
Sbjct  1    MKQALIRKPSSRSFTHPAFFDLKGNMLSWLCFTWSLVVLAGAQD-VTQQAAEFLKQFDAR  59

Query  36   AEDLFYQSSLASWNYNTNITEENVQNMNNAGDKWSAFLKEQSTLAQMYPLQEIQNLTVKL  95
            A+DL+Y +S+ASWNYNTN+TEEN + M+    + +S F +E S  A MY + +I N T++L
Sbjct  60   ADDLYYAASIASWNYNTNLTEENAKIMHEKDNIFSKFYEEASKNASMYNVNQITNETIRL  119

Query  96   QLQALQQNGSSVLSEDKSKRLNTILNTMSTIYSTGKVCNPDNPQECLLLEPGLNEIMANS  155
            QL  LQ    ++   ++D+   L+T+L  MST+YSTG VC  D+P  CL LEPGL++IM N+
Sbjct  120  QLHLLQNVPTNSSTKDQ---LDTVLRKMSTMYSTGTVCKQDDPFNCLPLEPGLDDIMENN  176

Query  156  LDYNERLWAWESWRSEVGKQLRPLYEEYVVLKNEMARANHYEDYGDYWRGDYEVNGVDGY  215
              Y+ERLWAWE WR++VGK++RPLYE YV LKN+ AR   Y DYGDYWR +YEV+     Y
Sbjct  177  WSYSERLWAWEGWRADVGKKMRPLYESYVELKNKYARLRGYADYGDYWRANYEVDLPKEY  236

Query  216  DYSRGQLIEDVEHTFEEIKPLYEHLHAYVRAKLMNAY-PSYISPIGCLPAHLLGDMWGRF  274
             Y R QLI DVE+T ++I PLY+HLHAYVR  L   Y P +I+  G +PAHLLGDMWGRF
Sbjct  237  QYQRAQLITDVENTLQQIMPLYKHLHAYVRRHLYKHYGPEFINLEGAIPAHLLGDMWGRF  296

Query  275  WTNLYSLTVPFGQKPNIDVTDAMVDQAWDAQRIFKEAEKFFVSVGLPNMTQGFWENSMLT  334
            WTNLY L VPF  K +IDVT AMV + W    IFK AE+FF S+GL  MT  FW NSML
Sbjct  297  WTNLYPLMVPFPNKTSIDVTSAMVTKKWTVNSIFKAAEQFFTSIGLFPMTDNFWNNSMLE  356

Query  335  DPGNVQKAVCHPTAWDLGKGDFRILMCTKVTMDDFLTAHHEMGHIQYDMAYAAQPFLLRN  394
            +P + +K VCHPTAWD+GK D+RI MCTK+ M+DFLTAHHEMGHI+YDMAY+ QPFLLRN
Sbjct  357  EPKDGRKVVCHPTAWDMGKKDYRIKMCTKINMEDFLTAHHEMGHIEYDMAYSDQPFLLRN  416

Query  395  GANEGFHEAVGEIMSLSAATPKHLKSIGLLSPDFQEDNETEINFLLKQALTIVGTLPFTY  454
            GANEGFHEAVGEIMSLSAATPK+LKS+GLL   FQED ET+INFLLKQALTIVGT+PFTY
Sbjct  417  GANEGFHEAVGEIMSLSAATPKYLKSLGLLEHTFQEDTETDINFLLKQALTIVGTMPFTY  476

Query  455  MLEKWRWMVFKGEIPKDQWMKKWWEMKREIVGVVEPVPHDETYCDPASLFHVSNDYSFIR  514
            MLEKWRWMVF  +IPKDQWMKKWWEMKREIVGVVEP+PH+E YCDPA+LFHV+NDYSFIR
Sbjct  477  MLEKWRWMVFAEQIPKDQWMKKWWEMKREIVGVVEPLPHNEEYCDPAALFHVANDYSFIR  536

Query  515  YYTRTLYQFQFQEALCQAAKHEGPLHKCDISNSTEAGQKLFNMLRLGKSEPWTLALENVV  574
            YYTRT+YQFQFQEALCQAA H G L+KC+IS+ST+AG  L +ML LG S+PWT ALE++
Sbjct  537  YYTRTIYQFQFQEALCQAAGHTGELYKCEISHSTDAGHILKDMLALGSSQPWTKALESIT  596

Query  575  GAKNMNVRPLLNYFEPLFTWLKDQNKNSFVGWSTDWSPYADQSIKVRISLKSALGDKAYE  634
             ++ M+  P  +YF+PL  WL+ QN N  VGW+ +W+PY+  +IKVRISLK ALGD AY
Sbjct  597  KSQKMDATPFRHYFDPLLKWLEKQNSNENVGWNVNWTPYSKYAIKVRISLKRALGDDAYN  656

Query  635  WNDNEMYLFRSSVAYAMRQYFLKVKNQMILFGEEDVRVANLKPRISFNFFVTAPKNVSDI  694
            W  +EMYLF+S++AYAM++YFL++KN+ +LF  ++V V+ +  RISF F V+ P N+S++
Sbjct  657  WTASEMYLFKSTIAYAMQKYFLEIKNKTVLFQTDNVHVSPVTERISFYFTVSMPTNISEL  716

Query  695  IPRTEVEKAIRMSRSRINDAFRLNDNSLEFLGIQPTLGPPNQPPVSIWLIVFGVVMGVIV  754
            +P++EVE+AI +SR RIN+AFRL D +LEF+G+ PTL PP + P+++WL+VFGVV+G++V
Sbjct  717  VPKSEVEEAISLSRDRINEAFRLTDQTLEFVGLLPTLAPPYESPITVWLVVFGVVIGIVV  776

Query  755  VGIVILIFTGIRDRKKKNKA-RSGENPYASIDISKGENNPGFQNTDDVQTSF  805
            +GI+ LI  G +DRKKK +A ++     A+I    G++N  FQ  +   T+F
Sbjct  777  IGIITLIIIGQKDRKKKKQAAKTDAIETAAIHEDCGQSNSTFQLDEAATTTF  828
```

Hayashi *et al.* Figure 2

#### Global Alignment » results for RID-8B0TUUHX114 Human ACE2 vs Mammals ACE2

| Species | | | Protein ID | Homology |
|---|---|---|---|---|
| Human | [Homo sapiens] | NCBI Reference Sequence: | NP_001358344.1 | |
| Mouse | [Mus musculus] | NCBI Reference Sequence: | NP_081562.2 | 89.0% |
| Mouse | [Mus musculus] | NCBI Reference Sequence: | NP_001123985.1 | 89.0% |
| Mouse | [Mus musculus] | NCBI Reference Sequence: | XP_028743609.1 | 89.0% |
| Mouse | [Mus musculus] | NCBI Reference Sequence: | XP_021009138.1 | 89.0% |
| Mouse | [Mus musculus] | NCBI Reference Sequence: | XP_021043935.1 | 90.0% |
| Mouse | [Mus musculus] | NCBI Reference Sequence: | XP_031226742.1 | 90.0% |
| Mouse | [Mus musculus] | NCBI Reference Sequence: | XP_006973269.1 | 89.0% |
| Mouse | [Mus musculus] | NCBI Reference Sequence: | XP_012585871.1 | 86.0% |
| Rat | [Rattus norvegicus] | NCBI Reference Sequence: | NP_001012006.1 | 90.0% |
| Rat | [Rattus norvegicus] | NCBI Reference Sequence: | XP_004866157.1 | 91.0% |
| Rat | [Rattus norvegicus] | NCBI Reference Sequence: | XP_012887572.1 | 89.0% |
| Rat | [Dipodomys ordii] | NCBI Reference Sequence: | XP_012887573.1 | 89.0% |
| Rat | [Rattus rattus] | NCBI Reference Sequence: | XP_032746145.1 | 87.0% |
| Rat | [Fukomys damarensis] | NCBI Reference Sequence: | XP_010643477.1 | 91.0% |
| Rat | [Nannospalax galili] | NCBI Reference Sequence: | XP_008839098.1 | 91.0% |
| Hamster | [Cricetulus griseus] | NCBI Reference Sequence: | XP_003503283.1 | 91.0% |
| Hamster | [Cricetulus griseus] | NCBI Reference Sequence: | XP_027288607.1 | 91.0% |
| Hamster | [Mesocricetus auratus] | NCBI Reference Sequence: | XP_005074266.1 | 91.0% |
| Mink | [Neovison vison] | NCBI Reference Sequence: | CCP86723.1 | 93.0% |
| Salmo | [Salmo salar ] | NCBI Reference Sequence: | XP_014062928.1 | 74.0% |

https://blast.ncbi.nlm.nih.gov/Blast.cgi

**angiotensin-converting enzyme 2 precursor [Mus musculus]**

**Sequence ID:** NP_001123985.1 **Length:** 805 **Number of Matches:** 1

#### Human vs Mouse

See 7 more title(s)

**Range 1: 12 to 805** GenPept Graphics ▼ Next Match ▲

| Score | Expect | Method | Identities | Positives | Gaps |
|---|---|---|---|---|---|
| 1369 bits(3543) | 0.0 | Compositional matrix adjust. | 650/794(82%) | 710/794(89%) | 0/794(0%) |

```
Query  12   VAVTAAQSTIEEQAKTFLDKFNHEAEDLFYQSSLASWNYNTNITEENVQNMNNAGDKWSA  71
            VAVT AQS  EE AKTFL+ FN EAEDL YQSSLASWNYNTNITEEN Q M+ A  KWSA
Sbjct  12   VAVTTAQSLTEENAKTFLNNFNQEAEDLSYQSSLASWNYNTNITEENAQKMSEAAAKWSA  71

Query  72   FLKEQSTLAQMYPLQEIQNLTVKLQLQALQQNGSSVLSEDKSKRLNTILNTMSTIYSTGK  131
            F +EQS  AQ + LQEIQ   +K QLQALQQ+GSS LS DK+K+LNTILNTMSTIYSTGK
Sbjct  72   FYEEQSKTAQSFSLQEIQTPIIKRQLQALQQSGSSALSADKNKQLNTILNTMSTIYSTGK  131

Query  132  VCNPDNPQECLLLEPGLNEIMANSLDYNERLWAWESWRSEVGKQLRPLYEEYVVLKNEMA  191
            VCNP NPQECLLLEPGL+EIMA S DYN RLWAWE WR+EVGKQLRPLYEEYVVLKNEMA
Sbjct  132  VCNPKNPQECLLLEPGLDEIMATSTDYNSRLWAWEGWRAEVGKQLRPLYEEYVVLKNEMA  191

Query  192  RANHYEDYGDYWRGDYEVNGVDGYDYSRGQLIEDVEHTFEEIKPLYEHLHAYVRAKLMNA  251
            RAN+Y DYGDYWRGDYE  G DGY+Y+R QLIEDVE TF EIKPLYEHLHAYVR KLM+
Sbjct  192  RANNYNDYGDYWRGDYEAEGADGYNYNRNQLIEDVERTFAEIKPLYEHLHAYVRRKLMDT  251

Query  252  YPSYISPIGCLPAHLLGDMWGRFWTNLYSLTVPFGQKPNIDVTDAMVDQAWDAQRIFKEA  311
            YPSYISP GCLPAHLLGDMWGRFWTNLY LTVPF QKPNIDVTDAM++Q WDA+RIF+EA
Sbjct  252  YPSYISPTGCLPAHLLGDMWGRFWTNLYPLTVPFAQKPNIDVTDAMMNQGWDAERIFQEA  311

Query  312  EKFFVSVGLPNMTQGFWENSMLTDPGNVQKAVCHPTAWDLGKGDFRILMCTKVTMDDFLT  371
            EKFFVSVGLP+MTQGFW NSMLT+P  + +K VCHPTAWDLG GDFRI MCTKVTMD+FLT
Sbjct  312  EKFFVSVGLPHMTQGFWANSMLTEPADGRKVVCHPTAWDLGHGDFRIKMCTKVTMDNFLT  371

Query  372  AHHEMGHIQYDMAYAAQPFLLRNGANEGFHEAVGEIMSLSAATPKHLKSIGLLSPDFQED  431
            AHHEMGHIQYDMAYA QPFLLRNGANEGFHEAVGEIMSLSAATPKHLKSIGLL  DFQED
Sbjct  372  AHHEMGHIQYDMAYARQPFLLRNGANEGFHEAVGEIMSLSAATPKHLKSIGLLPSDFQED  431

Query  432  NETEINFLLKQALTIVGTLPFTYMLEKWRWMVFKGEIPKDQWMKKWWEMKREIVGVVEPV  491
            +ETEINFLLKQALTIVGTLPFTYMLEKWRWMVF+GEIPK+QWMKKWWEMKREIVGVVEP+
Sbjct  432  SETEINFLLKQALTIVGTLPFTYMLEKWRWMVFRGEIPKEQWMKKWWEMKREIVGVVEPL  491

Query  492  PHDETYCDPASLFHVSNDYSFIRYYTRTLYQFQFQEALCQAAKHEGPLHKCDISNSTEAG  551
            PHDETYCDPASLFHVSNDYSFIRYYTRT+YQFQFQEALCQAAK+ G LHKCDISNSTEAG
Sbjct  492  PHDETYCDPASLFHVSNDYSFIRYYTRTIYQFQFQEALCQAAKYNGSLHKCDISNSTEAG  551

Query  552  QKLFNMLRLGKSEPWTLALENVVGAKNMNVRPLLNYFEPLFTWLKDQNKNSFVGWSTDWS  611
            QKL  ML LG SEPWT ALENVVGA+NM+V+PLLNYF+PLF WLK+QN+NSFVGW+T+WS
Sbjct  552  QKLLKMLSLGNSEPWTKALENVVGARNMDVKPLLNYFQPLFDWLKEQNRNSFVGWNTEWS  611

Query  612  PYADQSIKVRISLKSALGDKAYEWNDNEMYLFRSSVAYAMRQYFLKVKNQMILFGEEDVR  671
            PYADQSIKVRISLKSALG  AYEW +NEM+LFRSSVAYAMR+YF  +KNQ + F EEDVR
Sbjct  612  PYADQSIKVRISLKSALGANAYEWTNNEMFLFRSSVAYAMRKYFSIIKNQTVPFLEEDVR  671

Query  672  VANLKPRISFNFFVTAPKNVSDIIPRTEVEKAIRMSRSRINDAFRLNDNSLEFLGIQPTL  731
            V++LKPR+SF FFVT+P+NVSD+IPR+EVE AIRMSR RIND F LNDNSLEFLGI PTL
Sbjct  672  VSDLKPRVSFYFFVTSPQNVSDVIPRSEVEDAIRMSRGRINDVFGLNDNSLEFLGIHPTL  731

Query  732  GPPNQPPVSIWLIVFGVVMGVIVVGIVILIFTGIRDRKKKNKARSGENPYASIDISKGEN  791
             PP QPPV+IWLI+FGVVM ++VVGI+ILI TGI+ RKKKN+ +   ENPY S+DI KGE+
Sbjct  732  EPPYQPPVTIWLIIFGVVMALVVVGIIILIVTGIKGRKKKNETKREENPYDSMDIGKGES  791

Query  792  NPGFQNTDDVQTSF   805
            N GFQN+DD QTSF
Sbjct  792  NAGFQNSDDAQTSF   805
```


**angiotensin-converting enzyme 2 precursor [Mus musculus]**

**Sequence ID:** NP_001123985.1 **Length:** 805 **Number of Matches:** 1

Human vs Mouse

See 7 more title(s)

**Range 1: 12 to 805** GenPept Graphics ▼ Next Match ▲ Pre

| Score | Expect | Method | Identities | Positives | Gaps |
|---|---|---|---|---|---|
| 1369 bits(3543) | 0.0 | Compositional matrix adjust. | 650/794(82%) | 710/794(89%) | 0/794(0%) |

```
Query  12   VAVTAAQSTIEEQAKTFLDKFNHEAEDLFYQSSLASWNYNTNITEENVQNMNNAGDKWSA  71
            VAVT AQS  EE AKTFL+ FN EAEDL YQSSLASWNYNTNITEEN Q M+ A  KWSA
Sbjct  12   VAVTTAQSLTEENAKTFLNNFNQEAEDLSYQSSLASWNYNTNITEENAQKMSEAAAKWSA  71

Query  72   FLKEQSTLAQMYPLQEIQNLTVKLQLQALQQNGSSVLSEDKSKRLNTILNTMSTIYSTGK  131
            F +EQS  AQ + LQEIQ   +K QLQALQQ+GSS LS DK+K+LNTILNTMSTIYSTGK
Sbjct  72   FYEEQSKTAQSFSLQEIQTPIIKRQLQALQQSGSSALSADKNKQLNTILNTMSTIYSTGK  131

Query  132  VCNPDNPQECLLLEPGLNEIMANSLDYNERLWAWESWRSEVGKQLRPLYEEYVVLKNEMA  191
            VCNP NPQECLLLEPGL+EIMA S DYN RLWAWE WR+EVGKQLRPLYEEYVVLKNEMA
Sbjct  132  VCNPKNPQECLLLEPGLDEIMATSTDYNSRLWAWEGWRAEVGKQLRPLYEEYVVLKNEMA  191

Query  192  RANHYEDYGDYWRGDYEVNGVDGYDYSRGQLIEDVEHTFEEIKPLYEHLHAYVRAKLMNA  251
            RAN+Y DYGDYWRGDYE  G DGY+Y+R QLIEDVE TF EIKPLYEHLHAYVR KLM+
Sbjct  192  RANNYNDYGDYWRGDYEAEGADGYNYNRNQLIEDVERTFAEIKPLYEHLHAYVRRKLMDT  251

Query  252  YPSYISPIGCLPAHLLGDMWGRFWTNLYSLTVPFGQKPNIDVTDAMVDQAWDAQRIFKEA  311
            YPSYISP GCLPAHLLGDMWGRFWTNLY LTVPF QKPNIDVTDAM++Q WDA+RIF+EA
Sbjct  252  YPSYISPTGCLPAHLLGDMWGRFWTNLYPLTVPFAQKPNIDVTDAMMNQGWDAERIFQEA  311

Query  312  EKFFVSVGLPNMTQGFWENSMLTDPGNVQKAVCHPTAWDLGKGDFRILMCTKVTMDDFLT  371
            EKFFVSVGLP+MTQGFW NSMLT+P  + +K VCHPTAWDLG GDFRI MCTKVTMD+FLT
Sbjct  312  EKFFVSVGLPHMTQGFWANSMLTEPADGRKVVCHPTAWDLGHGDFRIKMCTKVTMDNFLT  371

Query  372  AHHEMGHIQYDMAYAAQPFLLRNGANEGFHEAVGEIMSLSAATPKHLKSIGLLSPDFQED  431
            AHHEMGHIQYDMAYA QPFLLRNGANEGFHEAVGEIMSLSAATPKHLKSIGLL  DFQED
Sbjct  372  AHHEMGHIQYDMAYARQPFLLRNGANEGFHEAVGEIMSLSAATPKHLKSIGLLPSDFQED  431

Query  432  NETEINFLLKQALTIVGTLPFTYMLEKWRWMVFKGEIPKDQWMKKWWEMKREIVGVVEPV  491
            +ETEINFLLKQALTIVGTLPFTYMLEKWRWMVF+GEIPK+QWMKKWWEMKREIVGVVEP+
Sbjct  432  SETEINFLLKQALTIVGTLPFTYMLEKWRWMVFRGEIPKEQWMKKWWEMKREIVGVVEPL  491

Query  492  PHDETYCDPASLFHVSNDYSFIRYYTRTLYQFQFQEALCQAAKHEGPLHKCDISNSTEAG  551
            PHDETYCDPASLFHVSNDYSFIRYYTRT+YQFQFQEALCQAAK+ G LHKCDISNSTEAG
Sbjct  492  PHDETYCDPASLFHVSNDYSFIRYYTRTIYQFQFQEALCQAAKYNGSLHKCDISNSTEAG  551

Query  552  QKLFNMLRLGKSEPWTLALENVVGAKNMNVRPLLNYFEPLFTWLKDQNKNSFVGWSTDWS  611
            QKL  ML LG SEPWT ALENVVGA+NM+V+PLLNYF+PLF WLK+QN+NSFVGW+T+WS
Sbjct  552  QKLLKMLSLGNSEPWTKALENVVGARNMDVKPLLNYFQPLFDWLKEQNRNSFVGWNTEWS  611

Query  612  PYADQSIKVRISLKSALGDKAYEWNDNEMYLFRSSVAYAMRQYFLKVKNQMILFGEEDVR  671
            PYADQSIKVRISLKSALG  AYEW +NEM+LFRSSVAYAMR+YF  +KNQ + F EEDVR
Sbjct  612  PYADQSIKVRISLKSALGANAYEWTNNEMFLFRSSVAYAMRKYFSIIKNQTVPFLEEDVR  671

Query  672  VANLKPRISFNFFVTAPKNVSDIIPRTEVEKAIRMSRSRINDAFRLNDNSLEFLGIQPTL  731
            V++LKPR+SF FFVT+P+NVSD+IPR+EVE AIRMSR RIND F LNDNSLEFLGI PTL
Sbjct  672  VSDLKPRVSFYFFVTSPQNVSDVIPRSEVEDAIRMSRGRINDVFGLNDNSLEFLGIHPTL  731

Query  732  GPPNQPPVSIWLIVFGVVMGVIVVGIVILIFTGIRDRKKKNKARSGENPYASIDISKGEN  791
             PP QPPV+IWLI+FGVVM ++VVGI+ILI TGI+ RKKKN+ +   ENPY S+DI KGE+
Sbjct  732  EPPYQPPVTIWLIIFGVVMALVVVGIIILIVTGIKGRKKKNETKREENPYDSMDIGKGES  791

Query  792  NPGFQNTDDVQTSF   805
            N GFQN+DD QTSF
Sbjct  792  NAGFQNSDDAQTSF   805
```

#### angiotensin-converting enzyme 2 [Peromyscus leucopus]

Sequence ID: XP_028743609.1 Length: 805 Number of Matches: 1

**Human vs Mouse**

Range 1: 12 to 805 GenPept Graphics ▼ Next Matc

| Score | Expect | Method | Identities | Positives | Gaps |
|---|---|---|---|---|---|
| 1373 bits(3554) | 0.0 | Compositional matrix adjust. | 658/794(83%) | 712/794(89%) | 0/794 |

```
Query  12   VAVTAAQSTIEEQAKTFLDKFNHEAEDLFYQSSLASWNYNTNITEENVQNMNNAGDKWSA  71
            VAVT AQS IEEQAK FLDKFN EAEDL YQS+LASWNYNTNITEEN Q MN A  KWSA
Sbjct  12   VAVTTAQSIIEEQAKIFLDKFNQEAEDLSYQSALASWNYNTNITEENAQKMNEASAKWSA  71

Query  72   FLKEQSTLAQMYPLQEIQNLTVKLQLQALQQNGSSVLSEDKSKRLNTILNTMSTIYSTGK  131
            F +EQS LA+  Y LQEI NL +K QLQALQQ+GSS LS DK+K+LNTILN MSTIYSTGK
Sbjct  72   FYEEQSKLAKNYSLQEIPNLIIKRQLQALQQSGSSALSADKTKQLNTILNAMSTIYSTGK  131

Query  132  VCNPDNPQECLLLEPGLNEIMANSLDYNERLWAWESWRSEVGKQLRPLYEEYVVLKNEMA  191
            VC P NPQECLLLEPGL+ IMA S DYNERLWAWE WR+EVGKQLRPLYEEYVVLKNEMA
Sbjct  132  VCKPKNPQECLLLEPGLDVIMATSTDYNERLWAWEGWRAEVGKQLRPLYEEYVVLKNEMA  191

Query  192  RANHYEDYGDYWRGDYEVNGVDGYDYSRGQLIEDVEHTFEEIKPLYEHLHAYVRAKLMNA  251
            RAN+Y DYGDYWRGDYE  G +GY+Y+R QLIEDVE  F+EIKPLYEHLHAYVR KLM+
Sbjct  192  RANNYRDYGDYWRGDYEAEGAEGYNYNRNQLIEDVERIFQEIKPLYEHLHAYVRTKLMDT  251

Query  252  YPSYISPIGCLPAHLLGDMWGRFWTNLYSLTVPFGQKPNIDVTDAMVDQAWDAQRIFKEA  311
            YPSYI+P GCLPAHLLGDMWGRFWTNLY LTVPFGQKPNIDVTDAM+ Q WDA+RIFKEA
Sbjct  252  YPSYINPTGCLPAHLLGDMWGRFWTNLYPLTVPFGQKPNIDVTDAMLKQGWDAERIFKEA  311

Query  312  EKFFVSVGLPNMTQGFWENSMLTDPGNVQKAVCHPTAWDLGKGDFRILMCTKVTMDDFLT  371
            EKFFVS+GLP MTQGFWENSML DPG+ +K VCHPTAWDLGKGDFRI MCT VTMD+FLT
Sbjct  312  EKFFVSIGLPPMTQGFWENSMLVDPGDDRKVVCHPTAWDLGKGDFRIKMCTTVTMDNFLT  371

Query  372  AHHEMGHIQYDMAYAAQPFLLRNGANEGFHEAVGEIMSLSAATPKHLKSIGLLSPDFQED  431
            AHHEMGHIQYDMAYA QPFLLRNGANEGFHEAVGEIMSLSAATP+HLKSIGLL  DF+ED
Sbjct  372  AHHEMGHIQYDMAYATQPFLLRNGANEGFHEAVGEIMSLSAATPEHLKSIGLLPSDFRED  431

Query  432  NETEINFLLKQALTIVGTLPFTYMLEKWRWMVFKGEIPKDQWMKKWWEMKREIVGVVEPV  491
            +ETEINFLLKQALTIVGTLPFTYMLEKWRWMVF GEIPK+QWM+KWWEMKREIVGVVEP+
Sbjct  432  SETEINFLLKQALTIVGTLPFTYMLEKWRWMVFTGEIPKEQWMQKWWEMKREIVGVVEPL  491

Query  492  PHDETYCDPASLFHVSNDYSFIRYYTRTLYQFQFQEALCQAAKHEGPLHKCDISNSTEAG  551
            PHDETYCDPA+LFHVSNDYSFIRYYTRT+YQFQFQEALCQAAKH+GPLHKCDISNSTEAG
Sbjct  492  PHDETYCDPAALFHVSNDYSFIRYYTRTIYQFQFQEALCQAAKHDGPLHKCDISNSTEAG  551

Query  552  QKLFNMLRLGKSEPWTLALENVVGAKNMNVRPLLNYFEPLFTWLKDQNKNSFVGWSTDWS  611
            QKL NMLRLG SEPWTLALENVVGA+NM+VRPLLNYFEPLF WLK+QNKNS VGW+TDWS
Sbjct  552  QKLLNMLRLGNSEPWTLALENVVGARNMDVRPLLNYFEPLFAWLKEQNKNSLVGWNTDWS  611

Query  612  PYADQSIKVRISLKSALGDKAYEWNDNEMYLFRSSVAYAMRQYFLKVKNQMILFGEEDVR  671
            PYADQSIKVRISLKSALG  AYEWNDNEMYLFRSSVAYAMR YF   K Q + FG ED+
Sbjct  612  PYADQSIKVRISLKSALGKNAYEWNDNEMYLFRSSVAYAMRVYFAMNKTQNVPFGVEDIH  671

Query  672  VANLKPRISFNFFVTAPKNVSDIIPRTEVEKAIRMSRSRINDAFRLNDNSLEFLGIQPTL  731
            V++LKPR+SFNFFVT+P+NVSDIIPR +VE AIR SR RIND F L+DNSLEFLGI PTL
Sbjct  672  VSDLKPRVSFNFFVTSPQNVSDIIPRKDVEDAIRFSRGRINDVFGLDDNSLEFLGIYPTL  731

Query  732  GPPNQPPVSIWLIVFGVVMGVIVVGIVILIFTGIRDRKKKNKARSGENPYASIDISKGEN  791
             PP QPPV+IWLI+FG+VMG++VVGIVILIFTGI+ RKKKN+ +  ENPY S+DI KGE+
Sbjct  732  APPYQPPVTIWLIIFGIVMGIVVVGIVILIFTGIKGRKKKNETKREENPYDSMDIGKGES  791

Query  792  NPGFQNTDDVQTSF  805
            N GFQN DD QTSF
Sbjct  792  NAGFQNNDDAQTSF  805
```

#### angiotensin-converting enzyme 2 [Mus caroli]

Sequence ID: XP_021009138.1 Length: 805 Number of Matches: 1

**Human vs Mouse**

Range 1: 12 to 805 GenPept Graphics ▼ Next Match ▲ Pre

| Score | Expect | Method | Identities | Positives | Gaps |
|---|---|---|---|---|---|
| 1369 bits(3544) | 0.0 | Compositional matrix adjust. | 651/794(82%) | 708/794(89%) | 0/794(0%) |

```
Query  12   VAVTAAQSTIEEQAKTFLDKFNHEAEDLFYQSSLASWNYNTNITEENVQNMNNAGDKWSA  71
            VAVT AQS  EE AKTFL+KFN EAEDL YQSSLASWNYNTNITEEN Q MN A  KWSA
Sbjct  12   VAVTTAQSLTEENAKTFLNKFNQEAEDLSYQSSLASWNYNTNITEENAQKMNEAAAKWSA  71

Query  72   FLKEQSTLAQMYPLQEIQNLTVKLQLQALQQNGSSVLSEDKSKRLNTILNTMSTIYSTGK  131
            F +EQS  AQ + LQEIQ   +K QLQALQQ+GSS LS DK+K+LNTILNTMSTIYSTGK
Sbjct  72   FYEEQSKTAQSFSLQEIQTPIIKRQLQALQQSGSSALSADKNKQLNTILNTMSTIYSTGK  131

Query  132  VCNPDNPQECLLLEPGLNEIMANSLDYNERLWAWESWRSEVGKQLRPLYEEYVVLKNEMA  191
            VCNP NPQECLLLEPGL+EIMA S DY+ RLWAWE WR+EVGKQLRPLYEEYVVLKNEMA
Sbjct  132  VCNPKNPQECLLLEPGLDEIMATSTDYHSRLWAWEGWRAEVGKQLRPLYEEYVVLKNEMA  191

Query  192  RANHYEDYGDYWRGDYEVNGVDGYDYSRGQLIEDVEHTFEEIKPLYEHLHAYVRAKLMNA  251
            RAN+Y DYGDYWRGDYE  G DGY Y+R QLIEDVE TF EIKPLYEHLHAYVR KLM+
Sbjct  192  RANNYNDYGDYWRGDYEAEGADGYSYNRNQLIEDVERTFAEIKPLYEHLHAYVRRKLMDT  251

Query  252  YPSYISPIGCLPAHLLGDMWGRFWTNLYSLTVPFGQKPNIDVTDAMVDQAWDAQRIFKEA  311
            YPSYISP GCLPAHLLGDMWGRFWTNLY LTVPF QKPNIDVTDAM++Q WDA+RIFKEA
Sbjct  252  YPSYISPTGCLPAHLLGDMWGRFWTNLYPLTVPFAQKPNIDVTDAMMNQGWDAERIFKEA  311

Query  312  EKFFVSVGLPNMTQGFWENSMLTDPGNVQKAVCHPTAWDLGKGDFRILMCTKVTMDDFLT  371
            EKFFVSVGLP+MTQGFW NSMLT+P + +K VCHPTAWDLG GDFRI MCTKVTMD+FLT
Sbjct  312  EKFFVSVGLPHMTQGFWANSMLTEPADGRKVVCHPTAWDLGHGDFRIKMCTKVTMDNFLT  371

Query  372  AHHEMGHIQYDMAYAAQPFLLRNGANEGFHEAVGEIMSLSAATPKHLKSIGLLSPDFQED  431
            AHHEMGHIQYDMAYA QPFLLRNGANEGFHEAVGEIMSLSAATPKHLKSIGLL  +FQED
Sbjct  372  AHHEMGHIQYDMAYATQPFLLRNGANEGFHEAVGEIMSLSAATPKHLKSIGLLPSNFQED  431

Query  432  NETEINFLLKQALTIVGTLPFTYMLEKWRWMVFKGEIPKDQWMKKWWEMKREIVGVVEPV  491
            +ETEINFLLKQALTIVGTLPFTYMLEKWRWMVF+GEIPK+QWMKKWWEMKREIVGVVEP+
Sbjct  432  SETEINFLLKQALTIVGTLPFTYMLEKWRWMVFRGEIPKEQWMKKWWEMKREIVGVVEPL  491

Query  492  PHDETYCDPASLFHVSNDYSFIRYYTRTLYQFQFQEALCQAAKHEGPLHKCDISNSTEAG  551
            PHDETYCDPASLFHVSNDYSFIRYYTRT+YQFQFQEALCQAAK+ GPLHKCDISNSTEAG
Sbjct  492  PHDETYCDPASLFHVSNDYSFIRYYTRTIYQFQFQEALCQAAKYNGPLHKCDISNSTEAG  551

Query  552  QKLFNMLRLGKSEPWTLALENVVGAKNMNVRPLLNYFEPLFTWLKDQNKNSFVGWSTDWS  611
            QKL  ML LG SEPWT ALENVVGA+NM+V+PLLNYF+PLF WLK+QN+NSFVGW+TDWS
Sbjct  552  QKLLKMLSLGNSEPWTKALENVVGARNMDVKPLLNYFQPLFDWLKEQNRNSFVGWNTDWS  611

Query  612  PYADQSIKVRISLKSALGDKAYEWNDNEMYLFRSSVAYAMRQYFLKVKNQMILFGEEDVR  671
            PYADQSIKVRISLKSALG  AY W +NEM+LFRSSVAYAMR+YF  VK Q + F EEDVR
Sbjct  612  PYADQSIKVRISLKSALGVNAYNWTNNEMFLFRSSVAYAMREYFSTVKKQTVPFLEEDVR  671

Query  672  VANLKPRISFNFFVTAPKNVSDIIPRTEVEKAIRMSRSRINDAFRLNDNSLEFLGIQPTL  731
            V +LKPR+SF FFVT+P+NVSD+IPR+EVE AIR SR RIND FRLNDNSLEFLGI PTL
Sbjct  672  VRDLKPRVSFYFFVTSPQNVSDVIPRSEVEDAIRRSRGRINDIFRLNDNSLEFLGIHPTL  731

Query  732  GPPNQPPVSIWLIVFGVVMGVIVVGIVILIFTGIRDRKKKNKARSGENPYASIDISKGEN  791
             PP QPP++IWLI+FGVVM ++VVGI+ILI TGI+ RKKKN+ +  ENPY S+DI KGE+
Sbjct  732  EPPYQPPITIWLIIFGVVMALVVVGIIILIVTGIKGRKKKNETKREENPYDSMDIGKGES  791

Query  792  NPGFQNTDDVQTSF  805
            N GFQN+DD QTSF
Sbjct  792  NAGFQNSDDAQTSF  805
```

#### angiotensin-converting enzyme 2 [Mus pahari]

Sequence ID: XP_021043935.1 Length: 805 Number of Matches: 1

**Human vs Mouse**

Range 1: 12 to 805 GenPept Graphics ▼ Next Match

| Score | Expect | Method | Identities | Positives | Gaps |
|---|---|---|---|---|---|
| 1377 bits(3565) | 0.0 | Compositional matrix adjust. | 658/794(83%) | 718/794(90%) | 0/794( |

```
Query  12   VAVTAAQSTIEEQAKTFLDKFNHEAEDLFYQSSLASWNYNTNITEENVQNMNNAGDKWSA  71
            VAVT AQS  EE AKTFL+KFN EAEDL YQSSLASWNYNTNITEEN Q MN A  KWSA
Sbjct  12   VAVTTAQSLTEENAKTFLNKFNQEAEDLSYQSSLASWNYNTNITEENAQKMNEAAAKWSA  71

Query  72   FLKEQSTLAQMYPLQEIQNLTVKLQLQALQQNGSSVLSEDKSKRLNTILNTMSTIYSTGK  131
            F +EQS  AQ + LQEIQN  +K QLQALQQ+GSS LS DK+K+LNTILNTMSTIYSTGK
Sbjct  72   FYEEQSKTAQNFSLQEIQNPVIKRQLQALQQSGSSALSADKNKQLNTILNTMSTIYSTGK  131

Query  132  VCNPDNPQECLLLEPGLNEIMANSLDYNERLWAWESWRSEVGKQLRPLYEEYVVLKNEMA  191
            VCNP NPQECL+LEPGL+EIMA S DYN RLWAWE WR+EVGKQLRPLYEEYVVLKNEMA
Sbjct  132  VCNPKNPQECLVLEPGLDEIMATSTDYNTRLWAWEGWRAEVGKQLRPLYEEYVVLKNEMA  191

Query  192  RANHYEDYGDYWRGDYEVNGVDGYDYSRGQLIEDVEHTFEEIKPLYEHLHAYVRAKLMNA  251
            RAN+Y+DYGDYWRGDYE  G DGY+Y+R QLIEDVE TF EIKPLYEHLHAYVR KLM+
Sbjct  192  RANNYKDYGDYWRGDYEAEGTDGYNYNRNQLIEDVERTFAEIKPLYEHLHAYVRRKLMDT  251

Query  252  YPSYISPIGCLPAHLLGDMWGRFWTNLYSLTVPFGQKPNIDVTDAMVDQAWDAQRIFKEA  311
            YPSYISP GCLPAHLLGDMWGRFWTNLY LTVPF QKPNIDVTDAM +Q+WDA+RIFKEA
Sbjct  252  YPSYISPTGCLPAHLLGDMWGRFWTNLYPLTVPFAQKPNIDVTDAMTNQSWDAERIFKEA  311

Query  312  EKFFVSVGLPNMTQGFWENSMLTDPGNVQKAVCHPTAWDLGKGDFRILMCTKVTMDDFLT  371
            EKFFVSVGLP+MTQGFW NSMLT+P + +K VCHPTAWDLG GDFRI MCTKVTMD+FLT
Sbjct  312  EKFFVSVGLPHMTQGFWANSMLTEPADGRKVVCHPTAWDLGHGDFRIKMCTKVTMDNFLT  371

Query  372  AHHEMGHIQYDMAYAAQPFLLRNGANEGFHEAVGEIMSLSAATPKHLKSIGLLSPDFQED  431
            AHHEMGHIQYDMAYA QPFLLRNGANEGFHEAVGEIMSLSAATPKHLKSIGLL  +FQED
Sbjct  372  AHHEMGHIQYDMAYATQPFLLRNGANEGFHEAVGEIMSLSAATPKHLKSIGLLPSNFQED  431

Query  432  NETEINFLLKQALTIVGTLPFTYMLEKWRWMVFKGEIPKDQWMKKWWEMKREIVGVVEPV  491
            +ETEINFLLKQALTIVGTLPFTYMLEKWRWMVF+GEIPK+QWMKKWWEMKREIVGVVEP+
Sbjct  432  SETEINFLLKQALTIVGTLPFTYMLEKWRWMVFQGEIPKEQWMKKWWEMKREIVGVVEPL  491

Query  492  PHDETYCDPASLFHVSNDYSFIRYYTRTLYQFQFQEALCQAAKHEGPLHKCDISNSTEAG  551
            PHDETYCDPASLFHVSNDYSFIRYYTRT+YQFQFQEALCQAAK++GPLHKCDISNSTEAG
Sbjct  492  PHDETYCDPASLFHVSNDYSFIRYYTRTIYQFQFQEALCQAAKYDGPLHKCDISNSTEAG  551

Query  552  QKLFNMLRLGKSEPWTLALENVVGAKNMNVRPLLNYFEPLFTWLKDQNKNSFVGWSTDWS  611
            QKL  ML LG SEPWTLALE VVGA+NM+V+PLLNYF+PLF WLK+QN+NSFVGW+TDWS
Sbjct  552  QKLLKMLSLGNSEPWTLALEKVVGARNMDVKPLLNYFQPLFDWLKEQNRNSFVGWNTDWS  611

Query  612  PYADQSIKVRISLKSALGDKAYEWNDNEMYLFRSSVAYAMRQYFLKVKNQMILFGEEDVR  671
            PYADQSIKVRISLKSALG+ AYEW DNEM+LFRSSVAYAMR+YF +VKNQ + F EEDVR
Sbjct  612  PYADQSIKVRISLKSALGNNAYEWTDNEMFLFRSSVAYAMREYFSRVKNQTVPFREEDVR  671

Query  672  VANLKPRISFNFFVTAPKNVSDIIPRTEVEKAIRMSRSRINDAFRLNDNSLEFLGIQPTL  731
            V++LKPR+SFNFFVT+P+NVSD+IPR+EVE AIRMSR RIND F LNDNSLEFLGI PTL
Sbjct  672  VSDLKPRVSFNFFVTSPENVSDVIPRSEVEDAIRMSRGRINDIFGLNDNSLEFLGIHPTL  731

Query  732  GPPNQPPVSIWLIVFGVVMGVIVVGIVILIFTGIRDRKKKNKARSGENPYASIDISKGEN  791
             PP QPPV+IWLI+FGVVMG++VVGI+ILI TGI+ RKKKN+ +  ENPY S+DI KGE+
Sbjct  732  EPPYQPPVTIWLIIFGVVMGLVVVGIIILIVTGIKGRKKKNETKREENPYDSMDIGKGES  791

Query  792  NPGFQNTDDVQTSF  805
            N GFQN DD QTSF
Sbjct  792  NAGFQNNDDAQTSF  805
```

#### angiotensin-converting enzyme 2 [Mastomys coucha]

Sequence ID: XP_031226742.1 Length: 806 Number of Matches: 1

**Human vs Mouse**

Range 1: 12 to 806 GenPept Graphics ▼ Next Match ▲ Pre

| Score | Expect | Method | Identities | Positives | Gaps |
|---|---|---|---|---|---|
| 1352 bits(3499) | 0.0 | Compositional matrix adjust. | 647/795(81%) | 717/795(90%) | 0/795(0%) |

```
Query  11   LVAVTAAQSTIEEQAKTFLDKFNHEAEDLFYQSSLASWNYNTNITEENVQNMNNAGDKWS  70
            + AVT AQS  EE AKTFL+KFN EAEDL YQSSLASWNYNTNITEEN Q MN A  KWS
Sbjct  12   VAAVTTAQSLTEENAKTFLNKFNQEAEDLSYQSSLASWNYNTNITEENAQKMNEAAAKWS  71

Query  71   AFLKEQSTLAQMYPLQEIQNLTVKLQLQALQQNGSSVLSEDKSKRLNTILNTMSTIYSTG  130
            AF +EQS +AQ + LQEI+    +KLQLQALQQ+GSS LS DK K+LNTILNTMSTIYSTG
Sbjct  72   AFYEEQSKIAQNFSLQEIKTPIIKLQLQALQQSGSSALSADKVKQLNTILNTMSTIYSTG  131

Query  131  KVCNPDNPQECLLLEPGLNEIMANSLDYNERLWAWESWRSEVGKQLRPLYEEYVVLKNEM  190
            KVCNP NPQ+CLLLEPGL+EIMA S DYN RLWAWE WR++VGKQLRPLYEEYVVLKNEM
Sbjct  132  KVCNPKNPQQCLLLEPGLDEIMATSTDYNSRLWAWEGWRADVGKQLRPLYEEYVVLKNEM  191

Query  191  ARANHYEDYGDYWRGDYEVNGVDGYDYSRGQLIEDVEHTFEEIKPLYEHLHAYVRAKLMN  250
            ARAN+Y+DYGDYWRGDYE  G +GY+Y+R QLIEDV+ TF EIKPLYEHLHAYVR KLM+
Sbjct  192  ARANNYKDYGDYWRGDYEAEGAEGYNYNRNQLIEDVDRTFAEIKPLYEHLHAYVRTKLMD  251

Query  251  AYPSYISPIGCLPAHLLGDMWGRFWTNLYSLTVPFGQKPNIDVTDAMVDQAWDAQRIFKE  310
             YPSYISP GCLPAHLLGDMWGRFWTNLY LTVPF QKPNIDVTDAM++Q+WDA+RIFKE
Sbjct  252  TYPSYISPTGCLPAHLLGDMWGRFWTNLYPLTVPFAQKPNIDVTDAMMNQSWDAERIFKE  311

Query  311  AEKFFVSVGLPNMTQGFWENSMLTDPGNVQKAVCHPTAWDLGKGDFRILMCTKVTMDDFL  370
            AEKFFVSVGLP+MTQGFW NSMLT+P + +K VCHPTAWDLG GDFRI MCTKVTMD++L
Sbjct  312  AEKFFVSVGLPHMTQGFWANSMLTEPEDDRKVVCHPTAWDLGHGDFRIKMCTKVTMDNYL  371

Query  371  TAHHEMGHIQYDMAYAAQPFLLRNGANEGFHEAVGEIMSLSAATPKHLKSIGLLSPDFQE  430
            TAHHEMGHIQYDMAYA QPFLLRNGANEGFHEAVGEIMSLSAATPKHLKSIGLL  +FQE
Sbjct  372  TAHHEMGHIQYDMAYATQPFLLRNGANEGFHEAVGEIMSLSAATPKHLKSIGLLPSNFQE  431

Query  431  DNETEINFLLKQALTIVGTLPFTYMLEKWRWMVFKGEIPKDQWMKKWWEMKREIVGVVEP  490
            D+ETEINFLLKQALTIVGTLPFTYMLEKWRWMVF+GEIPK+QWMKKWWEMKREIVGVVEP
Sbjct  432  DSETEINFLLKQALTIVGTLPFTYMLEKWRWMVFQGEIPKEQWMKKWWEMKREIVGVVEP  491

Query  491  VPHDETYCDPASLFHVSNDYSFIRYYTRTLYQFQFQEALCQAAKHEGPLHKCDISNSTEA  550
            +PHDETYCDPASLFHVSNDYSFIRYYTRT+YQFQFQEALC+AAKH+GPLHKCDISNSTEA
Sbjct  492  LPHDETYCDPASLFHVSNDYSFIRYYTRTIYQFQFQEALCKAAKHDGPLHKCDISNSTEA  551

Query  551  GQKLFNMLRLGKSEPWTLALENVVGAKNMNVRPLLNYFEPLFTWLKDQNKNSFVGWSTDW  610
            GQKL NMLRLG SEPWTLALENVVGA+NM+V+PLLNYFEPLF WLK+QN+NSFVGW+ DW
Sbjct  552  GQKLLNMLRLGNSEPWTLALENVVGARNMDVKPLLNYFEPLFVWLKEQNRNSFVGWNADW  611

Query  611  SPYADQSIKVRISLKSALGDKAYEWNDNEMYLFRSSVAYAMRQYFLKVKNQMILFGEEDV  670
            SPYADQSIKVRISLKSALG  AYEW ++EM+LFRSSVAYAMR+YF ++K Q + F EEDV
Sbjct  612  SPYADQSIKVRISLKSALGSNAYEWTNDEMFLFRSSVAYAMRKYFSEIKKQTVPFREEDV  671

Query  671  RVANLKPRISFNFFVTAPKNVSDIIPRTEVEKAIRMSRSRINDAFRLNDNSLEFLGIQPT  730
             V +L+PR+SFNFFVT+P+NVSD+IPR+EVE AI MSR RIND F L+DNSLEFLGI PT
Sbjct  672  WVRDLRPRVSFNFFVTSPQNVSDVIPRSEVENAISMSRGRINDVFGLDDNSLEFLGINPT  731

Query  731  LGPPNQPPVSIWLIVFGVVMGVIVVGIVILIFTGIRDRKKKNKARSGENPYASIDISKGE  790
            L PP QPPV+IWLI+FGVVMG++VVGI+ILI TGI+ RKKKN+ +  ENPY S+DI KGE
Sbjct  732  LEPPYQPPVTIWLIIFGVVMGLVVVGIIILIITGIKGRKKKNETKREENPYDSMDIGKGE  791

Query  791  NNPGFQNTDDVQTSF  805
            +N GFQN+DD QTSF
Sbjct  792  SNAGFQNSDDAQTSF  806
```

**PREDICTED: angiotensin-converting enzyme 2 [Peromyscus maniculatus bairdi**

Sequence ID: XP_006973269.1 Length: 805 Number of Matches: 1

#### Human vs Mouse

Range 1: 12 to 805 GenPept Graphics ▼ Next Match

| Score | Expect | Method | Identities | Positives | Gaps |
|---|---|---|---|---|---|
| 1384 bits(3583) | 0.0 | Compositional matrix adjust. | 662/794(83%) | 714/794(89%) | 0/794( |

```
Query  12   VAVTAAQSTIEEQAKTFLDKFNHEAEDLFYQSSLASWNYNTNITEENVQNMNNAGDKWSA  71
            VAVT AQS IEEQAK FLDKFN EAEDL YQS+LASWNYNTNITEEN Q MN A  KWSA
Sbjct  12   VAVTTAQSIIEEQAKIFLDKFNQEAEDLSYQSALASWNYNTNITEENAQKMNEASAKWSA  71

Query  72   FLKEQSTLAQMYPLQEIQNLTVKLQLQALQQNGSSVLSEDKSKRLNTILNTMSTIYSTGK  131
            F +EQS LA+ Y LQEIQNL +K QLQALQQ+GSS LS DK+K+LNTILN MSTIYSTGK
Sbjct  72   FYEEQSKLAKNYSLQEIQNLIIKRQLQALQQSGSSALSADKNKQLNTILNAMSTIYSTGK  131

Query  132  VCNPDNPQECLLLEPGLNEIMANSLDYNERLWAWESWRSEVGKQLRPLYEEYVVLKNEMA  191
            VC P NPQECLLLEPGL+ IMA S DYNERLWAWE WR+EVGKQLRPLYEEYVVLKNEMA
Sbjct  132  VCKPKNPQECLLLEPGLDVIMATSTDYNERLWAWEGWRAEVGKQLRPLYEEYVVLKNEMA  191

Query  192  RANHYEDYGDYWRGDYEVNGVDGYDYSRGQLIEDVEHTFEEIKPLYEHLHAYVRAKLMNA  251
            RAN+Y DYGDYWRGDYE  G +GY+Y+R QLIEDVE TF+EIKPLYEHLHAYVR KLM+
Sbjct  192  RANNYRDYGDYWRGDYEAEGAEGYNYNRNQLIEDVERTFQEIKPLYEHLHAYVRTKLMDT  251

Query  252  YPSYISPIGCLPAHLLGDMWGRFWTNLYSLTVPFGQKPNIDVTDAMVDQAWDAQRIFKEA  311
            YPSYI+P GCLPAHLLGDMWGRFWTNLY LTVPFGQKPNIDVTDAM+ Q WDA+RIFKEA
Sbjct  252  YPSYINPTGCLPAHLLGDMWGRFWTNLYPLTVPFGQKPNIDVTDAMLKQGWDAERIFKEA  311

Query  312  EKFFVSVGLPNMTQGFWENSMLTDPGNVQKAVCHPTAWDLGKGDFRILMCTKVTMDDFLT  371
            EKFFVS+GLP+MTQGFW NSML DPG+ +K VCHPTAWDLGKGDFRI MCT VTMD+FLT
Sbjct  312  EKFFVSIGLPHMTQGFWRNSMLVDPGDDRKVVCHPTAWDLGKGDFRIKMCTTVTMDNFLT  371

Query  372  AHHEMGHIQYDMAYAAQPFLLRNGANEGFHEAVGEIMSLSAATPKHLKSIGLLSPDFQED  431
            AHHEMGHIQYDMAYA QPFLLRNGANEGFHEAVGEIMSLSAATP+HLKSIGLL  DF ED
Sbjct  372  AHHEMGHIQYDMAYATQPFLLRNGANEGFHEAVGEIMSLSAATPEHLKSIGLLPSDFHED  431

Query  432  NETEINFLLKQALTIVGTLPFTYMLEKWRWMVFKGEIPKDQWMKKWWEMKREIVGVVEPV  491
            +ETEINFLLKQALTIVGTLPFTYMLEKWRWMVF GEIPK+QWM+KWWEMKREIVGVVEPV
Sbjct  432  SETEINFLLKQALTIVGTLPFTYMLEKWRWMVFTGEIPKEQWMQKWWEMKREIVGVVEPV  491

Query  492  PHDETYCDPASLFHVSNDYSFIRYYTRTLYQFQFQEALCQAAKHEGPLHKCDISNSTEAG  551
            PHDETYCDPA+LFHVSNDYSFIRYYTRT+YQFQFQEALCQAAKH+GPLHKCDISNSTEAG
Sbjct  492  PHDETYCDPAALFHVSNDYSFIRYYTRTIYQFQFQEALCQAAKHDGPLHKCDISNSTEAG  551

Query  552  QKLFNMLRLGKSEPWTLALENVVGAKNMNVRPLLNYFEPLFTWLKDQNKNSFVGWSTDWS  611
            QKL NMLRLG SEPWTLALENVVGA+NM+VRPLLNYFEPLF WLK+QNKNS VGW+TDWS
Sbjct  552  QKLLNMLRLGNSEPWTLALENVVGARNMDVRPLLNYFEPLFAWLKEQNKNSLVGWNTDWS  611

Query  612  PYADQSIKVRISLKSALGDKAYEWNDNEMYLFRSSVAYAMRQYFLKVKNQMILFGEEDVR  671
            PYADQSIKVRISLKSALG  AYEWNDNEMYLFRSSVAYAMR YF K K Q + FG ED+
Sbjct  612  PYADQSIKVRISLKSALGKNAYEWNDNEMYLFRSSVAYAMRVYFAKNKTQNVPFGVEDIH  671

Query  672  VANLKPRISFNFFVTAPKNVSDIIPRTEVEKAIRMSRSRINDAFRLNDNSLEFLGIQPTL  731
            V++LKPR+SFNFFVT+P+NVSDIIPR +VE AIR SR RIND F L+DNSLEFLGI PTL
Sbjct  672  VSDLKPRVSFNFFVTSPQNVSDIIPRKDVEDAIRFSRGRINDVFGLDDNSLEFLGIYPTL  731

Query  732  GPPNQPPVSIWLIVFGVVMGVIVVGIVILIFTGIRDRKKKNKARSGENPYASIDISKGEN  791
             PP QPPV+IWLI+FGVVMG++VVGIVILIFTGI+ RKKKN+ +  ENPY S+DI KGE+
Sbjct  732  APPYQPPVTIWLIIFGVVMGIVVVGIVILIFTGIKGRKKKNETKREENPYDSMDIGKGES  791

Query  792  NPGFQNTDDVQTSF  805
            N GFQN DD QTSF
Sbjct  792  NAGFQNNDDAQTSF  805
```

**PREDICTED: angiotensin-converting enzyme 2 [Condylura cristata]**

Sequence ID: XP_012585871.1 Length: 800 Number of Matches: 1

#### Human vs Mouse

Range 1: 1 to 800 GenPept Graphics ▼ Next Match ▲ Prev

| Score | Expect | Method | Identities | Positives | Gaps |
|---|---|---|---|---|---|
| 1298 bits(3360) | 0.0 | Compositional matrix adjust. | 621/805(77%) | 699/805(86%) | 5/805(0%) |

```
Query  1    MSSSSWLLLSLVAVTAAQSTIEEQAKTFLDKFNHEAEDLFYQSSLASWNYNTNITEENVQ  60
            MS  SWLLLSLVAV AAQS  E Q K FL+ FN EAE+L Y SSLASW+YNTNIT+ENVQ
Sbjct  1    MSGFSWLLLSLVAVAAAQSDTENQTKKFLETFNREAEELSYNSSLASWDYNTNITDENVQ  60

Query  61   NMNNAGDKWSAFLKEQSTLAQMYPLQEIQNLTVKLQLQALQQNGSSVLSEDKSKRLNTIL  120
             MN AG KWSAF +E+S  A+ + +  IQ+  VKLQL++LQQ GSS L+E+K +RLNTIL
Sbjct  61   KMNEAGAKWSAFYEEKSKDAERFQIDMIQDPIVKLQLKSLQQKGSSALTEEKIQRLNTIL  120

Query  121  NTMSTIYSTGKVCNPDNPQECLLLEPGLNEIMANSLDYNERLWAWESWRSEVGKQLRPLY  180
            N MSTIYSTG+VCNP NPQECLLLEPGL++IMA S DY  +LWAWE WR++VGK+LRPLY
Sbjct  121  NEMSTIYSTGEVCNPHNPQECLLLEPGLDDIMATSNDYEAKLWAWEGWRAKVGKRLRPLY  180

Query  181  EEYVVLKNEMARANHYEDYGDYWRGDYEVNGVDGYDYSRGQLIEDVEHTFEEIKPLYEHL  240
            EEYV LKNEMA+AN+YEDYGDYWRGDYE +      Y+R QLIEDVE TF EIKPLYE L
Sbjct  181  EEYVSLKNEMAKANNYEDYGDYWRGDYETDS-----YTRNQLIEDVERTFAEIKPLYEQL  235

Query  241  HAYVRAKLMNAYPSYISPIGCLPAHLLGDMWGRFWTNLYSLTVPFGQKPNIDVTDAMVDQ  300
            HAYVR+KLM AYP YI+P GCLPAHLLGDMWGRFWTNLY LTVPFG KPNIDVTD MV Q
Sbjct  236  HAYVRSKLMKAYPYYINPTGCLPAHLLGDMWGRFWTNLYRLTVPFGHKPNIDVTDEMVKQ  295

Query  301  AWDAQRIFKEAEKFFVSVGLPNMTQGFWENSMLTDPGNVQKAVCHPTAWDLGKGDFRILM  360
             WDA RIF+EAEKFFVSVGLPNMTQGFWENSMLT+P + +K VCHPTAWDLGKGDFRI M
Sbjct  296  GWDANRIFREAEKFFVSVGLPNMTQGFWENSMLTEPKDGRKVVCHPTAWDLGKGDFRIKM  355

Query  361  CTKVTMDDFLTAHHEMGHIQYDMAYAAQPFLLRNGANEGFHEAVGEIMSLSAATPKHLKS  420
            CTKVTMDDFLTAHHEMGHIQYDMAYA QP+LLR+GANEGFHEAVGEIMSLSAATPKHLKS
Sbjct  356  CTKVTMDDFLTAHHEMGHIQYDMAYAKQPYLLRSGANEGFHEAVGEIMSLSAATPKHLKS  415

Query  421  IGLLSPDFQEDNETEINFLLKQALTIVGTLPFTYMLEKWRWMVFKGEIPKDQWMKKWWEM  480
            +GLL  DF ED ETEINFLLKQALTIVGTLPFTYMLEKWRWMVFK EIPKDQWMKKWWEM
Sbjct  416  LGLLPADFSEDPETEINFLLKQALTIVGTLPFTYMLEKWRWMVFKEEIPKDQWMKKWWEM  475

Query  481  KREIVGVVEPVPHDETYCDPASLFHVSNDYSFIRYYTRTLYQFQFQEALCQAAKHEGPLH  540
            KREIVGV EPVPHDE YCDPA+LFHVSNDYSFIRYYTRT+YQFQFQEALC+ A+H GPLH
Sbjct  476  KREIVGVAEPVPHDENYCDPATLFHVSNDYSFIRYYTRTIYQFQFQEALCRTAQHTGPLH  535

Query  541  KCDISNSTEAGQKLFNMLRLGKSEPWTLALENVVGAKNMNVRPLLNYFEPLFTWLKDQNK  600
            KCDISNS EAG KL  ML LG+SE W LALE VVG K MNV+PLLNYFEPLFTWLK+QN+
Sbjct  536  KCDISNSKEAGAKLLEMLSLGRSESWPLALEKVVGEKTMNVKPLLNYFEPLFTWLKEQNR  595

Query  601  NSFVGWSTDWSPYADQSIKVRISLKSALGDKAYEWNDNEMYLFRSSVAYAMRQYFLKVKN  660
            NSFVGWST     ++Q+IKVRISLKSALGD+AYEWN+NEMYLF+SS+AYAMR+YFLKVKN
Sbjct  596  NSFVGWSTSCPSNSEQNIKVRISLKSALGDEAYEWNENEMYLFQSSIAYAMREYFLKVKN  655

Query  661  QMILFGEEDVRVANLKPRISFNFFVTAPKNVSDIIPRTEVEKAIRMSRSRINDAFRLNDN  720
            + I FGE +V ++N KPRISF+F VT+P+N S + P+++VE+AIR+SR RINDAFRL+DN
Sbjct  656  ETIPFGEANVCISNFKPRISFSFIVTSPENPSQVFPKSQVEEAIRLSRGRINDAFRLDDN  715

Query  721  SLEFLGIQPTLGPPNQPPVSIWLIVFGVVMGVIVVGIVILIFTGIRDRKKKNKARSGENP  780
            SLEF+GI PTL PP +PPV IWLI FG+VM V+V+GIV+LIFTGIR+R+K++K  S ENP
Sbjct  716  SLEFVGIYPTLAPPYKPPVDIWLIAFGIVMAVVVIGIVVLIFTGIRERRKRSKETSQENP  775

Query  781  YASIDISKGENNPGFQNTDDVQTSF  805
            YAS D++ GENNPGFQN+DD QTSF
Sbjct  776  YASEDLTTGENNPGFQNSDDTQTSF  800
```

angiotensin-converting enzyme 2 precursor [Rattus norvegicus]

Sequence ID: NP_001012006.1 Length: 805 Number of Matches: 1

See 4 more title(s)

**Human vs Rat**

Range 1: 12 to 805 GenPept Graphics Next Match

| Score | Expect | Method | Identities | Positives | Gaps |
|---|---|---|---|---|---|
| 1360 bits(3520) | 0.0 | Compositional matrix adjust. | 654/794(82%) | 715/794(90%) | 0/794( |

```
Query  12   VAVTAAQSTIEEQAKTFLDKFNHEAEDLFYQSSLASWNYNTNITEENVQNMNNAGDKWSA  71
            VAV  AQS IEE+A++FL+KFN EAEDL YQSSLASWNYNTNITEEN Q MN A  KWSA
Sbjct  12   VAVATAQSLIEEKAESFLNKFNQEAEDLSYQSSLASWNYNTNITEENAQKMNEAAAKWSA  71

Query  72   FLKEQSTLAQMYPLQEIQNLTVKLQLQALQQNGSSVLSEDKSKRLNTILNTMSTIYSTGK  131
            F +EQS +AQ + LQEIQN T+K QL+ALQQ+GSS LS DK+K+LNTILNTMSTIYSTGK
Sbjct  72   FYEEQSKIAQNFSLQEIQNATIKRQLKALQQSGSSALSPDKNKQLNTILNTMSTIYSTGK  131

Query  132  VCNPDNPQECLLLEPGLNEIMANSLDYNERLWAWESWRSEVGKQLRPLYEEYVVLKNEMA  191
            VCN  NPQEC LLEPGL+EIMA S DYN RLWAWE WR+EVGKQLRPLYEEYVVLKNEMA
Sbjct  132  VCNSMNPQECFLLEPGLDEIMATSTDYNRRLWAWEGWRAEVGKQLRPLYEEYVVLKNEMA  191

Query  192  RANHYEDYGDYWRGDYEVNGVDGYDYSRGQLIEDVEHTFEEIKPLYEHLHAYVRAKLMNA  251
            RAN+YEDYGDYWRGDYE  GV+GY+Y+R QLIEDVE+TF+EIKPLYE LHAYVR KLM  
Sbjct  192  RANNYEDYGDYWRGDYEAEGVEGYNYNRNQLIEDVENTFKEIKPLYEQLHAYVRTKLMEV  251

Query  252  YPSYISPIGCLPAHLLGDMWGRFWTNLYSLTVPFGQKPNIDVTDAMVDQAWDAQRIFKEA  311
            YPSYISP GCLPAHLLGDMWGRFWTNLY LT PF QKPNIDVTDAMV+Q+WDA+RIFKEA
Sbjct  252  YPSYISPTGCLPAHLLGDMWGRFWTNLYPLTTPFLQKPNIDVTDAMVNQSWDAERIFKEA  311

Query  312  EKFFVSVGLPNMTQGFWENSMLTDPGNVQKAVCHPTAWDLGKGDFRILMCTKVTMDDFLT  371
            EKFFVSVGLP MT GFW NSMLT+PG+ +K VCHPTAWDLG GDFRI MCTKVTMD+FLT
Sbjct  312  EKFFVSVGLPQMTPGFWTNSMLTEPGDDRKVVCHPTAWDLGHGDFRIKMCTKVTMDNFLT  371

Query  372  AHHEMGHIQYDMAYAAQPFLLRNGANEGFHEAVGEIMSLSAATPKHLKSIGLLSPDFQED  431
            AHHEMGHIQYDMAYA QPFLLRNGANEGFHEAVGEIMSLSAATPKHLKSIGLL   +FQED
Sbjct  372  AHHEMGHIQYDMAYAKQPFLLRNGANEGFHEAVGEIMSLSAATPKHLKSIGLLPSNFQED  431

Query  432  NETEINFLLKQALTIVGTLPFTYMLEKWRWMVFKGEIPKDQWMKKWWEMKREIVGVVEPV  491
            NETEINFLLKQALTIVGTLPFTYMLEKWRWMVF+ +IP++QW KKWWEMKREIVGVVEP+
Sbjct  432  NETEINFLLKQALTIVGTLPFTYMLEKWRWMVFQDKIPREQWTKKWWEMKREIVGVVEPL  491

Query  492  PHDETYCDPASLFHVSNDYSFIRYYTRTLYQFQFQEALCQAAKHEGPLHKCDISNSTEAG  551
            PHDETYCDPASLFHVSNDYSFIRYYTRT+YQFQFQEALCQAAKH+GPLHKCDISNSTEAG
Sbjct  492  PHDETYCDPASLFHVSNDYSFIRYYTRTIYQFQFQEALCQAAKHDGPLHKCDISNSTEAG  551

Query  552  QKLFNMLRLGKSEPWTLALENVVGAKNMNVRPLLNYFEPLFTWLKDQNKNSFVGWSTDWS  611
            QKL NML LG S PWTLALENVVG++NM+V+PLLNYF+PLF WLK+QN+NS VGWSTDWS
Sbjct  552  QKLLNMLSLGNSGPWTLALENVVGSRNMDVKPLLNYFQPLFVWLKEQNRNSTVGWSTDWS  611

Query  612  PYADQSIKVRISLKSALGDKAYEWNDNEMYLFRSSVAYAMRQYFLKVKNQMILFGEEDVR  671
            PYADQSIKVRISLKSALG  AYEW DNEMYLFRSSVAYAMR+YF + KNQ + FGE DV 
Sbjct  612  PYADQSIKVRISLKSALGKNAYEWTDNEMYLFRSSVAYAMREYFSREKNQTVPFGEADVW  671

Query  672  VANLKPRISFNFFVTAPKNVSDIIPRTEVEKAIRMSRSRINDAFRLNDNSLEFLGIQPTL  731
            V++LKPR+SFNFFVT+PKNVSDIIPR+EVE+AIRMSR RIND F LNDNSLEFLGI PTL
Sbjct  672  VSDLKPRVSFNFFVTSPKNVSDIIPRSEVEEAIRMSRGRINDIFGLNDNSLEFLGIYPTL  731

Query  732  GPPNQPPVSIWLIVFGVVMGVIVVGIVILIFTGIRDRKKKNKARSGENPYASIDISKGEN  791
             PP +PPV+IWLI+FGVVMG +VVGIVILI TGI+ RKKKN+ +  ENPY S+DI KGE+
Sbjct  732  KPPYEPPVTIWLIIFGVVMGTVVVGIVILIVTGIKGRKKKNETKREENPYDSMDIGKGES  791

Query  792  NPGFQNTDDVQTSF  805
            N GFQN+DD QTSF
Sbjct  792  NAGFQNSDDAQTSF  805
```

angiotensin-converting enzyme 2 [Heterocephalus glaber]

Sequence ID: XP_004866157.1 Length: 805 Number of Matches: 1

**Human vs Rat**

Range 1: 1 to 805 GenPept Graphics Next Match P

| Score | Expect | Method | Identities | Positives | Gaps |
|---|---|---|---|---|---|
| 1441 bits(3729) | 0.0 | Compositional matrix adjust. | 681/805(85%) | 733/805(91%) | 0/805(0%) |

```
Query  1    MSSSSWLLLSLVAVTAAQSTIEEQAKTFLDKFNHEAEDLFYQSSLASWNYNTNITEENVQ  60
            MS S WL L+LV VTAAQ T EEQAKTFLDKFN EAEDL YQSSLASWNYNTNITEENVQ
Sbjct  1    MSGSLWLFLTLVPVTAAQLTREEQAKTFLDKFNQEAEDLSYQSSLASWNYNTNITEENVQ  60

Query  61   NMNNAGDKWSAFLKEQSTLAQMYPLQEIQNLTVKLQLQALQQNGSSVLSEDKSKRLNTIL  120
             MN AG  WSAF +EQS LA+ Y LQEIQNLTVK QLQALQ +GSSVLS DK+K+LNTIL
Sbjct  61   KMNEAGALWSAFYEEQSKLAKAYSLQEIQNLTVKRQLQALQHSGSSVLSTDKNKQLNTIL  120

Query  121  NTMSTIYSTGKVCNPDNPQECLLLEPGLNEIMANSLDYNERLWAWESWRSEVGKQLRPLY  180
            N MSTIYSTGKVCNP+NPQECLLLEPGL++IMA S DY+ERLWAWE WRSEVGKQLRPLY
Sbjct  121  NMMSTIYSTGKVCNPNNPQECLLLEPGLDDIMAKSTDYSERLWAWEGWRSEVGKQLRPLY  180

Query  181  EEYVVLKNEMARANHYEDYGDYWRGDYEVNGVDGYDYSRGQLIEDVEHTFEEIKPLYEHL  240
            E+YV LKNEMARAN+YEDYGDYWRGDYE   G DGYDYSR QLI DVE TF EIKPLYE L
Sbjct  181  EQYVALKNEMARANNYEDYGDYWRGDYEAEGADGYDYSRNQLIGDVERTFAEIKPLYEQL  240

Query  241  HAYVRAKLMNAYPSYISPIGCLPAHLLGDMWGRFWTNLYSLTVPFGQKPNIDVTDAMVDQ  300
            HAYVRAKLMNAYPS ISPIGCLPAHLLGDMWGRFWTNLY LTVPFGQKPNIDVTDAMV Q
Sbjct  241  HAYVRAKLMNAYPSRISPIGCLPAHLLGDMWGRFWTNLYPLTVPFGQKPNIDVTDAMVSQ  300

Query  301  AWDAQRIFKEAEKFFVSVGLPNMTQGFWENSMLTDPGNVQKAVCHPTAWDLGKGDFRILM  360
            +WDA++IFKEAEKFFVSV LP+MTQGFW+NSMLT+PG+ +K VCHPTAWDLGK DFRI M
Sbjct  301  SWDAEKIFKEAEKFFVSVSLPHMTQGFWQNSMLTEPGDGRKVVCHPTAWDLGKDDFRIKM  360

Query  361  CTKVTMDDFLTAHHEMGHIQYDMAYAAQPFLLRNGANEGFHEAVGEIMSLSAATPKHLKS  420
            CTKVTMD FLTAHHEMGHIQYDMAYA QPFLLRNGANEGFHEAVGEIMSLSAATPKHLKS
Sbjct  361  CTKVTMDHFLTAHHEMGHIQYDMAYAIQPFLLRNGANEGFHEAVGEIMSLSAATPKHLKS  420

Query  421  IGLLSPDFQEDNETEINFLLKQALTIVGTLPFTYMLEKWRWMVFKGEIPKDQWMKKWWEM  480
            IGLL PDF ED+ETEINFLLKQALTIVGTLPFTYMLEKWRWMVF+GEIPK+QWMKKWWEM
Sbjct  421  IGLLPPDFHEDSETEINFLLKQALTIVGTLPFTYMLEKWRWMVFRGEIPKEQWMKKWWEM  480

Query  481  KREIVGVVEPVPHDETYCDPASLFHVSNDYSFIRYYTRTLYQFQFQEALCQAAKHEGPLH  540
            KREIVGVVEPVPHDETYCDPASLFHVSNDYSFIRYYTRT+YQFQFQEALCQAAKH GPLH
Sbjct  481  KREIVGVVEPVPHDETYCDPASLFHVSNDYSFIRYYTRTIYQFQFQEALCQAAKHVGPLH  540

Query  541  KCDISNSTEAGQKLFNMLRLGKSEPWTLALENVVGAKNMNVRPLLNYFEPLFTWLKDQNK  600
            KCDISNSTEAGQKL NMLRLGKSEPWTLALENVVGAKNM+VRPLLNYFEPLFTWLKDQN+
Sbjct  541  KCDISNSTEAGQKLLNMLRLGKSEPWTLALENVVGAKNMDVRPLLNYFEPLFTWLKDQNR  600

Query  601  NSFVGWSTDWSPYADQSIKVRISLKSALGDKAYEWNDNEMYLFRSSVAYAMRQYFLKVKN  660
            NSFVGWST+WSPY+ +SIKVRISLK ALGD AY+WN NEMYLFRSS+A+AMR+YFLKVKN
Sbjct  601  NSFVGWSTEWSPYSKESIKVRISLKLALGDNAYQWNANEMYLFRSSIAFAMRKYFLKVKN  660

Query  661  QMILFGEEDVRVANLKPRISFNFFVTAPKNVSDIIPRTEVEKAIRMSRSRINDAFRLNDN  720
            +M+ F  EDV V++  PRISF F VTAP N++DIIPR+EVE AI MSRSRIND F L+DN
Sbjct  661  EMVPFRAEDVWVSDETPRISFIFIVTAPNNITDIIPRSEVEDAISMSRSRINDIFGLDDN  720

Query  721  SLEFLGIQPTLGPPNQPPVSIWLIVFGVVMGVIVVGIVILIFTGIRDRKKKNKARSGENP  780
            SLEFLGIQPTLGPP QPPV+IWLIVFGVVMG+++VGIVILI TGIRDR+KKN+ +  ENP
Sbjct  721  SLEFLGIQPTLGPPYQPPVTIWLIVFGVVMGLVLVGIVILIITGIRDRRKKNRTKREENP  780

Query  781  YASIDISKGENNPGFQNTDDVQTSF  805
            Y+S D  KGENN GFQN++D+QTSF
Sbjct  781  YSSADTGKGENNIGFQNSEDIQTSF  805
```

**PREDICTED: angiotensin-converting enzyme 2 isoform X1 [Dipodomys ordii]**

Sequence ID: XP_012887572.1 Length: 805 Number of Matches: 1

**Human vs Rat**

Range 1: 12 to 805 GenPept Graphics Next Match

| Score | Expect | Method | Identities | Positives | Gaps |
|---|---|---|---|---|---|
| 1394 bits(3609) | 0.0 | Compositional matrix adjust. | 652/794(82%) | 714/794(89%) | 0/794(0%) |

```
Query  12   VAVTAAQSTIEEQAKTFLDKFNHEAEDLFYQSSLASWNYNTNITEENVQNMNNAGDKWSA  71
            +AVTA+QS+IEE AKTFLD FN EAEDL YQSSLASWNYNTNITEEN Q MN AG  WSA
Sbjct  12   IAVTASQSSIEELAKTFLDNFNQEAEDLSYQSSLASWNYNTNITEENAQRMNEAGAIWSA  71

Query  72   FLKEQSTLAQMYPLQEIQNLTVKLQLQALQQNGSSVLSEDKSKRLNTILNTMSTIYSTGK  131
            F +EQ+ LA++Y LQEIQN  +K QLQ LQQ+GSS LSEDKSKRLNTILN MSTIYSTG
Sbjct  72   FYEEQAKLAKIYSLQEIQNPILKRQLQFLQQSGSSALSEDKSKRLNTILNKMSTIYSTGT  131

Query  132  VCNPDNPQECLLLEPGLNEIMANSLDYNERLWAWESWRSEVGKQLRPLYEEYVVLKNEMA  191
            VCNP+NPQECLLLEPGL++IMA S DY+ERLW WE WRSEVGKQLRPLYEEYVVLKNEMA
Sbjct  132  VCNPNNPQECLLLEPGLDDIMAKSTDYSERLWVWEGWRSEVGKQLRPLYEEYVVLKNEMA  191

Query  192  RANHYEDYGDYWRGDYEVNGVDGYDYSRGQLIEDVEHTFEEIKPLYEHLHAYVRAKLMNA  251
            RAN+YEDYGDYWRGDYE  G DGY+Y+R QLIEDVE TF EIKPLYEHLHAYVRAKLM+
Sbjct  192  RANNYEDYGDYWRGDYEAEGADGYNYNRNQLIEDVERTFAEIKPLYEHLHAYVRAKLMDI  251

Query  252  YPSYISPIGCLPAHLLGDMWGRFWTNLYSLTVPFGQKPNIDVTDAMVDQAWDAQRIFKEA  311
            YPS+I+P GCLPAHLLGDMWGRFWTNLYSL +PF QKPNID+TDAMV QAWDA RIFKEA
Sbjct  252  YPSHINPTGCLPAHLLGDMWGRFWTNLYSLVIPFEQKPNIDITDAMVQQAWDADRIFKEA  311

Query  312  EKFFVSVGLPNMTQGFWENSMLTDPGNVQKAVCHPTAWDLGKGDFRILMCTKVTMDDFLT  371
            EKFFVSVGLP MTQGFWENSMLT+PG+ +K VCHPTAWDLGKGDFRI MCTKVTMD+FLT
Sbjct  312  EKFFVSVGLPKMTQGFWENSMLTEPGDNRKVVCHPTAWDLGKGDFRIKMCTKVTMDNFLT  371

Query  372  AHHEMGHIQYDMAYAAQPFLLRNGANEGFHEAVGEIMSLSAATPKHLKSIGLLSPDFQED  431
            AHHEMGHIQYDMAYA QPFLLRNGANEGFHEAVGEIMSLSA TPKHLKSIGLL P+F ED
Sbjct  372  AHHEMGHIQYDMAYATQPFLLRNGANEGFHEAVGEIMSLSAVTPKHLKSIGLLPPNFHED  431

Query  432  NETEINFLLKQALTIVGTLPFTYMLEKWRWMVFKGEIPKDQWMKKWWEMKREIVGVVEPV  491
            NETEINFLLKQALTIV TLPFT+MLEKWRWMVF+GEIP++QWMK WWEMKREIVGVVEPV
Sbjct  432  NETEINFLLKQALTIVATLPFTFMLEKWRWMVFRGEIPQEQWMKTWWEMKREIVGVVEPV  491

Query  492  PHDETYCDPASLFHVSNDYSFIRYYTRTLYQFQFQEALCQAAKHEGPLHKCDISNSTEAG  551
            PHDETYCDPASLFHVSND+SFIRYYTRT+YQFQFQEALC+AAK+EGPLHKCDISNS EAG
Sbjct  492  PHDETYCDPASLFHVSNDFSFIRYYTRTIYQFQFQEALCKAAKYEGPLHKCDISNSVEAG  551

Query  552  QKLFNMLRLGKSEPWTLALENVVGAKNMNVRPLLNYFEPLFTWLKDQNKNSFVGWSTDWS  611
             KL NMLRLGKSEPWTLALENVVGAKNM+VRPLLNYFEPLF WL++QNKNSFVGW+T W+
Sbjct  552  HKLLNMLRLGKSEPWTLALENVVGAKNMDVRPLLNYFEPLFIWLQEQNKNSFVGWNTAWN  611

Query  612  PYADQSIKVRISLKSALGDKAYEWNDNEMYLFRSSVAYAMRQYFLKVKNQMILFGEEDVR  671
            PY DQSIKVRISLKSALGDKAYEWNDNEMYLF+SSVAYA+R+YF    +NQ I F EE+V+
Sbjct  612  PYNDQSIKVRISLKSALGDKAYEWNDNEMYLFQSSVAYALRKYFSATQNQTIPFREENVK  671

Query  672  VANLKPRISFNFFVTAPKNVSDIIPRTEVEKAIRMSRSRINDAFRLNDNSLEFLGIQPTL  731
            V NL   RISF F+VT P N SDI+PR EVE AIRMSR RIND F L+DNSLEFLGI PTL
Sbjct  672  VENLTQRISFTFYVTMPNNSSDIVPRDEVEAAIRMSRGRINDIFCLDDNSLEFLGINPTL  731

Query  732  GPPNQPPVSIWLIVFGVVMGVIVVGIVILIFTGIRDRKKKNKARSGENPYASIDISKGEN  791
            GPP QPPV++WLI FGVVMG++V+G+V LI   GIR+R+K+++ R   +NPYAS+DISKGE
Sbjct  732  GPPYQPPVTVWLIAFGVVMGLVVIGLVFLIVVGIRERRKRSEIRRE DNPYASVDISKGEM  791

Query  792  NPGFQNTDDVQTSF   805
            N GFQNT+DVQTSF
Sbjct  792  NAGFQNTEDVQTSF   805
```

**PREDICTED: angiotensin-converting enzyme 2 isoform X2 [Dipodomys ordii]**

Sequence ID: XP_012887573.1 Length: 740 Number of Matches: 2

**Human vs Rat**

Range 1: 12 to 713 GenPept Graphics Next Match

| Score | Expect | Method | Identities | Positives | Gaps |
|---|---|---|---|---|---|
| 1244 bits(3218) | 0.0 | Compositional matrix adjust. | 584/702(83%) | 631/702(89%) | 0/702(0%) |

```
Query  12   VAVTAAQSTIEEQAKTFLDKFNHEAEDLFYQSSLASWNYNTNITEENVQNMNNAGDKWSA  71
            +AVTA+QS+IEE AKTFLD FN EAEDL YQSSLASWNYNTNITEEN Q MN AG  WSA
Sbjct  12   IAVTASQSSIEELAKTFLDNFNQEAEDLSYQSSLASWNYNTNITEENAQRMNEAGAIWSA  71

Query  72   FLKEQSTLAQMYPLQEIQNLTVKLQLQALQQNGSSVLSEDKSKRLNTILNTMSTIYSTGK  131
            F +EQ+ LA++Y LQEIQN  +K QLQ LQQ+GSS LSEDKSKRLNTILN MSTIYSTG
Sbjct  72   FYEEQAKLAKIYSLQEIQNPILKRQLQFLQQSGSSALSEDKSKRLNTILNKMSTIYSTGT  131

Query  132  VCNPDNPQECLLLEPGLNEIMANSLDYNERLWAWESWRSEVGKQLRPLYEEYVVLKNEMA  191
            VCNP+NPQECLLLEPGL++IMA S DY+ERLW WE WRSEVGKQLRPLYEEYVVLKNEMA
Sbjct  132  VCNPNNPQECLLLEPGLDDIMAKSTDYSERLWVWEGWRSEVGKQLRPLYEEYVVLKNEMA  191

Query  192  RANHYEDYGDYWRGDYEVNGVDGYDYSRGQLIEDVEHTFEEIKPLYEHLHAYVRAKLMNA  251
            RAN+YEDYGDYWRGDYE  G DGY+Y+R QLIEDVE TF EIKPLYEHLHAYVRAKLM+
Sbjct  192  RANNYEDYGDYWRGDYEAEGADGYNYNRNQLIEDVERTFAEIKPLYEHLHAYVRAKLMDI  251

Query  252  YPSYISPIGCLPAHLLGDMWGRFWTNLYSLTVPFGQKPNIDVTDAMVDQAWDAQRIFKEA  311
            YPS+I+P GCLPAHLLGDMWGRFWTNLYSL +PF QKPNID+TDAMV QAWDA RIFKEA
Sbjct  252  YPSHINPTGCLPAHLLGDMWGRFWTNLYSLVIPFEQKPNIDITDAMVQQAWDADRIFKEA  311

Query  312  EKFFVSVGLPNMTQGFWENSMLTDPGNVQKAVCHPTAWDLGKGDFRILMCTKVTMDDFLT  371
            EKFFVSVGLP MTQGFWENSMLT+PG+ +K VCHPTAWDLGKGDFRI MCTKVTMD+FLT
Sbjct  312  EKFFVSVGLPKMTQGFWENSMLTEPGDNRKVVCHPTAWDLGKGDFRIKMCTKVTMDNFLT  371

Query  372  AHHEMGHIQYDMAYAAQPFLLRNGANEGFHEAVGEIMSLSAATPKHLKSIGLLSPDFQED  431
            AHHEMGHIQYDMAYA QPFLLRNGANEGFHEAVGEIMSLSA TPKHLKSIGLL P+F ED
Sbjct  372  AHHEMGHIQYDMAYATQPFLLRNGANEGFHEAVGEIMSLSAVTPKHLKSIGLLPPNFHED  431

Query  432  NETEINFLLKQALTIVGTLPFTYMLEKWRWMVFKGEIPKDQWMKKWWEMKREIVGVVEPV  491
            NETEINFLLKQALTIV TLPFT+MLEKWRWMVF+GEIP++QWMK WWEMKREIVGVVEPV
Sbjct  432  NETEINFLLKQALTIVATLPFTFMLEKWRWMVFRGEIPQEQWMKTWWEMKREIVGVVEPV  491

Query  492  PHDETYCDPASLFHVSNDYSFIRYYTRTLYQFQFQEALCQAAKHEGPLHKCDISNSTEAG  551
            PHDETYCDPASLFHVSND+SFIRYYTRT+YQFQFQEALC+AAK+EGPLHKCDISNS EAG
Sbjct  492  PHDETYCDPASLFHVSNDFSFIRYYTRTIYQFQFQEALCKAAKYEGPLHKCDISNSVEAG  551

Query  552  QKLFNMLRLGKSEPWTLALENVVGAKNMNVRPLLNYFEPLFTWLKDQNKNSFVGWSTDWS  611
             KL NMLRLGKSEPWTLALENVVGAKNM+VRPLLNYFEPLF WL++QNKNSFVGW+T W+
Sbjct  552  HKLLNMLRLGKSEPWTLALENVVGAKNMDVRPLLNYFEPLFIWLQEQNKNSFVGWNTAWN  611

Query  612  PYADQSIKVRISLKSALGDKAYEWNDNEMYLFRSSVAYAMRQYFLKVKNQMILFGEEDVR  671
            PY DQSIKVRISLKSALGDKAYEWNDNEMYLF+SSVAYA+R+YF    +NQ I F EE+V+
Sbjct  612  PYNDQSIKVRISLKSALGDKAYEWNDNEMYLFQSSVAYALRKYFSATQNQTIPFREENVK  671

Query  672  VANLKPRISFNFFVTAPKNVSDIIPRTEVEKAIRMSRSRIND   713
            V NL   RISF F+VT P N SDI+PR EVE AIR S   R  D
Sbjct  672  VENLTQRISFTFYVTMPNNSSDIVPRDEVEAAIRRSEIRRED   713
```

**angiotensin-converting enzyme 2 [Rattus rattus]**

Sequence ID: XP_032746145.1 Length: 793 Number of Matches: 1

#### Human vs Rat

Range 1: 1 to 793 GenPept Graphics Next Matc

| Score | Expect | Method | Identities | Positives | Gaps |
| --- | --- | --- | --- | --- | --- |
| 1320 bits(3416) | 0.0 | Compositional matrix adjust. | 640/807(79%) | 709/807(87%) | 16/80 |

```
Query  1    MSSSSWLLLSLVAVTAAQSTIEEQAKTFLDKFNHEAEDLFYQSSLASWNYNTNITEENVQ  60
            MS S WLLLSLVAV  AQS IEE+A++FL+KFN EAEDL YQSSLASWNYNTNITEEN Q
Sbjct  1    MSRSPWLLLSLVAVATAQSLIEEKAESFLNKFNQEAEDLSYQSSLASWNYNTNITEENAQ  60

Query  61   NMNNAGDKWSAFLKEQSTLAQMYPLQEIQNLTVKLQLQALQQNGSSVLSEDKSKRLNTIL  120
             MN A  KWSAF +EQS +AQ  + LQEIQ+ T+K QL+ALQQ+GSS LS DK+K+LNTIL
Sbjct  61   KMNEAAAKWSAFYEEQSKIAQNFSLQEIQDATIKRQLKALQQSGSSALSPDKNKQLNTIL  120

Query  121  NTMSTIYSTGKVCNPDNPQECLLLEPGLNEIMANSLDYNERLWAWESWRSEVGKQLRPLY  180
            NTMSTIYSTGKVCN  NPQEC +LEPGL+EIMA S DYN RLWAWE WR+EVGKQLRPLY
Sbjct  121  NTMSTIYSTGKVCNSMNPQECFVLEPGLDEIMATSTDYNRRLWAWEGWRAEVGKQLRPLY  180

Query  181  EEYVVLKNEMARANHYEDYGDYWRGDYEVNGVDGYDYSRGQLIEDVEHTFEEIKPLYEHL  240
            EEYVVLKNEMARAN+YEDYGDYWRGDYE   GV+GY+Y+R QLIEDVE+TF+EIKPLYE L
Sbjct  181  EEYVVLKNEMARANNYEDYGDYWRGDYEAEGVEGYNYNRNQLIEDVENTFKEIKPLYEQL  240

Query  241  HAYVRAKLMNAYPSYISPIGCLPAHLLGDMWGRFWTNLYSLTVPFGQKPNIDVTDAMVDQ  300
            HAYVR KLM+ YPSYISP GCLPAHLLGDMWGRFWTNLY LT PF QKPNIDVTDAMV+Q
Sbjct  241  HAYVRTKLMDVYPSYISPTGCLPAHLLGDMWGRFWTNLYPLTTPFLQKPNIDVTDAMVNQ  300

Query  301  AWDAQRIFKEAEKFFVSVGLPNMTQGFWENSMLTDPGNVQKAVCHPTAWDLGKGDFRILM  360
            +WDA+RIFKEAEKFFVSVGLP MT GFW NSMLT+PG+ +K VCHPTAWDLG GDFRI M
Sbjct  301  SWDAERIFKEAEKFFVSVGLPQMTPGFWTNSMLTEPGDDRKVVCHPTAWDLGHGDFRIKM  360

Query  361  CTKVTMDDFLTAHHEMGHIQYDMAYAAQPFLLRNGANEGFHEAVGEIMSLSAATPKHLKS  420
            CTKVTMD+FLTAHHEMGHIQYDMAYA QPFLLRNGANEGFHEAVGEIMSLSAATPKHLKS
Sbjct  361  CTKVTMDNFLTAHHEMGHIQYDMAYAKQPFLLRNGANEGFHEAVGEIMSLSAATPKHLKS  420

Query  421  IGLLSPDFQEDNETEINFLLKQALTIVGTLPFTYMLEKWRWMVFKGEIPKDQWMKKWWEM  480
            IGLL   +FQED+ETEINFLLKQAL IVGTLPFTYMLEKWRWMVF+ +IP++QW +KWWEM
Sbjct  421  IGLLPSNFQEDDETEINFLLKQALIIVGTLPFTYMLEKWRWMVFQDKIPREQWTQKWWEM  480

Query  481  KREIVGVVEPVPHDETYCDPASLFHVSNDYSFIRYYTRTLYQFQFQEALCQAAKHEGPLH  540
            KREIVGVVEP+PHDETYCDPASLFHVSNDYSFIRYYTRT+YQFQFQEALCQAAKH+GPLH
Sbjct  481  KREIVGVVEPLPHDETYCDPASLFHVSNDYSFIRYYTRTIYQFQFQEALCQAAKHDGPLH  540

Query  541  KCDISNSTEAGQKLFNMLRLGKSEPWTLALENVVGAKNMNVRPLLNYFEPLFTWLKDQNK  600
            KCDISNSTEAGQKL  NML LG S PWTLALENVVG++NM+V+PLLNYF+PLF WLK+QN
Sbjct  541  KCDISNSTEAGQKLLNMLSLGNSGPWTLALENVVGSRNMDVKPLLNYFQPLFVWLKEQNS  600

Query  601  NSFVGWSTDWSP--YADQSIKVRISLKSALGDKAYEWNDNEMYLFRSSVAYAMRQYFLKV  658
            NS VGW+TDWSP   + D +              YEW DNEMYLFRSSVAYAMR+YF +
Sbjct  601  NSTVGWNTDWSPCDFTDCT--------------HYEWTDNEMYLFRSSVAYAMREYFSRE  646

Query  659  KNQMILFGEEDVRVANLKPRISFNFFVTAPKNVSDIIPRTEVEKAIRMSRSRINDAFRLN  718
            KNQ + FGE DV V++LKPR+SFNFFVT+PKNVSDIIPR+EVE+AIRMSR RIND F LN
Sbjct  647  KNQTVPFGEADVWVSDLKPRVSFNFFVTSPKNVSDIIPRSEVEEAIRMSRGRINDIFGLN  706

Query  719  DNSLEFLGIQPTLGPPNQPPVSIWLIVFGVVMGVIVVGIVILIFTGIRDRKKKNKARSGE  778
            DNSLEFLGI PTL PP +PPV+IWLI+FGVVMG++VVGIVILI TGI+ RKKKN+ +   E
Sbjct  707  DNSLEFLGIYPTLKPPYEPPVTIWLIIFGVVMGMVVVGIVILIVTGIKGRKKKNETKREE  766

Query  779  NPYASIDISKGENNPGFQNTDDVQTSF   805
            NPY S+DI KGE+N GFQN+DD QTSF
Sbjct  767  NPYDSVDIGKGESNAGFQNSDDAQTSF   793
```

**PREDICTED: angiotensin-converting enzyme 2 [Fukomys damarensis]**

Sequence ID: XP_010643477.1 Length: 805 Number of Matches: 1

#### Human vs Rat

Range 1: 1 to 805 GenPept Graphics Next Match

| Score | Expect | Method | Identities | Positives | Gaps |
| --- | --- | --- | --- | --- | --- |
| 1446 bits(3743) | 0.0 | Compositional matrix adjust. | 682/805(85%) | 735/805(91%) | 0/805(0%) |

```
Query  1    MSSSSWLLLSLVAVTAAQSTIEEQAKTFLDKFNHEAEDLFYQSSLASWNYNTNITEENVQ  60
            MS S WLLLSLVAVTAAQ TIEEQAKTFLDKFN EAEDL YQ+SLASWNYNTNITEENVQ
Sbjct  1    MSGSFWLLLSLVAVTAAQLTIEEQAKTFLDKFNQEAEDLSYQNSLASWNYNTNITEENVQ  60

Query  61   NMNNAGDKWSAFLKEQSTLAQMYPLQEIQNLTVKLQLQALQQNGSSVLSEDKSKRLNTIL  120
             MN AG  WS F +EQS LA+ YPLQEIQNLTVK QLQ LQQ+ SS LS DK+K+LNTIL
Sbjct  61   KMNEAGAIWSVFYEEQSKLAKAYPLQEIQNLTVKRQLQVLQQSWSSALSADKNKQLNTIL  120

Query  121  NTMSTIYSTGKVCNPDNPQECLLLEPGLNEIMANSLDYNERLWAWESWRSEVGKQLRPLY  180
            N MSTIYSTGKVCNP+ PQECLLLEPGL++IMA S DY+ERLW WE WRSEVGKQLRPLY
Sbjct  121  NMMSTIYSTGKVCNPNKPQECLLLEPGLDDIMAKSTDYSERLWVWEGWRSEVGKQLRPLY  180

Query  181  EEYVVLKNEMARANHYEDYGDYWRGDYEVNGVDGYDYSRGQLIEDVEHTFEEIKPLYEHL  240
            E+YV LKNEMARAN+YEDYGDYWR DYE   G DGYDYSR QLIEDVE TF EIKPLYE L
Sbjct  181  EQYVALKNEMARANNYEDYGDYWRSDYEAEGADGYDYSRNQLIEDVERTFAEIKPLYEQL  240

Query  241  HAYVRAKLMNAYPSYISPIGCLPAHLLGDMWGRFWTNLYSLTVPFGQKPNIDVTDAMVDQ  300
            HAYVR KLMN YPS +SPIGCLPAHLLGDMWGRFWTNLY LTVPFGQKPNIDVT+AMV+Q
Sbjct  241  HAYVRTKLMNDYPSRLSPIGCLPAHLLGDMWGRFWTNLYPLTVPFGQKPNIDVTEAMVNQ  300

Query  301  AWDAQRIFKEAEKFFVSVGLPNMTQGFWENSMLTDPGNVQKAVCHPTAWDLGKGDFRILM  360
             WDA++IFKEAE+FFVSVGLP+MTQGFW+NSMLT+PG+ +K VCHPTAWDLGK DFRI M
Sbjct  301  FWDAEKIFKEAEQFFVSVGLPHMTQGFWQNSMLTEPGDGRKVVCHPTAWDLGKNDFRIKM  360

Query  361  CTKVTMDDFLTAHHEMGHIQYDMAYAAQPFLLRNGANEGFHEAVGEIMSLSAATPKHLKS  420
            CTKVTMD FLTAHHEMGHIQYDMAY+ QPFLLRNGANEGFHEAVGEIMSLSAATPKHLKS
Sbjct  361  CTKVTMDHFLTAHHEMGHIQYDMAYSTQPFLLRNGANEGFHEAVGEIMSLSAATPKHLKS  420

Query  421  IGLLSPDFQEDNETEINFLLKQALTIVGTLPFTYMLEKWRWMVFKGEIPKDQWMKKWWEM  480
            IGLL P+F EDNETEINFLLKQALTIVGTLPFTYMLEKWRWMVF+GEIPKDQWMKKWWEM
Sbjct  421  IGLLPPNFHEDNETEINFLLKQALTIVGTLPFTYMLEKWRWMVFRGEIPKDQWMKKWWEM  480

Query  481  KREIVGVVEPVPHDETYCDPASLFHVSNDYSFIRYYTRTLYQFQFQEALCQAAKHEGPLH  540
            KREIVGVVEP+PHDETYCDPASLFHVSNDYSFIRYYTRT+YQFQFQEALCQAAKH GPLH
Sbjct  481  KREIVGVVEPMPHDETYCDPASLFHVSNDYSFIRYYTRTIYQFQFQEALCQAAKHVGPLH  540

Query  541  KCDISNSTEAGQKLFNMLRLGKSEPWTLALENVVGAKNMNVRPLLNYFEPLFTWLKDQNK  600
            KCDISNSTEAGQKL NMLRLGKSEPWTLALENVVGAKNM+VRPLLNYFEPLFTWLKDQNK
Sbjct  541  KCDISNSTEAGQKLLNMLRLGKSEPWTLALENVVGAKNMDVRPLLNYFEPLFTWLKDQNK  600

Query  601  NSFVGWSTDWSPYADQSIKVRISLKSALGDKAYEWNDNEMYLFRSSVAYAMRQYFLKVKN  660
            NSFVGWST+WSPY+ +SIKVRISLKSALG  AY+WN NEMYLF+SSVAYAMRQYFLKVKN
Sbjct  601  NSFVGWSTEWSPYSQESIKVRISLKSALGVNAYQWNSNEMYLFQSSVAYAMRQYFLKVKN  660

Query  661  QMILFGEEDVRVANLKPRISFNFFVTAPKNVSDIIPRTEVEKAIRMSRSRINDAFRLNDN  720
            + + F EEDVRV++   PRISF F VTAP N++DIIPR+EVE AIRMSR RIND F L+DN
Sbjct  661  KTVPFREEDVRVSDETPRISFIFIVTAPNNITDIIPRSEVEYAIRMSRGRINDIFSLDDN  720

Query  721  SLEFLGIQPTLGPPNQPPVSIWLIVFGVVMGVIVVGIVILIFTGIRDRKKKNKARSGENP  780
            SLEFLGIQPTLGPP QPPV+IWLIVFGVVMG+++VGIVILIFTGIRDR+KKNK +  ENP
Sbjct  721  SLEFLGIQPTLGPPYQPPVTIWLIVFGVVMGLVLVGIVILIFTGIRDRRKKNKTKGEENP  780

Query  781  YASIDISKGENNPGFQNTDDVQTSF   805
            Y+S+DI KGENN GFQN++D+QTSF
Sbjct  781  YSSVDIGKGENNTGFQNSEDIQTSF   805
```

**angiotensin-converting enzyme 2 [Nannospalax galili]**

Sequence ID: XP_008839098.1  Length: 804  Number of Matches: 1

#### Human vs Rat

Range 1: 12 to 804 GenPept Graphics  ▼ Next Match ▲ Pr

| Score | Expect | Method | Identities | Positives | Gaps |
|---|---|---|---|---|---|
| 1368 bits(3540) | 0.0 | Compositional matrix adjust. | 654/794(82%) | 723/794(91%) | 1/794(0%) |

```
Query  12   VAVTAAQSTIEEQAKTFLDKFNHEAEDLFYQSSLASWNYNTNITEENVQNMNNAGDKWSA  71
            VAVTAAQS IEEQAKTFLDKFN EAEDL YQS+LASW+YNTNITEEN Q MN A  KWSA
Sbjct  12   VAVTAAQS-IEEQAKTFLDKFNQEAEDLSYQSALASWDYNTNITEENAQKMNEAAMKWSA  70

Query  72   FLKEQSTLAQMYPLQEIQNLTVKLQLQALQQNGSSVLSEDKSKRLNTILNTMSTIYSTGK  131
            F +EQS LA+ Y LQEIQ+L +K QLQALQ++G+S LS DK+K+LNTILNTMSTIYSTGK
Sbjct  71   FYEEQSKLAKKYSLQEIQDLVIKRQLQALQESGASALSPDKNKQLNTILNTMSTIYSTGK  130

Query  132  VCNPDNPQECLLLEPGLNEIMANSLDYNERLWAWESWRSEVGKQLRPLYEEYVVLKNEMA  191
            VC  +NPQECLLLEPGL+++MA S DYNERLWAWE WR+EVGKQLRPLYEEYVVLKNEMA
Sbjct  131  VCRSNNPQECLLLEPGLDDLMATSTDYNERLWAWEGWRAEVGKQLRPLYEEYVVLKNEMA  190

Query  192  RANHYEDYGDYWRGDYEVNGVDGYDYSRGQLIEDVEHTFEEIKPLYEHLHAYVRAKLMNA  251
            RAN+YEDYGDYWRGDYE  G DGY+Y+R QLI+DVE TF EIKPLYE LHAYVR KLM+A
Sbjct  191  RANNYEDYGDYWRGDYEAEGADGYEYNRNQLIQDVERTFAEIKPLYEQLHAYVRTKLMDA  250

Query  252  YPSYISPIGCLPAHLLGDMWGRFWTNLYSLTVPFGQKPNIDVTDAMVDQAWDAQRIFKEA  311
            YPS ISP GCLPAHLLGDMWGRFWTN+Y LTVPFGQK NIDVTDAMV Q W A+RIFKEA
Sbjct  251  YPSRISPTGCLPAHLLGDMWGRFWTNIYPLTVPFGQKQNIDVTDAMVQQGWGAERIFKEA  310

Query  312  EKFFVSVGLPNMTQGFWENSMLTDPGNVQKAVCHPTAWDLGKGDFRILMCTKVTMDDFLT  371
            EKFFVSV LP MTQGFWENSMLT+PG  ++ VCHPTAWDLGKGDFRI MCTKVTMD+FLT
Sbjct  311  EKFFVSVDLPQMTQGFWENSMLTEPGGDRQVVCHPTAWDLGKGDFRIKMCTKVTMDNFLT  370

Query  372  AHHEMGHIQYDMAYAAQPFLLRNGANEGFHEAVGEIMSLSAATPKHLKSIGLLSPDFQED  431
            AHHEMGHIQYDMAYA QPFLLRNGANEGFHEAVGEIMSLSAATP+HLKSIGLL  DF++D
Sbjct  371  AHHEMGHIQYDMAYATQPFLLRNGANEGFHEAVGEIMSLSAATPQHLKSIGLLPSDFRDD  430

Query  432  NETEINFLLKQALTIVGTLPFTYMLEKWRWMVFKGEIPKDQWMKKWWEMKREIVGVVEPV  491
            NETEINFLLKQALTIVGTLPFTYMLEKWRWMVFKGEIPK++WMKKWWEMKREIVGVVEP+
Sbjct  431  NETEINFLLKQALTIVGTLPFTYMLEKWRWMVFKGEIPKEEWMKKWWEMKREIVGVVEPL  490

Query  492  PHDETYCDPASLFHVSNDYSFIRYYTRTLYQFQFQEALCQAAKHEGPLHKCDISNSTEAG  551
            PHDETYCDPASLFHVSNDYSFIRYYTRT+YQFQFQEALC+AA+H GPLH+CDISNST+AG
Sbjct  491  PHDETYCDPASLFHVSNDYSFIRYYTRTIYQFQFQEALCRAAEHVGPLHQCDISNSTKAG  550

Query  552  QKLFNMLRLGKSEPWTLALENVVGAKNMNVRPLLNYFEPLFTWLKDQNKNSFVGWSTDWS  611
            QKL NMLRLG SEPWTLALENVVGA+NM+VRPLLNYFEPL  WLK+QN+NSFVGWSTDWS
Sbjct  551  QKLLNMLRLGSSEPWTLALENVVGARNMDVRPLLNYFEPLLVWLKEQNRNSFVGWSTDWS  610

Query  612  PYADQSIKVRISLKSALGDKAYEWNDNEMYLFRSSVAYAMRQYFLKVKNQMILFGEEDVR  671
            PYADQSIKVRISLKSALG+ AY+WNDNEMYLFRSS+AYAMR+YFL+VK Q I FGEE++
Sbjct  611  PYADQSIKVRISLKSALGENAYKWNDNEMYLFRSSIAYAMRKYFLEVKTQTIPFGEENIL  670

Query  672  VANLKPRISFNFFVTAPKNVSDIIPRTEVEKAIRMSRSRINDAFRLNDNSLEFLGIQPTL  731
            V+++KPRISFNF VTAPKNVS+IIPR+EVE++IR+SR RIND F L+DNSLEFLGI PTL
Sbjct  671  VSDVKPRISFNFIVTAPKNVSEIIPRSEVEQSIRLSRGRINDIFYLDDNSLEFLGIHPTL  730

Query  732  GPPNQPPVSIWLIVFGVVMGVIVVGIVILIFTGIRDRKKKNKARSGENPYASIDISKGEN  791
             PP QPPV+IWLI+FGVVMG++VVGI+ILI TGI+ RKKKN+ +  ENPYAS++I KGE+
Sbjct  731  EPPYQPPVTIWLIIFGVVMGIVVVGIIILIVTGIKGRKKKNQTKKEENPYASVEIGKGES  790

Query  792  NPGFQNTDDVQTSF  805
            N GFQN DD QTSF
Sbjct  791  NAGFQNNDDAQTSF  804
```

**angiotensin-converting enzyme 2 [Cricetulus griseus]**

Sequence ID: XP_003503283.1 Length: 805 Number of Matches: 1

#### Human vs Hamster

Range 1: 12 to 805 GenPept Graphics

▼ Next Match

| Score | Expect | Method | Identities | Positives | Gaps |
|---|---|---|---|---|---|
| 1396 bits(3614) | 0.0 | Compositional matrix adjust. | 668/794(84%) | 726/794(91%) | 0/794(0 |

```
Query  12   VAVTAAQSTIEEQAKTFLDKFNHEAEDLFYQSSLASWNYNTNITEENVQNMNNAGDKWSA  71
            VAVT AQS IEEQAKTFLDKFN EAEDL YQS+LASWNYNTNITEEN Q MN A  KWSA
Sbjct  12   VAVTTAQSIIEEQAKTFLDKFNQEAEDLSYQSALASWNYNTNITEENAQKMNEAAAKWSA  71

Query  72   FLKEQSTLAQMYPLQEIQNLTVKLQLQALQQNGSSVLSEDKSKRLNTILNTMSTIYSTGK  131
            F +EQS LA+ Y LQE+QNL +K QLQALQQ+GSS LS DK+K+LNTILNTMSTIYSTGK
Sbjct  72   FYEEQSKLAKNYSLQEVQNLIIKRQLQALQQSGSSALSADKNKQLNTILNTMSTIYSTGK  131

Query  132  VCNPDNPQECLLLEPGLNEIMANSLDYNERLWAWESWRSEVGKQLRPLYEEYVVLKNEMA  191
            VCNP NPQECLLLEPGL++IMA S DYNERLWAWE WR+EVGKQLRPLYEEYVVLKNEMA
Sbjct  132  VCNPKNPQECLLLEPGLDDIMATSTDYNERLWAWEGWRAEVGKQLRPLYEEYVVLKNEMA  191

Query  192  RANHYEDYGDYWRGDYEVNGVDGYDYSRGQLIEDVEHTFEEIKPLYEHLHAYVRAKLMNA  251
            RAN+Y+DYGDYWRGDYE  G DGY+Y+  QLIEDVE TF+EIKPLYE LHAYVR KLM+
Sbjct  192  RANNYKDYGDYWRGDYEAEGADGYNYNGNQLIEDVERTFKEIKPLYEQLHAYVRTKLMDT  251

Query  252  YPSYISPIGCLPAHLLGDMWGRFWTNLYSLTVPFGQKPNIDVTDAMVDQAWDAQRIFKEA  311
            YPSYISP GCLPAHLLGDMWGRFWTNLY LTVPFGQKPNIDVTDAMV+Q WDA+RIFKEA
Sbjct  252  YPSYISPTGCLPAHLLGDMWGRFWTNLYPLTVPFGQKPNIDVTDAMVNQGWDAERIFKEA  311

Query  312  EKFFVSVGLPNMTQGFWENSMLTDPGNVQKAVCHPTAWDLGKGDFRILMCTKVTMDDFLT  371
            EKFFVSVGLP+MTQGFW NSMLTDPG+ +K VCHPTAWDLGKGDFRI MCTKVTMD+FLT
Sbjct  312  EKFFVSVGLPHMTQGFWGNSMLTDPGDDRKVVCHPTAWDLGKGDFRIKMCTKVTMDNFLT  371

Query  372  AHHEMGHIQYDMAYAAQPFLLRNGANEGFHEAVGEIMSLSAATPKHLKSIGLLSPDFQED  431
            AHHEMGHIQYDMAYA QPFLLRNGANEGFHEAVGEIMSLSAATPKHLKSIGLL  +F ED
Sbjct  372  AHHEMGHIQYDMAYATQPFLLRNGANEGFHEAVGEIMSLSAATPKHLKSIGLLPSNFHED  431

Query  432  NETEINFLLKQALTIVGTLPFTYMLEKWRWMVFKGEIPKDQWMKKWWEMKREIVGVVEPV  491
            NETEINFLLKQALTIVGTLPFTYMLEKWRWMVFKG+IPK++WM+KWWEMKREIVGVVEP+
Sbjct  432  NETEINFLLKQALTIVGTLPFTYMLEKWRWMVFKGDIPKEKWMEKWWEMKREIVGVVEPL  491

Query  492  PHDETYCDPASLFHVSNDYSFIRYYTRTLYQFQFQEALCQAAKHEGPLHKCDISNSTEAG  551
            PHDETYCDPA+LFHVSNDYSFIRYYTRT+YQFQFQEALCQAAKH+GPLHKCDISNSTEAG
Sbjct  492  PHDETYCDPAALFHVSNDYSFIRYYTRTIYQFQFQEALCQAAKHDGPLHKCDISNSTEAG  551

Query  552  QKLFNMLRLGKSEPWTLALENVVGAKNMNVRPLLNYFEPLFTWLKDQNKNSFVGWSTDWS  611
            QKL NMLRLGKSEPWTLALENVVGA+NM+VRPLLNYFEPL  WLK+QNKNSFVGW+TDWS
Sbjct  552  QKLLNMLRLGKSEPWTLALENVVGARNMDVRPLLNYFEPLSVWLKEQNKNSFVGWNTDWS  611

Query  612  PYADQSIKVRISLKSALGDKAYEWNDNEMYLFRSSVAYAMRQYFLKVKNQMILFGEEDVR  671
            PYADQSIKVRISLKSALG+ AYEWNDNEMYLFR++VAYAMR YF K K Q +LFG ED+R
Sbjct  612  PYADQSIKVRISLKSALGENAYEWNDNEMYLFRATVAYAMRVYFAKNKTQTVLFGVEDIR  671

Query  672  VANLKPRISFNFFVTAPKNVSDIIPRTEVEKAIRMSRSRINDAFRLNDNSLEFLGIQPTL  731
            V++LKPR+SFNFFVT+P+NVSDIIPR EVE+A+R SR RIND F L+DNSLEFLGI PTL
Sbjct  672  VSDLKPRVSFNFFVTSPQNVSDIIPRNEVEEAVRFSRGRINDVFGLDDNSLEFLGINPTL  731

Query  732  GPPNQPPVSIWLIVFGVVMGVIVVGIVILIFTGIRDRKKKNKARSGENPYASIDISKGEN  791
             PP QPPV+IWLI+FGVVMG++VVGIVILI TGIR RKK N+A+  ENPY S+DI KGE+
Sbjct  732  APPYQPPVTIWLIIFGVVMGIVVVGIVILIVTGIRARKKNNEAKREENPYDSVDIGKGES  791

Query  792  NPGFQNTDDVQTSF  805
            N GFQ+ DDVQTSF
Sbjct  792  NAGFQSNDDVQTSF  805
```

**angiotensin-converting enzyme 2 [Cricetulus griseus]**

Sequence ID: XP_027288607.1 Length: 805 Number of Matches: 1

See 1 more title(s)

#### Human vs Hamster

Range 1: 12 to 805 GenPept Graphics

▼ Next Match ▲ Pr

| Score | Expect | Method | Identities | Positives | Gaps |
|---|---|---|---|---|---|
| 1394 bits(3609) | 0.0 | Compositional matrix adjust. | 667/794(84%) | 726/794(91%) | 0/794(0%) |

```
Query  12   VAVTAAQSTIEEQAKTFLDKFNHEAEDLFYQSSLASWNYNTNITEENVQNMNNAGDKWSA  71
            VAVT AQS IEEQAKTFLDKFN EAEDL YQS+LASWNYNTNITEEN Q MN A  KWSA
Sbjct  12   VAVTTAQSIIEEQAKTFLDKFNQEAEDLSYQSALASWNYNTNITEENAQKMNEAAAKWSA  71

Query  72   FLKEQSTLAQMYPLQEIQNLTVKLQLQALQQNGSSVLSEDKSKRLNTILNTMSTIYSTGK  131
            F +EQS LA+ Y LQE+QNL +K QLQALQQ+GSS LS DK+K+LNTILNTMSTIYSTGK
Sbjct  72   FYEEQSKLAKNYSLQEVQNLIIKRQLQALQQSGSSALSADKNKQLNTILNTMSTIYSTGK  131

Query  132  VCNPDNPQECLLLEPGLNEIMANSLDYNERLWAWESWRSEVGKQLRPLYEEYVVLKNEMA  191
            VCNP NPQECLLLEPGL++IMA S DYNERLWAWE WR+EVGKQLRPLYEEYVVLKNEMA
Sbjct  132  VCNPKNPQECLLLEPGLDDIMATSTDYNERLWAWEGWRAEVGKQLRPLYEEYVVLKNEMA  191

Query  192  RANHYEDYGDYWRGDYEVNGVDGYDYSRGQLIEDVEHTFEEIKPLYEHLHAYVRAKLMNA  251
            RAN+Y+DYGDYWRGDYE  G DGY+Y+  QLIEDVE TF+EIKPLYE LHAYVR KLM+
Sbjct  192  RANNYKDYGDYWRGDYEAEGADGYNYNGNQLIEDVERTFKEIKPLYEQLHAYVRTKLMDT  251

Query  252  YPSYISPIGCLPAHLLGDMWGRFWTNLYSLTVPFGQKPNIDVTDAMVDQAWDAQRIFKEA  311
            YPS+ISP GCLPAHLLGDMWGRFWTNLY LTVPFGQKPNIDVTDAMV+Q WDA+RIFKEA
Sbjct  252  YPSFISPTGCLPAHLLGDMWGRFWTNLYPLTVPFGQKPNIDVTDAMVNQGWDAERIFKEA  311

Query  312  EKFFVSVGLPNMTQGFWENSMLTDPGNVQKAVCHPTAWDLGKGDFRILMCTKVTMDDFLT  371
            EKFFVSVGLP+MTQGFW NSMLTDPG+ +K VCHPTAWDLGKGDFRI MCTKVTMD+FLT
Sbjct  312  EKFFVSVGLPHMTQGFWGNSMLTDPGDDRKVVCHPTAWDLGKGDFRIKMCTKVTMDNFLT  371

Query  372  AHHEMGHIQYDMAYAAQPFLLRNGANEGFHEAVGEIMSLSAATPKHLKSIGLLSPDFQED  431
            AHHEMGHIQYDMAYA QPFLLRNGANEGFHEAVGEIMSLSAATPKHLKSIGLL  +F ED
Sbjct  372  AHHEMGHIQYDMAYATQPFLLRNGANEGFHEAVGEIMSLSAATPKHLKSIGLLPSNFHED  431

Query  432  NETEINFLLKQALTIVGTLPFTYMLEKWRWMVFKGEIPKDQWMKKWWEMKREIVGVVEPV  491
            NETEINFLLKQALTIVGTLPFTYMLEKWRWMVFKG+IPK++WM+KWWEMKREIVGVVEP+
Sbjct  432  NETEINFLLKQALTIVGTLPFTYMLEKWRWMVFKGDIPKEKWMEKWWEMKREIVGVVEPL  491

Query  492  PHDETYCDPASLFHVSNDYSFIRYYTRTLYQFQFQEALCQAAKHEGPLHKCDISNSTEAG  551
            PHDETYCDPA+LFHVSNDYSFIRYYTRT+YQFQFQEALCQAAKH+GPLHKCDISNSTEAG
Sbjct  492  PHDETYCDPAALFHVSNDYSFIRYYTRTIYQFQFQEALCQAAKHDGPLHKCDISNSTEAG  551

Query  552  QKLFNMLRLGKSEPWTLALENVVGAKNMNVRPLLNYFEPLFTWLKDQNKNSFVGWSTDWS  611
            QKL NMLRLGKSEPWTLALENVVGA+NM+VRPLLNYFEPL  WLK+QNKNSFVGW+TDWS
Sbjct  552  QKLLNMLRLGKSEPWTLALENVVGARNMDVRPLLNYFEPLSVWLKEQNKNSFVGWNTDWS  611

Query  612  PYADQSIKVRISLKSALGDKAYEWNDNEMYLFRSSVAYAMRQYFLKVKNQMILFGEEDVR  671
            PYADQSIKVRISLKSALG+ AYEWNDNEMYLFR++VAYAMR YF K K Q +LFG ED+R
Sbjct  612  PYADQSIKVRISLKSALGENAYEWNDNEMYLFRATVAYAMRVYFAKNKTQTVLFGVEDIR  671

Query  672  VANLKPRISFNFFVTAPKNVSDIIPRTEVEKAIRMSRSRINDAFRLNDNSLEFLGIQPTL  731
            V++LKPR+SFNFFVT+P+NVSDIIPR EVE+A+R SR RIND F L+DNSLEFLGI PTL
Sbjct  672  VSDLKPRVSFNFFVTSPQNVSDIIPRNEVEEAVRFSRGRINDVFGLDDNSLEFLGINPTL  731

Query  732  GPPNQPPVSIWLIVFGVVMGVIVVGIVILIFTGIRDRKKKNKARSGENPYASIDISKGEN  791
             PP QPPV+IWLI+FGVVMG++VVGIVILI TGIR RKK N+A+  ENPY S+DI KGE+
Sbjct  732  APPYQPPVTIWLIIFGVVMGIVVVGIVILIVTGIRARKKNNEAKREENPYDSVDIGKGES  791

Query  792  NPGFQNTDDVQTSF  805
            N GFQ+ DDVQTSF
Sbjct  792  NAGFQSNDDVQTSF  805
```

#### angiotensin-converting enzyme 2 [Mesocricetus auratus]

Sequence ID: XP_005074266.1 Length: 805 Number of Matches: 1

**Human vs Hamster**

Range 1: 12 to 805 GenPept Graphics ▼ Next Match ▲ Pr

| Score | Expect | Method | Identities | Positives | Gaps |
|---|---|---|---|---|---|
| 1398 bits(3619) | 0.0 | Compositional matrix adjust. | 669/794(84%) | 727/794(91%) | 0/794(0%) |

```
Query  12   VAVTAAQSTIEEQAKTFLDKFNHEAEDLFYQSSLASWNYNTNITEENVQNMNNAGDKWSA  71
            VAVT AQS IEEQAKTFLDKFN EAEDL YQS+LASWNYNTNITEEN Q MN A  KWSA
Sbjct  12   VAVTTAQSIIEEQAKTFLDKFNQEAEDLSYQSALASWNYNTNITEENAQKMNEAAAKWSA  71

Query  72   FLKEQSTLAQMYPLQEIQNLTVKLQLQALQQNGSSVLSEDKSKRLNTILNTMSTIYSTGK  131
            F +EQS LA+ Y LQE+QNLT+K QLQALQQ+GSS LS DK+K+LNTILNTMSTIYSTGK
Sbjct  72   FYEEQSKLAKNYSLQEVQNLTIKRQLQALQQSGSSALSADKNKQLNTILNTMSTIYSTGK  131

Query  132  VCNPDNPQECLLLEPGLNEIMANSLDYNERLWAWESWRSEVGKQLRPLYEEYVVLKNEMA  191
            VCNP NPQECLLLEPGL++IMA S DYNERLWAWE WR+EVGKQLRPLYEEYVVLKNEMA
Sbjct  132  VCNPKNPQECLLLEPGLDDIMATSTDYNERLWAWEGWRAEVGKQLRPLYEEYVVLKNEMA  191

Query  192  RANHYEDYGDYWRGDYEVNGVDGYDYSRGQLIEDVEHTFEEIKPLYEHLHAYVRAKLMNA  251
            RAN+YEDYGDYWRGDYE  G DGY+Y+  QLIEDVE TF+EIKPLYE LHAYVR KLMN
Sbjct  192  RANNYEDYGDYWRGDYEAEGADGYNYNGNQLIEDVERTFKEIKPLYEQLHAYVRTKLMNT  251

Query  252  YPSYISPIGCLPAHLLGDMWGRFWTNLYSLTVPFGQKPNIDVTDAMVDQAWDAQRIFKEA  311
            YPSYISP GCLPAHLLGDMWGRFWTNLY LTVPFGQKPNIDVTDAMV+Q W+A+RIFKEA
Sbjct  252  YPSYISPTGCLPAHLLGDMWGRFWTNLYPLTVPFGQKPNIDVTDAMVNQGWNAERIFKEA  311

Query  312  EKFFVSVGLPNMTQGFWENSMLTDPGNVQKAVCHPTAWDLGKGDFRILMCTKVTMDDFLT  371
            EKFFVSVGLP MTQGFWENSMLTDPG+ +K VCHPTAWDLGKGDFRI MCTKVTMD+FLT
Sbjct  312  EKFFVSVGLPYMTQGFWENSMLTDPGDDRKVVCHPTAWDLGKGDFRIKMCTKVTMDNFLT  371

Query  372  AHHEMGHIQYDMAYAAQPFLLRNGANEGFHEAVGEIMSLSAATPKHLKSIGLLSPDFQED  431
            AHHEMGHIQYDMAYA QPFLLRNGANEGFHEAVGEIMSLSAATP+HLKSIGLL  DFQED
Sbjct  372  AHHEMGHIQYDMAYATQPFLLRNGANEGFHEAVGEIMSLSAATPEHLKSIGLLPSDFQED  431

Query  432  NETEINFLLKQALTIVGTLPFTYMLEKWRWMVFKGEIPKDQWMKKWWEMKREIVGVVEPV  491
            NETEINFLLKQALTIVGTLPFTYMLEKWRWMVFKG+IPK+QWM+KWWEMKREIVGVVEP+
Sbjct  432  NETEINFLLKQALTIVGTLPFTYMLEKWRWMVFKGDIPKEQWMEKWWEMKREIVGVVEPL  491

Query  492  PHDETYCDPASLFHVSNDYSFIRYYTRTLYQFQFQEALCQAAKHEGPLHKCDISNSTEAG  551
            PHDETYCDPA+LFHVSNDYSFIRYYTRT+YQFQFQEALCQAAKH+GPLHKCDISNSTEAG
Sbjct  492  PHDETYCDPAALFHVSNDYSFIRYYTRTIYQFQFQEALCQAAKHDGPLHKCDISNSTEAG  551

Query  552  QKLFNMLRLGKSEPWTLALENVVGAKNMNVRPLLNYFEPLFTWLKDQNKNSFVGWSTDWS  611
            QKL NMLRLGKSEPWTLALENVVGA+NM+VRPLLNYFEPL  WLK+QNKNSFVGW+TDWS
Sbjct  552  QKLLNMLRLGKSEPWTLALENVVGARNMDVRPLLNYFEPLSVWLKEQNKNSFVGWNTDWS  611

Query  612  PYADQSIKVRISLKSALGDKAYEWNDNEMYLFRSSVAYAMRQYFLKVKNQMILFGEEDVR  671
            PYADQSIKVRISLKSALG+ AYEW+DNEMYLFR+SVAYAMR YF K K Q + FG ED+R
Sbjct  612  PYADQSIKVRISLKSALGENAYEWDDNEMYLFRASVAYAMRVYFAKNKTQTVPFGVEDIR  671

Query  672  VANLKPRISFNFFVTAPKNVSDIIPRTEVEKAIRMSRSRINDAFRLNDNSLEFLGIQPTL  731
            V++LKPR+SFNFFVT+P+NVSDIIPR EVE+A+R+SR RIND F L+DNSLEFLGI PTL
Sbjct  672  VSDLKPRVSFNFFVTSPQNVSDIIPRNEVEEAVRLSRGRINDVFGLDDNSLEFLGINPTL  731

Query  732  GPPNQPPVSIWLIVFGVVMGVIVVGIVILIFTGIRDRKKKNKARSGENPYASIDISKGEN  791
             PP QPPV+IWLI+FGVVMG++VVGI+ILIFTGI+ RKKKN+ +  ENPY S+DI KGE+
Sbjct  732  SPPYQPPVTIWLIIFGVVMGIVVVGIIILIFTGIKGRKKKNETKREENPYDSVDIGKGES  791

Query  792  NPGFQNTDDVQTSF  805
            N GF + DD QTSF
Sbjct  792  NAGFLSNDDAQTSF  805
```

#### Angiotensin-converting enzyme 2, partial [Neovison vison]

**Sequence ID:** CCP86723.1 **Length:** 471 **Number of Matches:** 1

### Human vs Mink

**Range 1: 1 to 471** GenPept Graphics ▼ Next Match ▲

| Score | Expect | Method | Identities | Positives | Gaps |
|---|---|---|---|---|---|
| 830 bits(2144) | 0.0 | Compositional matrix adjust. | 396/471(84%) | 441/471(93%) | 0/471(0%) |

```
Query  319  GLPNMTQGFWENSMLTDPGNVQKAVCHPTAWDLQKGDFRILMCTKVTMDDFLTAHHEMGH  378
            GLPNMT+GFW+NSMLT+PG+ +K VCHPTAWDLQK DFRI MCTKVTMDDFLTAHHEMGH
Sbjct  1    GLPNMTEGFWQNSMLTEPGDNRKVVCHPTAWDLQKHDFRIKMCTKVTMDDFLTAHHEMGH  60

Query  379  IQYDMAYAAQPFLLRNGANEGFHEAVGEIMSLSAATPKHLKSIGLLSPDFQEDNETEINF  438
            IQYDMAYAAQPFLLRNGANEGFHEAVGEIMSLSAATP HLK+IGLL PDF ED+ET+INF
Sbjct  61   IQYDMAYAAQPFLLRNGANEGFHEAVGEIMSLSAATPNHLKNIGLLPPDFSEDSETDINF  120

Query  439  LLKQALTIVGTLPFTYMLEKWRWMVFKGEIPKDQWMKKWWEMKREIVGVVEPVPHDETYC  498
            LLKQALTIVGTLPFTYMLEKWRWMVFKGEIPK+QWM+KWWEMKR+IVGVVEP+PHDETYC
Sbjct  121  LLKQALTIVGTLPFTYMLEKWRWMVFKGEIPKEQWMQKWWEMKRDIVGVVEPLPHDETYC  180

Query  499  DPASLFHVSNDYSFIRYYTRTLYQFQFQEALCQAAKHEGPLHKCDISNSTEAGQKLFNML  558
            DPA+LFHV+NDYSFIRYYTRT+YQFQFQEALCQ AKHEGPL+KCDISNS EAGQKL  ML
Sbjct  181  DPAALFHVANDYSFIRYYTRTIYQFQFQEALCQIAKHEGPLYKCDISNSREAGQKLHEML  240

Query  559  RLGKSEPWTLALENVVGAKNMNVRPLLNYFEPLFTWLKDQNKNSFVGWSTDWSPYADQSI  618
             LG+S+PWT ALE VVGAK M+VRPLLNYFEPLFTWLK+QN+NSFVGW+TDWSPYADQSI
Sbjct  241  SLGRSKPWTFALERVVGAKTMDVRPLLNYFEPLFTWLKEQNRNSFVGWNTDWSPYADQSI  300

Query  619  KVRISLKSALGDKAYEWNDNEMYLFRSSVAYAMRQYFLKVKNQMILFGEEDVRVANLKPR  678
            KVRISLKSALG+KAYEWNDNEMY F+SS+AYAMR+YF KVK Q I F ++DVRV++LKPR
Sbjct  301  KVRISLKSALGEKAYEWNDNEMYFFQSSIAYAMREYFSKVKKQTIPFVDKDVRVSDLKPR  360

Query  679  ISFNFFVTAPKNVSDIIPRTEVEKAIRMSRSRINDAFRLNDNSLEFLGIQPTLGPPNQPP  738
            ISFNF VT+P+N+SDIIPR +VE+AIR SR RINDAFRL+DNSLEFLGIQPTL PP QPP
Sbjct  361  ISFNFIVTSPENMSDIIPRADVEEAIRKSRGRINDAFRLDDNSLEFLGIQPTLEPPYQPP  420

Query  739  VSIWLIVFGVVMGVIVVGIVILIFTGIRDRKKKNKARSGENPYASIDISKG  789
            V+IWLIVFGVVMGV+VVGI +LIF+GIR+R+K N+ARS ENPYAS+D+SKG
Sbjct  421  VTIWLIVFGVVMGVVVVGIFLLIFSGIRNRRKNNQARSEENPYASVDLSKG  471
```

https://www.ncbi.nlm.nih.gov/nuccore/HAAF01014901.1

https://blast.ncbi.nlm.nih.gov/Blast.cgi#alnHdr_1064617951

**Sequence ID:** XP_014062928.1 **Length:** 686 **Number of Matches:** 1

### Human vs Atlantic salmon

**Range 1: 1 to 686** GenPept Graphics ▼ Next Match ▲ Pre

| Score | Expect | Method | Identities | Positives | Gaps |
|---|---|---|---|---|---|
| 877 bits(2267) | 0.0 | Compositional matrix adjust. | 418/693(60%) | 513/693(74%) | 14/693(2%) |

```
Query  120  LNTMSTIYSTGKVCNPDNPQECLLLEPGLNEIMAN-SLDYNERLWAWESWRSEVGKQLRP  178
            +N MS+IYSTG VC  ++P +C  LEPGL  +MAN   DY ERL  WE WR EVGK++RP
Sbjct  1    MNKMSSIYSTGTVCKREDPFDCQTLEPGLESVMANMDSDYYERLHVWEGWRVEVGKKMRP  60

Query  179  LYEEYVVLKNEMARANHYEDYGDYWRGDYEVNGVDGYDYSRGQLIEDVEHTFEEIKPLYE  238
            LYE+YV LKNE A+ N YEDYGDYWR +YE       Y+Y+RGQL+ DV H ++EI PLY+
Sbjct  61   LYEDYVDLKNEAAKLNGYEDYGDYWRSNYETIDDSPYNYARGQLMTDVRHIYKEILPLYK  120

Query  239  HLHAYVRAKLMNAYPSYISPIGCLPAHLLGDMWGRFWTNLYSLTVPFGQKPNIDVTDAMV  298
             LHAYVR+KL   +P +I P G LPAHLLGDMWGRFWT LY ++ PF +K +IDVTDAM+
Sbjct  121  ELHAYVRSKLQAKHPEHIHPEGGLPAHLLGDMWGRFWTGLYPISTPFPEKTDIDVTDAMI  180

Query  299  DQAWDAQRIFKEAEKFFVSVGLPNMTQGFWENSMLTDPGNVQKAVCHPTAWDLG-KGDFR  357
             Q W   R+F+EAEKFF+SVGL   M   FW++SML  P + +K VCHPTAWD+G + DFR
Sbjct  181  AQKWPKDRLFQEAEKFFMSVGLYKMFDNFWKDSMLEKPTDGRKVVCHPTAWDMGNREDFR  240

Query  358  ILMCTKVTMDDFLTAHHEMGHIQYDMAYAAQPFLLRNGANEGFHEAVGEIMSLSAATPKH  417
            I MCT+V MD FLTAHHEMGH QY MAY    +LLR+GANEGFHEAVGEIMSLSAATPKH
Sbjct  241  IKMCTEVNMDHFLTAHHEMGHNQYQMAYRNLSYLLRDGANEGFHEAVGEIMSLSAATPKH  300

Query  418  LKSIGLLSPDFQEDNETEINFLLKQALTIVGTLPFTYMLEKWRWMVFKGEIPKDQWMKKW  477
            LK++GLL  DF ED ETEINFL+KQALTIV TLPFTYMLE+WRW VF G IPKDQWM++W
Sbjct  301  LKALGLLPDDFVEDKETEINFLMKQALTIVATLPFTYMLEEWRWQVFLGTIPKDQWMQRW  360

Query  478  WEMKREIVGVVEPVPHDETYCDPASLFHVSNDYSFIRYYTRTLYQFQFQEALCQAAKHEG  537
            WEMKR++VGVVEP+P DETYCDP +LFHVS DYSFIRY+TRT+YQFQFQ+ALC+AA H G
Sbjct  361  WEMKRDMVGVVEPLPRDETYCDPPALFHVSGDYSFIRYFTRTIYQFQFQKALCEAAGHSG  420

Query  538  PLHKCDISNSTEAGQKLFNMLRLGKSEPWTLALENVVGAKNMNVRPLLNYFEPLFTWLKD  597
            PL KCDI+NST AG KL  ML  G+S+ WT ALE + G   M+  PLL+YF+ L  WL +
Sbjct  421  PLFKCDITNSTAAGDKLRTMLEFGRSKSWTRALETISGHAKMDSAPLLDYFKDLHVWLIE  480

Query  598  QNK--NSFVGWSTDWSPYADQSIKVRISLKSALGDKAYEWNDNEMYLFRSSVAYAMRQYF  655
            +N+  N   GW     P+++ + KVR+SLK+A+GDKAY WN NEMYLF++++AYAMRQY+
Sbjct  481  ENRKNNRKPGWRAAEDPFSENAYKVRLSLKAAMGDKAYIWNANEMYLFKANMAYAMRQYY  540

Query  656  LKVKNQMILFGEEDVRVANLKPRISFNFFVTAPKNVSDIIPRTEVEKAIRMSRSRINDAF  715
            L+V    +LF  E++         RISF F VT P N + +IP+ EVE AIR+SR RINDAF
Sbjct  541  LEVNKTEVLFTTENIHTYKETARISFYFVVTDPANPAVVIPKAEVEAAIRLSRGRINDAF  600

Query  716  RLNDNSLEFLGIQPTLGPPNQPPVSIWLIVFGVVMGVIVVGIVILIFTGIRDR---KKKN  772
            +L+D +LEF G+  TL PP + PV++WL+VFGVVMG++V     LI +G RDR
Sbjct  601  KLDDKTLEFEGLLATLAPPVEQPVTVWLVVFGVVMGLVVCMGCYLIISGFRDRKKKCAAK  660

Query  773  KARSGENPYASIDISKGENNPGFQNTDDVQTSF  805
               + ENPY       G  N  F+  +D QT F
Sbjct  661  AKENAENPY-------GVTNKTFEREEDEQTGF  686
```

**Human angiotensin-converting enzyme 2 precursor [Homo sapiens]**
NCBI Reference Sequence: NP_001358344.1
**Dog#1 angiotensin-converting enzyme 2 precursor [Canis lupus familiaris]**
NCBI Reference Sequence: NP_001158732.1
**Dog#2 angiotensin-converting enzyme 2 isoform X1 [Canis lupus familiaris]**
NCBI Reference Sequence: XP_005641049.1
**Dog#3 angiotensin-converting enzyme 2 isoform X1 [Canis lupus familiaris]**
NCBI Reference Sequence: XP_013966804.1
**Dog#4 angiotensin-converting enzyme 2 isoform X2 [Canis lupus familiaris]**
NCBI Reference Sequence: XP_022271214.1
**Cat#1 angiotensin-converting enzyme 2 isoform X1 [Felis catus]**
NCBI Reference Sequence: XP_023104564.1
**Cat#2 angiotensin-converting enzyme 2 precursor [Felis catus]**
NCBI Reference Sequence: NP_001034545.1
**Bat#1 PREDICTED: angiotensin-converting enzyme 2 isoform X1 [Myotis brandtii]**
NCBI Reference Sequence: XP_014399780.1
**Bat#2 PREDICTED: angiotensin-converting enzyme 2 isoform X1 [Myotis brandtii]**
NCBI Reference Sequence: XP_014399781.1
**Bat#3 PREDICTED: angiotensin-converting enzyme 2 isoform X2 [Myotis brandtii]**
NCBI Reference Sequence: XP_014399782.1
**Bat#4 PREDICTED: angiotensin-converting enzyme 2 isoform X3 [Myotis brandtii]**
NCBI Reference Sequence: XP_014399783.1
**Bat#5 angiotensin-converting enzyme 2 isoform X1 [Desmodus rotundus]**
NCBI Reference Sequence: XP_024425698.1
**Bat#6 angiotensin-converting enzyme 2 isoform X2 [Desmodus rotundus]**
NCBI Reference Sequence: XP_024425699.1

**Bat#7 angiotensin-converting enzyme 2 isoform X1 [Eptesicus fuscus]**
NCBI Reference Sequence: XP_008153150.1
**Bat#8 angiotensin-converting enzyme 2 isoform X2 [Eptesicus fuscus]**
NCBI Reference Sequence: XP_027986092.1
**Bat#9 angiotensin-converting enzyme 2 isoform X1 [Myotis lucifugus]**
NCBI Reference Sequence: XP_023609437.1
**Bat#10 angiotensin-converting enzyme 2 isoform X1 [Myotis lucifugus]**
NCBI Reference Sequence: XP_023609438.1
**Bat#11 angiotensin-converting enzyme 2 isoform X2 [Myotis lucifugus]**
NCBI Reference Sequence: XP_023609439.1
**Bat#12 angiotensin-converting enzyme 2 [Phyllostomus discolor]**
NCBI Reference Sequence: XP_028378317.1
**Bat#13 PREDICTED: angiotensin-converting enzyme 2 [Hipposideros armiger]**
NCBI Reference Sequence: XP_019522936.1
**Bat#14 PREDICTED: angiotensin-converting enzyme 2 [Hipposideros armiger]**
NCBI Reference Sequence: XP_019522943.1
**Bat#15 PREDICTED: angiotensin-converting enzyme 2 [Hipposideros armiger]**
NCBI Reference Sequence: XP_019522954.1
**Pangolin#1 PREDICTED: angiotensin-converting enzyme 2 [Manis javanica]**
NCBI Reference Sequence: XP_017505746.1
**Pangolin#2 PREDICTED: angiotensin-converting enzyme 2 [Manis javanica]**
NCBI Reference Sequence: XP_017505752.1
**Snake#1 LOW QUALITY PROTEIN: angiotensin-converting enzyme 2 [Notechis scutatus]**
NCBI Reference Sequence: XP_026530754.1
**Snake#2 angiotensin-converting enzyme 2 [Thamnophis elegans]**
NCBI Reference Sequence: XP_032082934.1

**Human angiotensin-converting enzyme 2 precursor [Homo sapiens]**
NCBI Reference Sequence: NP_001358344.1
**Mouse angiotensin-converting enzyme 2 precursor [Mus musculus]**
NCBI Reference Sequence: NP_081562.2
**Mouse angiotensin-converting enzyme 2 precursor [Mus musculus]**
NCBI Reference Sequence: NP_001123985.1
**Mouse angiotensin-converting enzyme 2 [Peromyscus leucopus]**
NCBI Reference Sequence: XP_028743609.1
**Mouse angiotensin-converting enzyme 2 [Mus caroli]**
NCBI Reference Sequence: XP_021009138.1
**Mouse angiotensin-converting enzyme 2 [Mus pahari]**
NCBI Reference Sequence: XP_021043935.1
**Mouse angiotensin-converting enzyme 2 [Mastomys coucha]**
NCBI Reference Sequence: XP_031226742.1
**Mouse angiotensin-converting enzyme 2 [Peromyscus maniculatus bairdii]**
NCBI Reference Sequence: XP_006973269.1
**Mouse angiotensin-converting enzyme 2 [Condylura cristata]**
NCBI Reference Sequence: XP_012585871.1
**Rat angiotensin-converting enzyme 2 precursor [Rattus norvegicus]**
NCBI Reference Sequence: NP_001012006.1
**Rat angiotensin-converting enzyme 2 [Heterocephalus glaber]**
NCBI Reference Sequence: XP_004866157.1
**Rat angiotensin-converting enzyme 2 isoform X1 [Dipodomys ordii]**
NCBI Reference Sequence: XP_012887572.1
**Rat angiotensin-converting enzyme 2 isoform X2 [Dipodomys ordii]**
NCBI Reference Sequence: XP_012887573.1

**Rat angiotensin-converting enzyme 2 [Rattus rattus]**
NCBI Reference Sequence: XP_032746145.1
**Rat angiotensin-converting enzyme 2 [Fukomys damarensis]**
NCBI Reference Sequence: XP_010643477.1
**Rat angiotensin-converting enzyme 2 [Nannospalax galili]**
NCBI Reference Sequence: XP_008839098.1
**Hamster angiotensin-converting enzyme 2 [Cricetulus griseus]**
NCBI Reference Sequence: XP_003503283.1
**Hamster angiotensin-converting enzyme 2 [Cricetulus griseus]**
NCBI Reference Sequence: XP_027288607.1
**Hamster angiotensin-converting enzyme 2 [Mesocricetus auratus]**
NCBI Reference Sequence: XP_005074266.1
**Mink angiotensin-converting enzyme 2, partial [Neovison vison]**
NCBI Reference Sequence: CCP86723.1
**Salmon angiotensin-converting enzyme 2 [Salmo salar]**
NCBI Reference Sequence: XP_014062928.1

*Homo sapiens*
human

ACE2
angiotensin I converting enzyme 2

805

| RefSeq transcripts (5) | RefSeq proteins (5) | Architecture | aa |
|---|---|---|---|
| NM_001371415.1 | NP_001358344.1 | | 805 |
| NM_021804.3 | NP_068576.1 | | 805 |
| XM_011545549.2 | XP_011543851.1 | | 786 |
| XM_011545551.3 | XP_011543853.1 | | 772 |
| XM_011545552.2 | XP_011543854.1 | | 694 |

Genome Browser

InterPro

https://www.ncbi.nlm.nih.gov/gene/59272/ortholog/?scope=7776

*Canis lupus familiaris*
dog

ACE2
angiotensin I converting enzyme 2

804

| RefSeq transcripts (4) | RefSeq proteins (4) | Architecture | aa |
|---|---|---|---|
| NM_001165260.1 | NP_001158732.1 | | 804 |
| XM_005640992.2 | XP_005641049.1 | | 804 |
| XM_014111329.2 | XP_013966804.1 | | 804 |
| XM_022415506.1 | XP_022271214.1 | | 683 |

Genome Browser  InterPro

https://www.ncbi.nlm.nih.gov/gene/59272/ortholog/?scope=7776

*Felis catus*
domestic cat

ACE2
angiotensin I converting enzyme 2

807

| RefSeq transcripts (2) | RefSeq proteins (2) | Architecture | aa |
|---|---|---|---|
| XM_023248796.1 | XP_023104564.1 | | 807 |
| NM_001039456.1 | NP_001034545.1 | | 805 |

Genome Browser

InterPro

https://www.ncbi.nlm.nih.gov/gene/59272/ortholog/?scope=7776

*Panthera tigris altaica* ACE2 797

Amur tiger

angiotensin I converting enzyme 2

| RefSeq transcripts (1) | RefSeq proteins (1) | Architecture | aa |
|---|---|---|---|
| XM_007090080.2 | XP_007090142.1 | 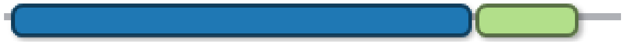 | 797 |

Genome Browser

https://www.ncbi.nlm.nih.gov/gene/59272/ortholog/?scope=7776

*Myotis brandtii* ACE2 819

Brandts bat

angiotensin I converting enzyme 2

| RefSeq transcripts (4) | RefSeq proteins (4) | Architecture | aa |
|---|---|---|---|
| XM_014544294.1 | XP_014399780.1 | | 819 |
| XM_014544295.1 | XP_014399781.1 | | 819 |
| XM_014544296.1 | XP_014399782.1 | | 799 |
| XM_014544297.1 | XP_014399783.1 | | 754 |

Genome Browser InterPro

https://www.ncbi.nlm.nih.gov/gene/59272/ortholog/?scope=7776

*Desmodus rotundus*
common vampire bat

ACE2
angiotensin I converting enzyme 2

804

| RefSeq transcripts (2) | RefSeq proteins (2) | Architecture | aa |
|---|---|---|---|
| XM_024569930.1 | XP_024425698.1 | | 804 |
| XM_024569931.1 | XP_024425699.1 | | 737 |

Genome Browser

https://www.ncbi.nlm.nih.gov/gene/59272/ortholog/?scope=7776

*Eptesicus fuscus*
big brown bat

ACE2
angiotensin I converting enzyme 2

811

| RefSeq transcripts (2) | RefSeq proteins (2) | Architecture | aa |
|---|---|---|---|
| XM_008154928.2 | XP_008153150.1 | | 811 |
| XM_028130291.1 | XP_027986092.1 | | 799 |

Genome Browser

https://www.ncbi.nlm.nih.gov/gene/59272/ortholog/?scope=7776

*Myotis lucifugus*
little brown bat

ACE2
angiotensin I converting enzyme 2

819

| RefSeq transcripts (3) | RefSeq proteins (3) | Architecture | aa |
|---|---|---|---|
| XM_023753669.1 | XP_023609437.1 | | 819 |
| XM_023753670.1 | XP_023609438.1 | | 819 |
| XM_023753671.1 | XP_023609439.1 | | 799 |

Genome Browser

https://www.ncbi.nlm.nih.gov/gene/59272/ortholog/?scope=7776

*Phyllostomus discolor*
pale spear-nosed bat

ACE2
angiotensin I converting enzyme 2

804

| RefSeq transcripts (1) | RefSeq proteins (1) | Architecture | aa |
|---|---|---|---|
| XM_028522516.1 | XP_028378317.1 | 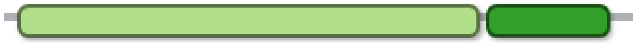 | 804 |

Genome Browser

https://www.ncbi.nlm.nih.gov/gene/59272/ortholog/?scope=7776

*Hipposideros armiger*
great roundleaf bat

ACE2
angiotensin I converting enzyme 2

806

| RefSeq transcripts (3) | RefSeq proteins (3) | Architecture | aa |
|---|---|---|---|
| XM_019667391.1 | XP_019522936.1 | | 806 |
| XM_019667398.1 | XP_019522943.1 | | 806 |
| XM_019667409.1 | XP_019522954.1 | | 806 |

Genome Browser

https://www.ncbi.nlm.nih.gov/gene/59272/ortholog/?scope=7776

*Manis javanica*
Malayan pangolin

ACE2
angiotensin I converting enzyme 2

805

| RefSeq transcripts (2) | RefSeq proteins (2) | Architecture | aa |
|---|---|---|---|
| XM_017650257.1 | XP_017505746.1 | | 805 |
| XM_017650263.1 | XP_017505752.1 | | 805 |

Genome Browser

https://www.ncbi.nlm.nih.gov/gene/59272/ortholog/?scope=7776

*Notechis scutatus* mainland tiger snake

ACE2 angiotensin I converting enzyme 2 — 828

| RefSeq transcripts (1) | RefSeq proteins (1) | Architecture | aa |
|---|---|---|---|
| XM_026674969.1 | XP_026530754.1 | | 828 |

Genome Browser

https://www.ncbi.nlm.nih.gov/gene/59272/ortholog/?scope=7776

*Thamnophis elegans* Western terrestrial garter snake | ACE2 angiotensin I converting enzyme 2 | 828

| RefSeq transcripts (1) | RefSeq proteins (1) | Architecture | aa |
|---|---|---|---|
| XM_032227043.1 | XP_032082934.1 | | 828 |

Genome Browser

https://www.ncbi.nlm.nih.gov/gene/59272/ortholog/?scope=7776

*Mus musculus*
house mouse

Ace2
angiotensin I converting enzyme (peptidyl-dipeptidase A) 2

805

| RefSeq transcripts (2) | RefSeq proteins (2) | Architecture | aa |
| --- | --- | --- | --- |
| NM_027286.4 | NP_081562.2 | 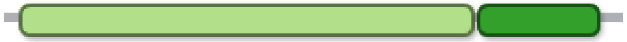 | 805 |
| NM_001130513.1 | NP_001123985.1 | 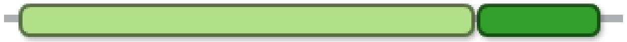 | 805 |

Genome Browser

InterPro

☐ *Peromyscus leucopus* — white-footed mouse

Ace2 — angiotensin I converting enzyme 2 — 805

| RefSeq transcripts (1) | RefSeq proteins (1) | Architecture | aa |
|---|---|---|---|
| XM_028887776.1 | XP_028743609.1 | 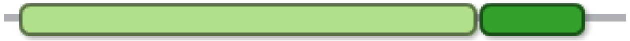 | 805 |

Genome Browser

☐ *Mus caroli* — Ryukyu mouse

Ace2 — angiotensin I converting enzyme 2 — 805

| RefSeq transcripts (1) | RefSeq proteins (1) | Architecture | aa |
|---|---|---|---|
| XM_021153479.2 | XP_021009138.1 | 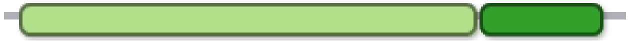 | 805 |

Genome Browser

☐ *Mus pahari*
shrew mouse

Ace2
angiotensin I converting enzyme 2

805

| RefSeq transcripts (1) | RefSeq proteins (1) | Architecture | aa |
|---|---|---|---|
| XM_021188276.2 | XP_021043935.1 | 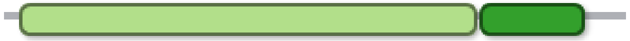 | 805 |

**Genome Browser**

☐ *Mastomys coucha*
southern multimammate mouse

Ace2
angiotensin I converting enzyme 2

806

| RefSeq transcripts (1) | RefSeq proteins (1) | Architecture | aa |
|---|---|---|---|
| XM_031370882.1 | XP_031226742.1 | | 806 |

**Genome Browser**

*Condylura cristata*
star-nosed mole

ACE2
angiotensin I converting enzyme 2

800

| RefSeq transcripts (1) | RefSeq proteins (1) | Architecture | aa |
|---|---|---|---|
| XM_012730417.1 | XP_012585871.1 | 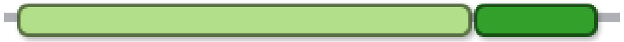 | 800 |

Genome Browser

*Peromyscus maniculatus bairdii*
prairie deer mouse

Ace2
angiotensin I converting enzyme 2

805

| RefSeq transcripts (1) | RefSeq proteins (1) | Architecture | aa |
|---|---|---|---|
| XM_006973207.2 | XP_006973269.1 | 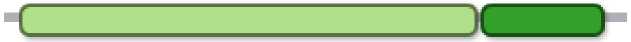 | 805 |

Genome Browser

*Rattus norvegicus* Norway rat — Ace2, angiotensin I converting enzyme 2 — 805

| RefSeq transcripts (1) | RefSeq proteins (1) | Architecture | aa |
|---|---|---|---|
| NM_001012006.1 | NP_001012006.1 | 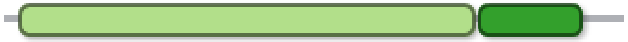 | 805 |

Genome Browser InterPro

*Heterocephalus glaber* naked mole-rat — Ace2, angiotensin I converting enzyme 2 — 805

| RefSeq transcripts (1) | RefSeq proteins (1) | Architecture | aa |
|---|---|---|---|
| XM_004866100.3 | XP_004866157.1 | 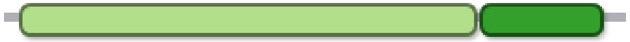 | 805 |

Genome Browser InterPro

☐ *Dipodomys ordii*
Ords kangaroo rat

Ace2
angiotensin I converting enzyme 2

805

| RefSeq transcripts (2) | RefSeq proteins (2) | Architecture | aa |
|---|---|---|---|
| XM_013032118.1 | XP_012887572.1 | 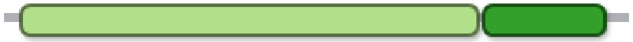 | 805 |
| XM_013032119.1 | XP_012887573.1 | 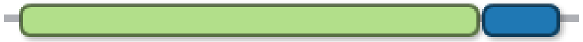 | 740 |

**Genome Browser**  InterPro

☐ *Rattus rattus*
black rat

Ace2
angiotensin I converting enzyme 2

793

| **RefSeq transcripts (1)** | **RefSeq proteins (1)** | **Architecture** | **aa** |
|---|---|---|---|
| XM_032890254.1 | XP_032746145.1 | | 793 |

**Genome Browser**

☐ *Fukomys damarensis*
Damara mole-rat

Ace2
angiotensin I converting enzyme 2

805

| **RefSeq transcripts (1)** | **RefSeq proteins (1)** | **Architecture** | **aa** |
|---|---|---|---|
| XM_010645175.2 | XP_010643477.1 | 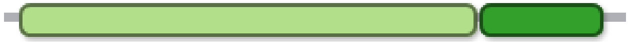 | 805 |

**Genome Browser**

*Nannospalax galili* Ace2 804
Upper Galilee mountains blind mole rat
angiotensin I converting enzyme 2

| RefSeq transcripts (1) | RefSeq proteins (1) | Architecture | aa |
|---|---|---|---|
| XM_008840876.2 | XP_008839098.1 | 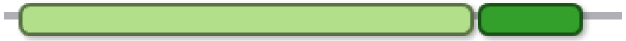 | 804 |

Genome Browser

*Cricetulus griseus* | Ace2 | 805

Chinese hamster

angiotensin I converting enzyme 2

| RefSeq transcripts (2) | RefSeq proteins (2) | Architecture | aa |
|---|---|---|---|
| XM_003503235.4 | XP_003503283.1 | 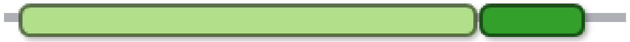 | 805 |
| XM_027432806.1 | XP_027288607.1 | 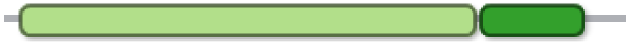 | 805 |

Genome Browser

*Mesocricetus auratus*
golden hamster

Ace2
angiotensin I converting enzyme 2

805

| RefSeq transcripts (1) | RefSeq proteins (1) | Architecture | aa |
|---|---|---|---|
| XM_005074209.2 | XP_005074266.1 | 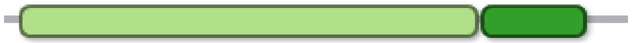 | 805 |

Genome Browser

InterPro
